# Metabolic flexibility allows generalist bacteria to become dominant in a frequently disturbed ecosystem

**DOI:** 10.1101/2020.02.12.945220

**Authors:** Ya-Jou Chen, Pok Man Leung, Sean K. Bay, Philip Hugenholtz, Adam J. Kessler, Guy Shelley, David W. Waite, Perran L. M. Cook, Chris Greening

**Author notes:** Correspondence can be addressed to: A/Prof Chris Greening, Prof Perran Cook.

## Abstract

Ecological theory suggests that habitat disturbance differentially influences distributions of generalist and specialist species. While well-established for macroorganisms, this theory has rarely been explored for microorganisms. Here we tested these principles in permeable (sandy) sediments, ecosystems with much spatiotemporal variation in resource availability and other conditions. Microbial community composition and function was profiled in intertidal and subtidal sediments using 16S amplicon sequencing and metagenomics, yielding 135 metagenome-assembled genomes. Microbial abundance and composition significantly differed with sediment depth and, to a lesser extent, sampling date. Several generalist taxa were highly abundant and prevalent in all samples, including within orders Woeseiales and Flavobacteriales; genome reconstructions indicate these facultatively anaerobic taxa are highly metabolically flexible and adapt to fluctuations in resource availability by using different electron donors and acceptors. In contrast, obligately anaerobic taxa such as sulfate reducers (Desulfobacterales, Desulfobulbales) and proposed candidate phylum MBNT15 were less abundant overall and only thrived in more stable deeper sediments. We substantiated these findings by measuring three metabolic processes in these sediments; whereas the generalist-associated processes of sulfide oxidation and hydrogenogenic fermentation occurred rapidly at all depths, the specialist-associated process of sulfate reduction was restricted to deeper sediments. In addition, a manipulative experiment confirmed generalists outcompete specialist taxa during simulated habitat disturbance. Altogether, these findings suggest that metabolically flexible taxa become dominant in these highly dynamic environments, whereas metabolic specialism restricts bacteria to narrower niches. Thus, an ecological theory describing distribution patterns for macroorganisms likely extends to microorganisms. Such findings have broad ecological and biogeochemical ramifications.

## Introduction

In macroecology, species are broadly classified as habitat generalists and specialists depending on their niche breadth^1,2^. Both deterministic and stochastic factors control the differential distributions of such species and in turn the maintenance of diversity^3^ With respect to deterministic factors, a pervasive ecological theory is that generalists and specialists differ in performance traits, for example resource utilisation; it is thought habitat generalists are more versatile but less efficient than habitat specialists, whereas specialists perform fewer activities more effectively. By extension, it can be predicted that specialists will outperform generalists in their optimal habitats, whereas generalists will be favoured in environments with high spatial and temporal heterogeneity^1,4^. In turn, there is evidence that both natural and anthropogenic habitat disturbance favours generalists and promotes homogenisation of community composition^5–7^. Other factors, notably dispersal traits and life history strategies, also influence distribution patterns^3,8^. While these tenets are well-established for animals and plants, few studies have extended them to microbial communities^9–11^.

The key ecological processes governing macroorganism community assembly are thought to extend to microorganisms. However, environmental filtering tends to predominate over neutral factors such as dispersal limitation^12–14^ In turn, these processes lead to an uneven prevalence of microbial taxa across ecosystems; most community members have low to intermediate occupancy (habitat specialists), but a small proportion of taxa tend to be highly prevalent and often abundant across space and time (habitat generalists)^15,16^. The performance traits that differentiate microbial generalists and specialists have been scarcely explored. It is probable that, like macroorganisms, a key factor that governs distribution patterns is the capacity and efficiency of resource utilisation. In this regard, an important feature that distinguishes microorganisms is their metabolic versatility^17^; whereas plants and animals are respectively restricted to photoautotrophic and chemoheterotrophic growth, many microorganisms can use multiple energy sources, carbon sources, and oxidants either simultaneously or alternately^11^. Likewise, the capacity for microorganisms to transition between active and dormant states contributes to the maintenance of diversity^18,19^. It is being increasingly realised that such flexibility in resource usage contributes to the dominance of major taxa in various environments^20–24^

Permeable (sandy) sediments are ideal sites to explore the concepts of microbial generalism and specialism. These ecosystems, spanning at least half the continental shelf, are important regulators of oceanic biogeochemical cycling and primary production^25–27^ Their mixing layers are continuously disrupted as a result of porewater advection, tidal flows, and other factors^28–30^ As a result, resident microorganisms experience large spatiotemporal variations in the availability of light, oxygen, and other resources^25,31^. In contrast, the microbial communities in the deeper sediment layers are infrequently disturbed and are generally exposed to dark anoxic conditions^32^. Despite these pressures, these sediments are known to harbour abundant, diverse, and active microbial communities^33–37^ Previous studies have indicated that there is a rapid community turnover across depth and season in Wadden Sea sediments^33^ However, some lineages such as the Woeseiales appear to be abundant and prevalent residents of all permeable sediments sampled worldwide ^21,38,39^. The functional basis for their dominance is unclear. We have recently published evidence that metabolic flexibility, including the ability of bacteria to shift from aerobic respiration to hydrogenogenic fermentation in response to oxic-anoxic transitions, is an important factor controlling the ecology and biogeochemistry of the communities in the mixing layer^38,40^.

In this study, we investigated the spatiotemporal distributions and metabolic traits of habitat generalists and specialists in permeable sediments from Middle Park Beach, Port Philip Bay, Australia. Given the above considerations, we hypothesised that the mixing and deep layers of permeable sediments would select for different microbial traits. The mixing layer, reflecting its spatiotemporal variability, would select for microbial generalists with broad metabolic capabilities. In contrast, the less frequently disturbed deep layer would select for relative specialists with restricted but efficient anaerobic lifestyles. To test this, we used high-resolution community profiling to determine the spatiotemporal distribution of bacterial and archaeal communities in shallow, intermediate, and deep sediments. In combination, we used genome-resolved metagenomics, biogeochemical assays, and perturbation experiments to determine the metabolic capabilities of the most dominant habitat generalists and specialists.

## Results and Discussion

### Habitat generalists dominate permeable sediments, but co-exist with depth-restricted specialists

We used the 16S rRNA gene as a marker to profile the diversity, abundance, and composition of the bacterial and archaeal communities in permeable sediments. 48 sand samples were profiled that were collected from intertidal and subtidal sediments at three different depths (shallow: 0-3 cm, intermediate: 14-17 cm, deep: 27-30 cm) and across eight different dates over the course of a year **(Table S1)**. Alpha diversity indices indicated that the sands support the co-existence of diverse microorganisms; Shannon index was high across the samples (6.79 ± 0.30) and no significant differences were observed across depth, zone, or time **(Fig. 1a, Fig. S1 & S2)**. However, a significant decrease in community abundance with depth (inferred from 16S rRNA gene copy number by qPCR) across the samples **(Fig. 1b)**. This correlated with the transition from the mixing zone (above 20 cm) to the sustained aphotic anoxic zone (below 20 cm), as indicated by a sharp decrease in chlorophyll *a* abundance **(Fig. 1c)** and an increase in acid-volatile sulfide concentrations (from below detection limits to 0.16 μmol g^-1^).

**Figure 1.**
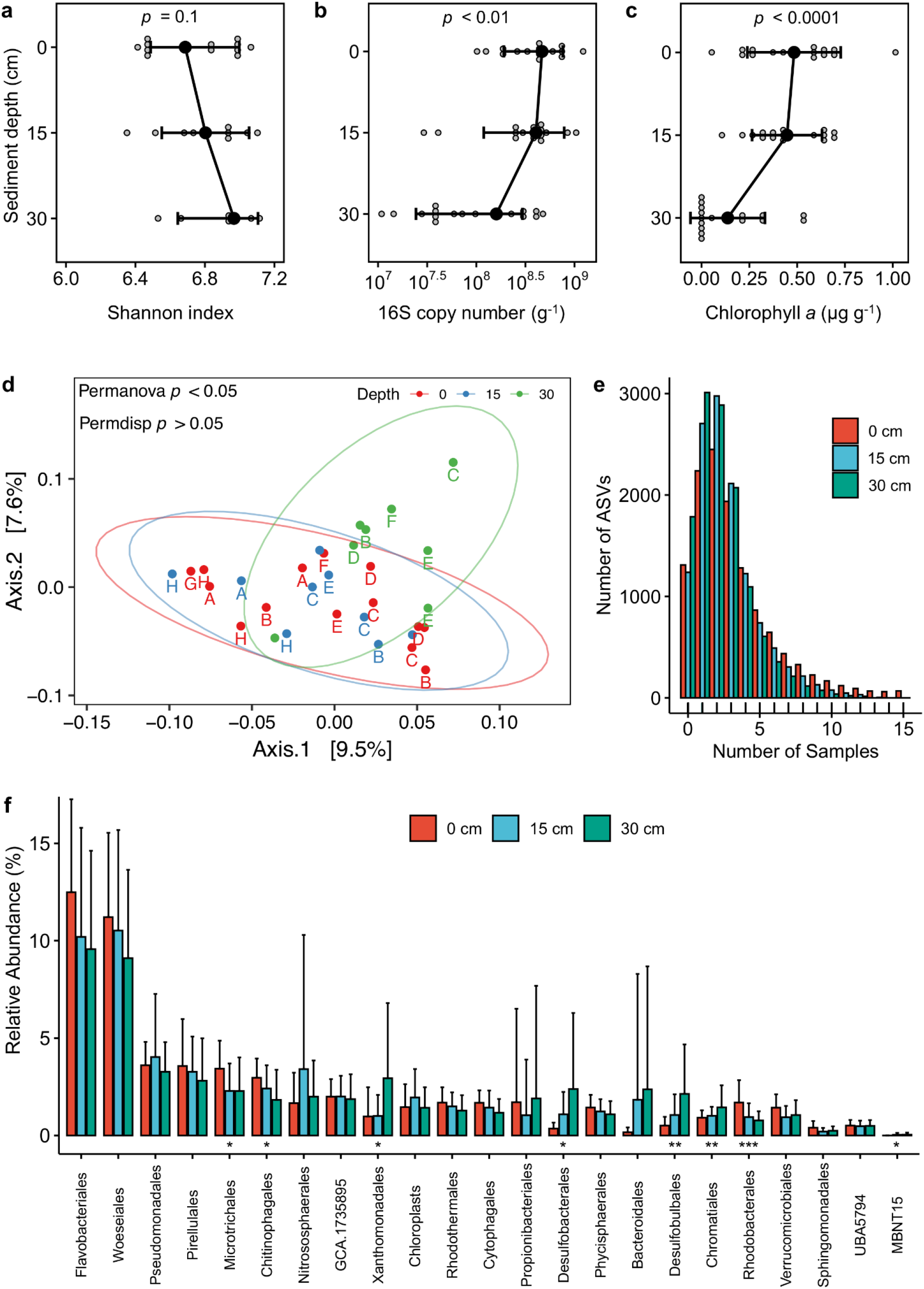
Diversity, abundance, and composition of bacterial and archaeal communities in permeable sediments. The figures are based on the results of 16S rRNA gene sequencing for 48 samples covering two tidal zones (intertidal, subtidal), three sediment depths (0-3 cm, 13-17 cm, 27-30 cm), and eight sampling times (between Oct 2016 to Oct 2017). Variations in **(a)** Shannon index (alpha diversity), **(b)** 16S copy number, and **(c)** chlorophyll *a* concentration are shown with depth; error bars show standard deviations of the mean and significance was tested using one-way ANOVAs. **(d)** Principal coordinates analysis (PCoA) plot visualizing pairwise dissimilatory (beta diversity) of communities using weighted Unifrac. **(e)** Occupancy frequency distribution of the amplicon sequence variants (ASVs) detected across the samples. The histograms show the number of samples that each taxon (ASV) was detected at each sediment depth. **(f)** Relative abundance of the twenty most abundant orders within the sediments, as well three others represented by genome bins; error bars show standard deviations of the mean and significance was tested using linear regression analyses with depth treated as a continuous variable (* *p* < 0.05, ** *p* < 0.01).

Community profiling indicated that the sands harbour diverse communities dominated by generalist taxa **(Table S1)**. There was considerable variation in taxonomic composition **(Fig. S3)** and beta diversity **(Fig. 1d)** across samples. These variations were moderately correlated with sediment depth *(R^2^* = 0.29) and weakly correlated with sampling date (*R*^2^ = 0.08) **(Fig. 1d; Table S2)**. Of the taxa (amplicon sequence variants, ASVs) detected, most exhibited low to intermediate occupancy, i.e. they were shared across several samples **(Fig. 1e & Fig. S4)**. Consistent with the theory that disturbance promotes community homogenisation, there was a higher number of shared taxa in the shallow sands (average occupancy of 3.3 samples; 71 ASVs shared across 15 samples) compared to deep sands (average occupancy of 2.3 samples; 0 ASVs shared across 15 samples). In line with previous observations^38^, the most abundant orders were Woeseiales (10.3 ± 4.7%) and Flavobacteriales (10.9 ± 5.1%), both of which were detected across all samples. Various other taxa, notably within the Pseudomonadales, Pirellulales, Microtrichales, Chitinophagales, and candidate gammaproteobacterial order GCA.001735895, were also prevalent and abundant **(Fig. 1f & Fig. S5)**. This suggests that these bacteria withstand large variations in habitat composition and resource availability in these sands. These bacterial groups were also the most abundant in metagenomes **(Table S3)**, based on community profiling using conserved single-copy ribosomal protein genes **(Table S4 & Fig. S6)**.

Nevertheless, there was evidence of some environmentally-driven differentiation in community composition. Community structure significantly differed between deep sediments and those of the shallow and intermediate sediments in the mixing zone **(Fig. 1d & Table S2)**, though overlapped at one sampling date. Consistently, the abundance of several orders significantly increased with depth, notably Desulfobacterales, Desulfobulbales, and Xanthomonadales **(Fig. 1f)**. Similarly, the proposed candidate phylum MBNT15^41^ was 20-fold more abundant in deeper samples based on amplicons **(Fig. 1f)** and metagenomes **(Fig. S5)**. This indicates that that the anoxic conditions of these sediments have selected for expansion of anaerobic specialists, including sulfate-reducing bacteria. However, read counts greatly varied across sampling dates; for example, while Desulfobacterales and Desulfobulbales attained relative abundances of 9% and 15% in the deep sediments, they were absent from the sediments of the same depth in the last two sampling dates. This indicates that these taxa, in contrast to the habitat generalists that they coexist with, are relatively sensitive to the disturbance events (e.g. oxygenation) that still occasionally affect deeper sediments. With respect to possible aerobic specialists, the orders Chitinophagales and Rhodobacterales were significantly more abundant in shallower sediments **(Fig. 1f)**. The latter order, which contains cultivated aerobic and photosynthetic members^42^, is likely to thrive under oxic photic conditions and may contribute to depth-related variations in chlorophyll *a* levels **(Fig. 1c)**.

### Metabolic flexibility differentiates habitat generalists and specialists

We used genome-resolved metagenomics to gain an insight into the metabolic traits of generalist and specialist community members. Sequencing, assembly, and binning of metagenomes of intertidal and subtidal sands from each sediment depth **(Table S3 & S5)** yielded 38 high-quality and 97 medium-quality metagenome-assembled genomes (MAGs)^43^ **(Table S6)**. We additionally reanalyzed the 12 MAGs that we previously reported from this study site^38^. Together, the resultant genomes span 13 phyla and 43 orders, including the most dominant taxa in the 16S profiles **(Fig. 1)**. We profiled the abundance of 44 marker genes in the short reads and derived genomes to gain an insight into the metabolic capabilities of the generalist and specialist community members **(Fig. 2)**. This confirmed microbial communities within sands adopt an extraordinary array of strategies for energy conservation.

**Figure 2.**
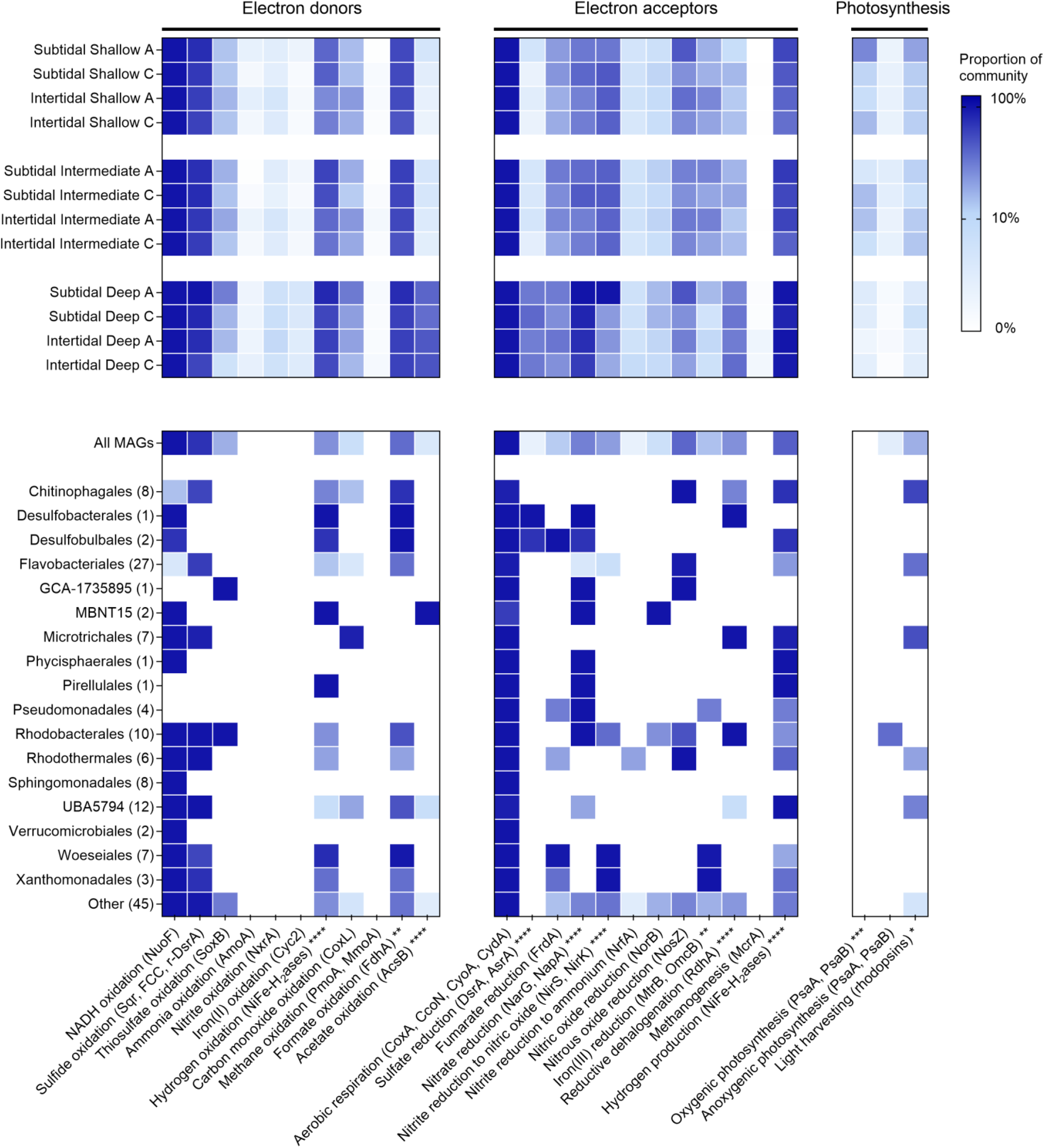
Metabolic capacity of microbial communities in permeable sediments. Homology-based searches were used to detect key metabolic genes in 12 metagenomes **(Table S3)** and 147 derived metagenome-assembled genomes (MAGs; **Table S6)**. The upper rows show the proportion of community members in each metagenome predicted to encode each gene based on the short reads; hits were normalized to gene length and single-copy ribosomal marker genes. Hits were summed for each process where more than one gene was searched for (up to 100%), with exception of oxygenic photosynthesis where PsaA and PsbA hits were averaged (reflecting both genes are required for this process to occur). The lower rows show the proportion of MAGs estimated to encode each gene, with results shown by order; hits are normalized based on estimated genome completeness. Metabolic marker genes involved in the oxidation of electron donors and reduction of electron acceptors are shown. Two-way ANOVAs were used to test whether there were significant differences in relative abundance of genes between depths (* *p* < 0.05, ** *p* < 0.01, *** *p* < 0.001, **** *p* < 0.0001 between shallow and deep sediments).

Most community members are predicted to be aerobic heterotrophs capable of using organic and inorganic energy sources. Based on short reads, most bacteria encoded enzymes for sulfide and thiosulfate oxidation, i.e. sulfide-quinone oxidoreductase (Sqr, 53%), flavocytochrome *c* sulfide dehydrogenase (FCC, 12%), reverse dissimilatory sulfite reductase (r-DsrA, 9%), and thiosulfohydrolase (SoxB, 16%) **(Fig. 2; Table S5)**. Concordantly, a similar proportion of the MAGs encoded these enzymes **(Fig. 2; Table S6)** and phylogenetic trees confirmed all binned sequences affiliated with canonical clades **(Fig. 3; Fig. S6 to S9)**. Most Sqr sequences, including from Woeseiales, Flavobacteriales, Rhodobacterales, and Microtrichales, affiliated with the type III clade **(Fig. 3a)** known to support sulfide-dependent growth^44,45^. Also widespread were the genes for consumption of carbon monoxide (CoxL, 19%; **Fig. S10**) and hydrogen gas (group 1 and 2 [NiFe]-hydrogenases, 48%; **Fig. S11**). Most bacteria also appear to have a large capacity to withstand variations in electron acceptor availability. In addition to encoding terminal oxidases for aerobic respiration **(Fig. 2)**, many are predicted to mediate stepwise denitrification through nitrate (NarG and NapA, 49%; **Fig. S12 & S13**), nitrite (NirS and NirK, 37%; **Fig. S14 & S15**), nitric oxide (NorB, 11 %; **Fig. S16**), and nitrous oxide (NosZ, 32%; **Fig. S17**), with fewer mediating dissimilatory nitrate reduction to ammonium (DNRA *via* NrfA, 7%; **Fig. S18**) **(Fig. 2)**. As we previously reported^38^, hydrogenotrophic sulfur reduction (group 1e [NiFe]-hydrogenases, 17%; **Fig. S11**) and facultative hydrogenogenic fermentation (group 3 [NiFe]-hydrogenases, 62%; **Fig. S19**) are also common. Diverse community members were also capable of reducing other compounds **(Table S5)**, such as ferric iron (MtrB, 20%; **Fig. S20)** and organohalides (RdhA, 21%; **Fig. S21)**. By contrast, few are predicted to mediate the specialist traits of ammonia, iron, nitrite, or methane oxidation, methanogenesis, acetogenesis, and, in the mixing zone, sulfate reduction **(Fig. 2; Table S5)**.

**Figure 3.**
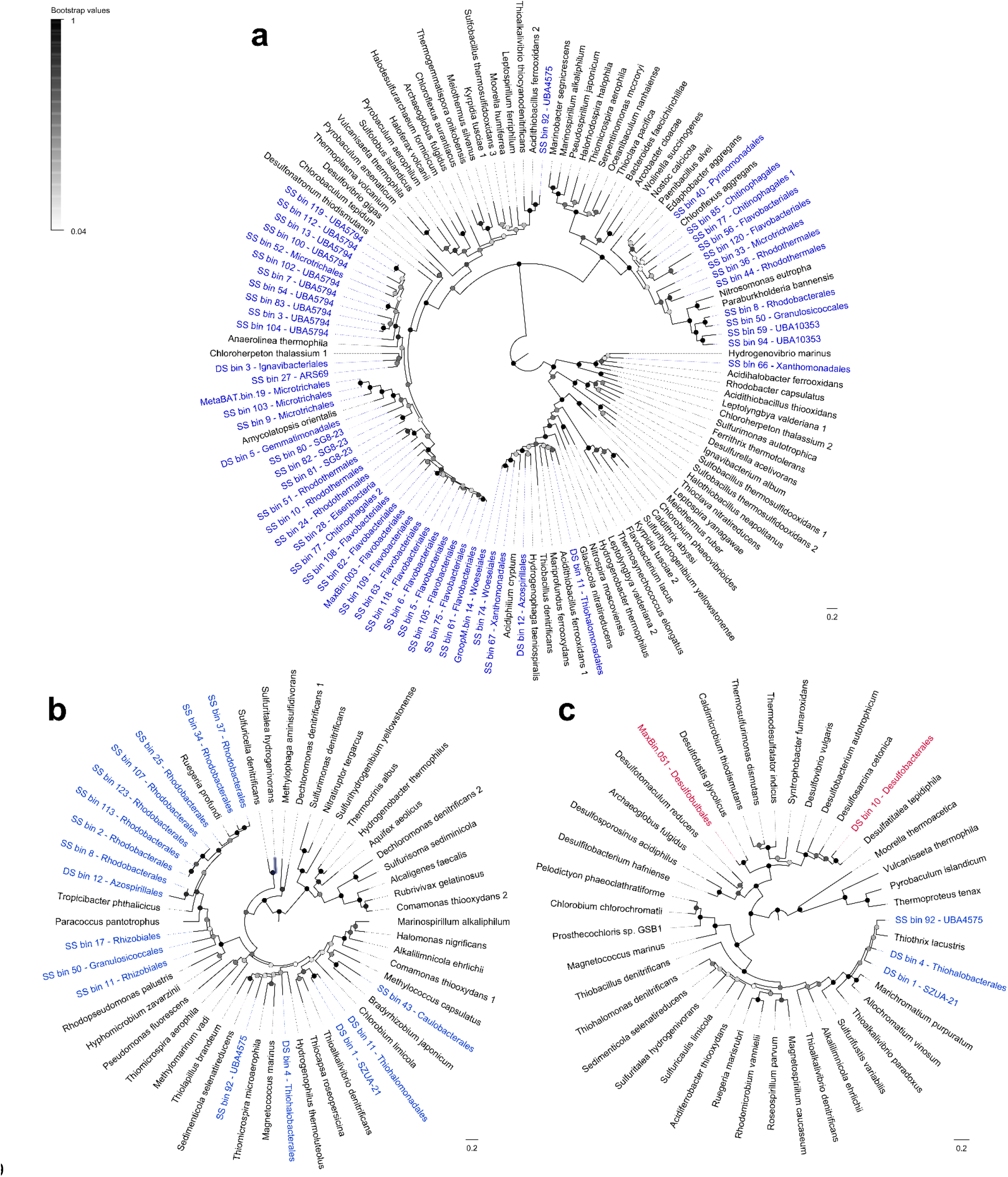
Phylogenetic trees of genes mediating sulfur cycling. Maximumlikelihood phylogenetic trees are shown for **(a)** sulfide-quinone oxidoreductase (Sqr), **(b)** flavocytochrome *c* sulfide dehydrogenase (FCC), and **(c)** dissimilatory sulfite reductase A subunit (DsrA). The tree shows sequences from permeable sediment metagenome-assembled genomes (colored) alongside representative reference sequences (black). The trees were constructed using the JTT matrix-based model, used all site, and were midpoint-rooted. Note Sqr, FCC, and the upper clade of DsrA (r-DsrA; encompassing Proteobacteria bins) are known to aerobic sulfide oxidation (bins colored in blue), whereas the middle clade of DsrA (encompassing Desulfobacterota bins) mediate anaerobic sulfite reduction (bins colored in red). Node junctions represent bootstrap support from 50 replicates. Full linear trees with accession numbers are provided in **Fig. S6** (Sqr), **Fig. S7** (FCC), and **Fig. S8** (DsrA).

Further analysis of the reconstructed genomes revealed that the most prevalent members are highly metabolically flexible **(Table S6 & Fig. 2)**. The Woeseiaceae bins, representing one of the most abundant and prevalent families in the sediments, encode enzymes for aerobic heterotrophy, aerobic sulfide oxidation, hydrogenotrophic sulfur reduction, denitrification, fumarate reduction **(Fig. S22)**, iron reduction, and hydrogenogenic fermentation. Flavobacteriaceae are similarly flexible, capable of harnessing energy from organic carbon, sulfide, formate, carbon monoxide, and sunlight via proteorhodopsin **(Fig. S23)**, as well as switching between aerobic respiration, anaerobic respiration, and fermentation. Other inferred generalists, including within highly abundant orders Pseudomonadales, Pirellulales, Microtrichales, Rhodothermales, and GCA-1735895 **(Fig. 1f)**, are also predicted to be able to use multiple energy sources and electron acceptors in these sediments **(Fig. 2)**. Altogether, this suggests most community members can accommodate environmental fluctuations in electron acceptor availability by switching between different respiratory and fermentative processes. Moreover, they can take advantage of a wide range of organic and inorganic energy sources that are likely to be abundant in these sediments. While most of the bacteria in the sediments were predicted to be flexible, we detected no alternative metabolic pathways across multiple near-complete MAGs from the Sphingomadales and Verrucomicrobiales **(Table S6)**; these bacteria may be aerobic organotrophic specialists, in line with their higher relative abundance in surface sands **(Fig. 1f)**.

The metagenomes also provide insights into the metabolic capabilities of community members with more restricted distributions (i.e. relative habitat specialists). Whereas the relative abundance of many generalist-associated genes (e.g. sulfide oxidation) did not change with depth, there was a significant fivefold increase in the relative abundance *(p* < 0.0001) of the marker genes for dissimilatory sulfate reduction (DsrA) **(Fig. 3c; Fig. S8)** and the Wood-Ljungdahl pathway (AcsB) in the metagenomes of deep sands compared to shallow and intermediate sands **(Fig. S20)**. This strongly correlates with the increased abundance of sulfate-reducing bacteria from the orders Desulfobulbales and Desulfobacterales at these depths **(Fig. 1f)** that encode these genes **(Fig. 2)**. These bacteria are likely able to thrive in this niche by coupling the oxidation of fermentative endproducts hydrogen (*via* group 1b and 1c [NiFe]-hydrogenases; **Fig. S11)** and acetate (through the oxidative Wood-Ljungdahl pathway; **Fig. S24)** to sulfate reduction. As highlighted in the phylogenetic trees of **Figure 3**, the genes for the inferred specialist process of sulfate reduction were far less abundant and widespread than those for sulfide oxidation. These sulfate-reducing orders nevertheless possess some respiratory flexibility, including the ability to use nitrate **(Fig. S13)** and organohalides **(Fig. S21)**, suggesting they can accommodate some changes in resource availability. However, in contrast to the facultative anaerobes that they coexist with, these obligate anaerobes are expected to be inhibited by oxygen given their terminal oxidases **(Fig. 2)** support detoxification rather than growth. Similarly, genome reconstructions indicate MBNT15 bacteria are obligate anaerobes that couple H_2_ and acetate oxidation to nitrate reduction. Thus, these lineages of Desulfobacterales, Desulfobulbales, and MBNT15 appear to be relative habitat specialists that thrive in anoxic deep sediments, but lack the metabolic capacity to compete in transiently oxygenated surface sediments.

### Metabolic processes associated with generalists and specialists show depth variations in permeable sediments

The above findings suggest that several alternative metabolic pathways, such as sulfide oxidation and hydrogenogenic fermentation, allow habitat generalists to adapt to changes in resource availability. The relative abundance of community members that mediate these processes, as well as the metabolic genes that they encode, is similar across depth **(Fig. 1f & 2)**. Thus, it can be expected that these processes occur in both shallow and deep sediments. To test this, we first measured rates of sulfide oxidation in sediments spiked with sodium sulfide under oxic conditions. Sulfide was rapidly consumed in a first-order kinetic process to below detection limits in both shallow and deep sediments **(Fig. 4c)**. We also measured hydrogenogenic fermentation in sands under anoxic conditions; glucose addition stimulated rapid accumulation of molecular hydrogen to micromolar levels in both surface and deep sands **(Fig. 4a)**.

**Figure 4.**
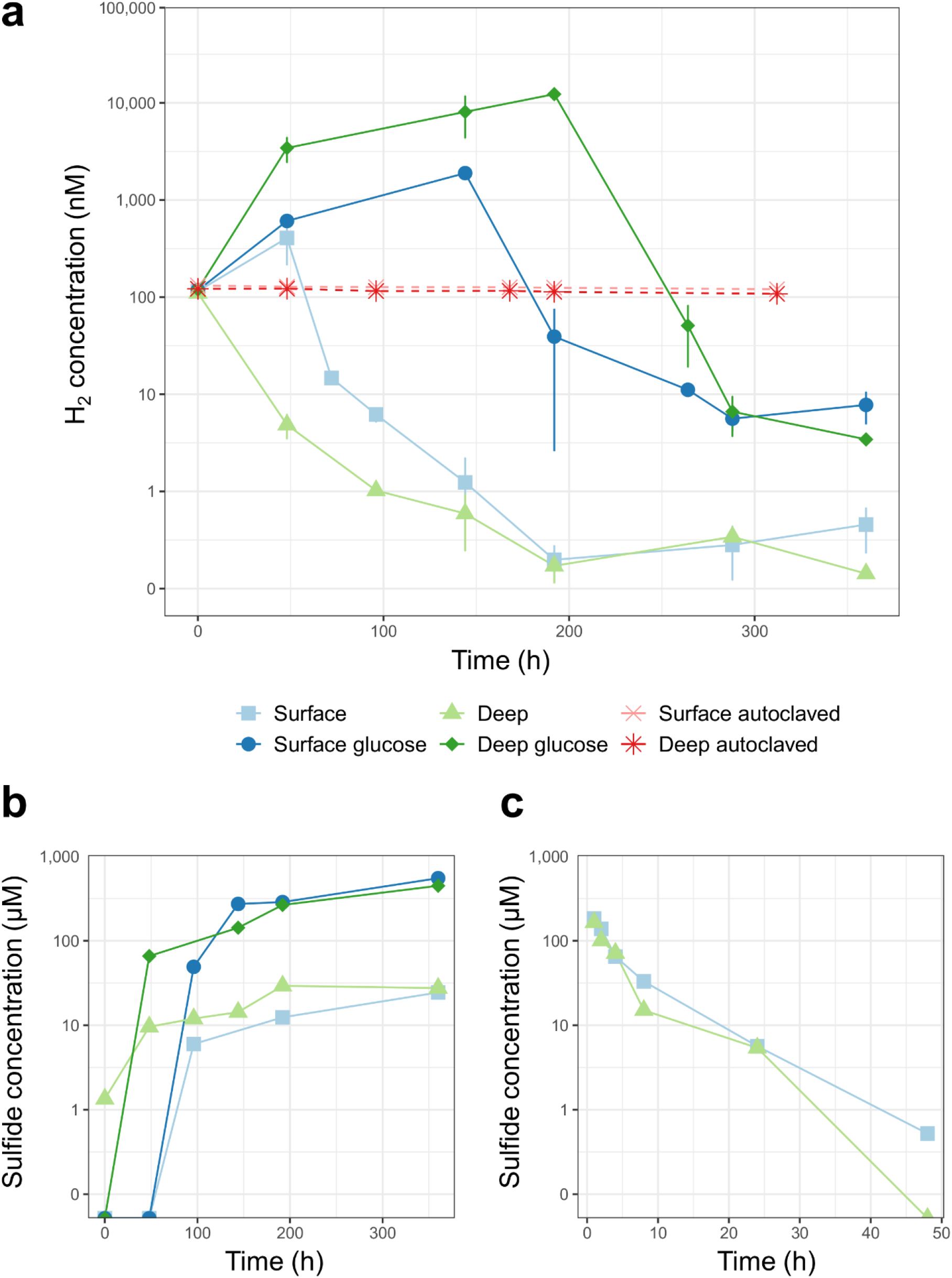
Metabolic activities of microbial communities in sediments. The first two panels show the capacity of sands to mediate hydrogenogenic fermentation and hydrogenotrophic sulfate reduction under anoxic conditions. Shallow and deep sediments were incubated in nitrogen-purged slurries in the presence of 100 ppmv H_2_ and, for spiked samples, 1 mM glucose. Changes in **(a)** H_2_ concentration and **(b)** sulfide concentration were measured during the experiment. For H_2_ measurements, error bars show standard deviations for three independent slurries. The third panel shows the capacity of oxic sands to mediate sulfide oxidation. Shallow and deep sediments were each incubated under oxic conditions in six independent slurries amended with 200 μM Na_2_S.9H_2_O. Changes in **(c)** sulfide concentration were measured during the timecourse, with one serum vial sacrificed per timepoint.

In contrast, the community and metagenome data indicate that sulfate reducers are habitat specialists that preferentially reside in the deeper sediments. To verify this, we measured rates of hydrogenotrophic sulfate reduction in anoxic H_2_-supplemented surface and deep sediments. As anticipated given the abundance of hydrogenotrophic sulfate reducers **(Fig. 1f)** and *dsrA* genes **(Fig. 2)**, the microbial communities in deep sediments consumed most H_2_ within 48 hours **(Fig. 4a)**, concomitant with accumulation of 10 μM sulfide **(Fig. 4b)**. In contrast, in line with our recent previous observations^38,40^, fermentation and respiration became uncoupled in surface sediments following the onset of anoxia; rates of fermentation initially exceeded respiration, resulting in net H_2_ accumulation and no detectable sulfide production within 48 hours. Hydrogenotrophic sulfate reduction only became dominant after prolonged incubations under anoxia **(Fig. 4a & 4b)**, likely due to growth of sulfate-reducing bacteria under these stable conditions.

### Metabolically flexible bacteria outcompete specialists during simulated disturbance events

The above insights from community, metagenomic, and biogeochemical profiling suggest that metabolic flexibility facilitates habitat generalism of microorganisms in permeable sediments. We performed a manipulative incubation experiment to test whether the above inferences are valid. Samples collected from shallow and deep sediments were incubated for 14 days under one of three conditions: continual light oxic conditions, continual dark anoxic conditions, and disturbed conditions (24-hour cycles between light oxic and dark anoxic conditions). Changes in the relative abundance of key orders previously highlighted in the community **(Fig. 1)** and metagenome **(Fig. 2)** analyses are shown in **Fig. 5**.

**Figure 5.**
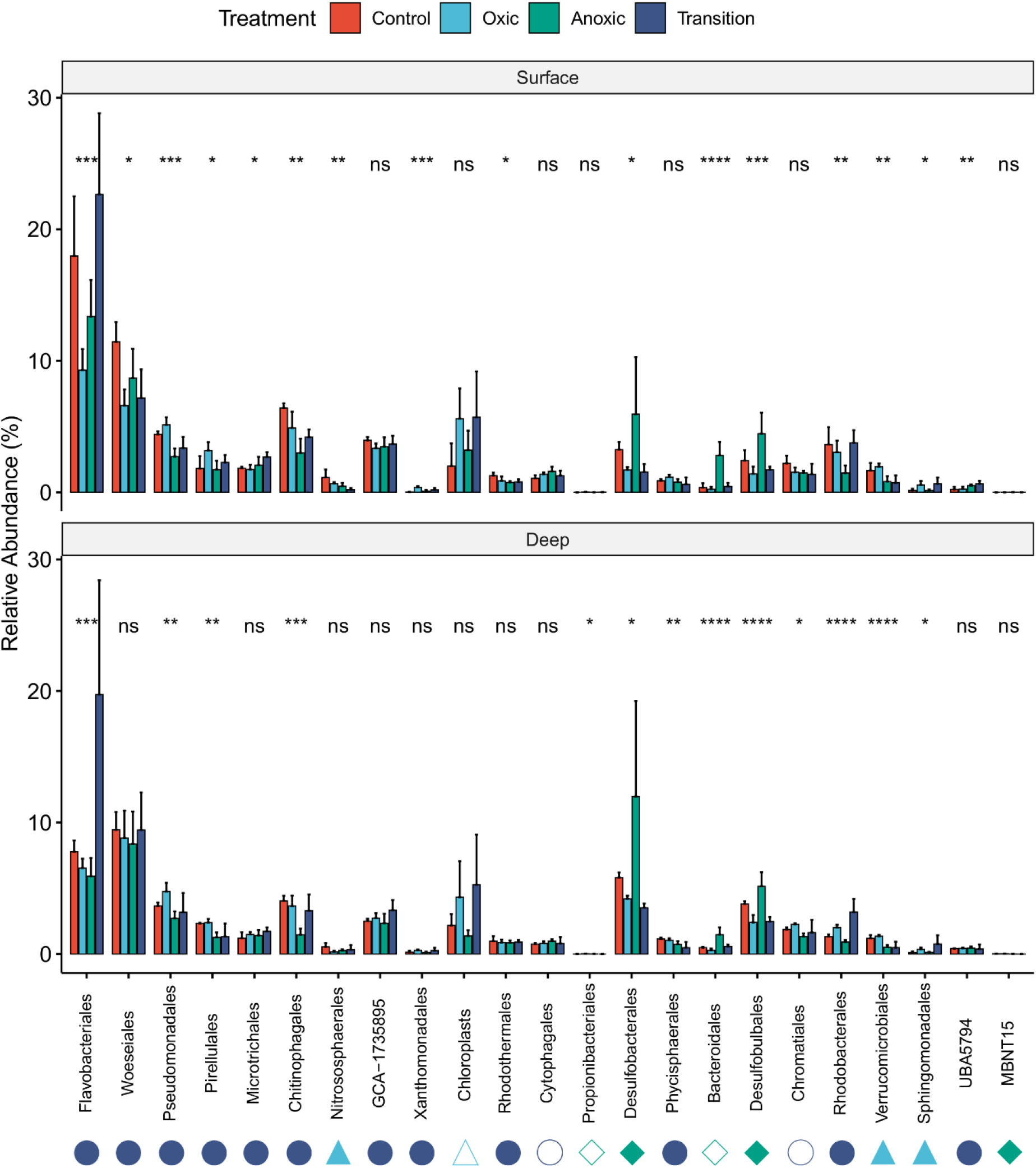
Responses of different orders to simulated environmental disturbance. The relative abundance of microbial orders from surface (top) and deep (bottom) sands is depicted with red bars. The changes of their relative abundance is shown after sands were incubated in slurries for two weeks in one of three conditions: continual light oxic conditions (light blue bars), continual dark anoxic conditions (green bars), or disrupted conditions (dark blue bars) in which slurries were shifted between light oxic and dark anoxic conditions every 24 hours. The 23 orders from **Figure 1** are depicted. Error bars show standard deviations of the mean and significance was tested using one-way ANOVAs (* *p* < 0.05, ** *p* < 0.01, *** *p* < 0.001, **** *p* < 0.0001). Shapes next to taxon names predict habitat preferences of each order based on the obtained bins: generalists (dark blue circles), oxic specialists (light blue triangles), and anoxic specialists (green diamonds). Given no bins were obtained for Cytophagales, Propionibacterales, Bacteroidales, Chromatiales, and chloroplasts, metabolic capabilities are inferred based on cultured organisms.

Most orders predicted to be metabolically flexible were able to tolerate being incubated under all three conditions. These inferred generalists were dominant in all samples, with highest relative abundances compared to inferred specialists in the original samples and disturbed incubations **(Fig. 5)**. Reflecting this, there were relatively minor changes in the relative abundance of Woeseiales, Microtrichales, Rhodothermales, and GCA_1735895 between time of sampling and following two weeks of incubations. We also monitored the patterns of lineages predicted to be aerobic specialists (Verrucomicrobiales, Sphingomondales) and anaerobic specialists (Desulfobacterales, Desulfobulbales, Bacteroidales) based on the reconstructed genomes **(Fig. 2)**. Consistent with expectations, the relative abundance of both groups declined by 40% in the disturbed slurries compared to the original samples. The inferred aerobic specialists, while always relatively minor community members, were most abundant in oxic incubations (4.7%) and least in anoxic sediments (2.1%). Inferred anoxic specialists showed the reciprocal pattern. They bloomed to one third of the community in the anoxic incubations (30%), but declined during oxygen exposure (8%) **(Fig. 5)**. It is likely that, under stable anoxic conditions, these anaerobic specialists rapidly mobilize available resources through their sulfate reduction and fermentation pathways.

Remarkably, some taxa thrived in response to disturbance. Flavobacteriales sampled from deep sediments increased in relative abundance by 2.5-fold in the disturbed incubations **(Fig. 5)**, largely driven by expansions of genus *Eudoraea* **(Table S7)**. Based on the metabolic capabilities of the three MAGs from this genus **(Table S6)**, it is possible such bacteria take advantage of necromass released during oxic-anoxic transitions by switching between aerobic respiration and hydrogenogenic fermentation pathways. Likewise, there were significant enrichments in the two dominant groups of phototrophs in the sediments, namely Rhodobacterales and diatoms (detected by chloroplast 16S sequences) **(Fig. 1f & Table S5)**. These taxa likely benefit from the increased light availability under both the light oxic and disturbed conditions compared to natural sediments, but must also possess sufficient metabolic flexibility to persist under dark anoxic conditions; such flexibility is apparent from the diverse repertoire of Rhodobacterales MAGs **(Fig. 2)**, as well as previous studies inferring diatoms survive dark anoxic conditions through nitrate respiration^46^ and microbiota-mediated hydrogenogenic fermentation^38,40^. Although this experiment generally substantiated metagenome-based inferences, a few taxa behaved contrary to predictions. Most notably, Chitinophagales significantly decreased under anoxic conditions despite harbouring genes for hydrogenogenic fermentation **(Fig. S18)**, suggesting members of this order either cannot survive in these conditions or are outcompeted by more efficient anaerobes; these observations are nevertheless consistent with the significant decrease in the relative abundance of this order with depth **(Fig. 1f)**.

### Broader ecological and biogeochemical significance of findings

In combination, these results provide multifaceted evidence that environmental disturbance influences distributions of microbial habitat generalists and specialists. The microbial communities in the mixing zone of permeable sediments experience frequent but irregular spatiotemporal variations in oxygen, sunlight, nutrients, and redox state^25^. Based on ecological theory, it would be expected that these variations would differentially affect generalists and specialists^1,4^. For the specialists, these changes would promote continual cycles of growth and death as conditions alternate between favourable and unfavourable. In contrast, generalists are expected to maintain more stable populations given they are more adaptable to environmental change. We observed that habitat generalists are indeed more competitive in these environments. Large and stable populations of taxa such as Woeseiales, Flavobacteriales, and Pseudomonadales were present in both the mixing and deep layers of the sampled sediments across sampling times, and were enriched under simulated disturbance conditions in the manipulative incubations. Thus, in line with observations for macroorganisms, environmental disturbance appears to promote homogenisation of microbial community composition.

Some relative habitat specialists nevertheless coexist with such generalists in these environments. Numerous taxa were detected with low occupancies and abundances, several of which bloomed under favourable conditions, most notably MBNT15. The manipulative incubation experiments confirmed that these inferred specialists only became enriched under more stable conditions (light oxic for aerobes, dark anoxic for anaerobes). Most notably, Desulfobacterales were the most abundant order in deep sediments at certain sampling times and during prolonged dark anoxic incubations, reflecting that sulfate-reducing bacteria thrive in stable hydrogen- and sulfate-rich environments. These taxa and other anaerobic specialists nevertheless exhibited sharp variations in relative abundance across the sampling dates, as well as significant declines under oxic and disturbed incubations. In line with ecological paradigms, this suggests that such specialists are highly sensitive to the disturbances that define the mixing zone and occasionally affect deeper sands, whereas the generalists that they coexist with are more adaptable. More data is required across various spatial and temporal scales to ultimately understand the physicochemical pressures and biological interactions that drive these differences. However, it is probable that oxygen availability is the most significant factor that regulates composition, for example through causing poisoning or promoting outcompetition by generalists better adapted to these conditions^38^.

In turn, our study lends strong support to the hypothesis that microbial habitat generalists and specialists have distinct metabolic capabilities. Based on the reconstructed genomes, the generalists in the community are predicted to be extremely metabolically flexible. Most notably, the Woeseiaceae that dominate these sands are among the most flexible microorganisms ever described, given they are predicted to use wide spectrum of electron donors (organic carbon, sulfide, hydrogen), oxidants (oxygen, nitrite, fumarate, sulfur, fermentation), and based on previous analysis^21^, carbon sources (heterotrophy, autotrophy). Flavobacteriaceae have similar metabolic breadth, likely underlying their expansion in response to disturbance. By contrast, relative habitat specialists from the Desulfobacterales and Desulfobulbales are distinguished by their capacity to the use the abundant electron acceptor sulfate, but also their inability to grow by aerobic respiration. These bacteria possess some metabolic flexibility, likely explaining why these orders were detected in low levels even in most surface sediments and oxygenated slurries; indeed, habitat generalism and metabolic flexibility alike should be considered as continuous traits. However, such obligate anaerobes are outcompeted by facultative anaerobes under disturbed conditions. These inferred differences were strongly supported by biogeochemical assays showing that, whereas sulfate reduction is limited to sediments under prolonged anoxia, metabolic traits associated with generalists are active through sediment zones. Further culture-dependent and culture-independent work, however, is required to comprehensively understand the metabolic capabilities of permeable sediment bacteria and their responses to environmental changes.

These findings also have important implications for how we conceive and model biogeochemical processes. Models describing these processes can either take an organism-centric approach or a systems perspective^47^ In the first case, the presence or absence of a particular organism will determine the process taking place and emphasis is placed on modelling the growth of that organism. In the second case, thermodynamics and physical conditions determine the processes taking place. Biogeochemists typically use the second approach to successfully predict and model sediment processes^48^ Under conditions of continual disturbance, we show that generalists dominate, and the energy conservation pathways that are used (particularly under anaerobic conditions) will not be those predicted from thermodynamics until specialists dominate (such as sulfate reduction). Under disturbed conditions, therefore, community structure and the presence of generalists (the organism-centric view) becomes an important consideration for predicting ecosystem processes. Consistent with this, it has been shown that physicochemical variables are strongest predictor of microbially driven ecosystem processes, but that microbial community structure can improve these predictions in some cases^49^. Future studies should incorporate disturbance as a co-variate when comparing the efficacy of organism and system scale models (both statistical and deterministic).

In combination, we conclude that habitat generalists thrive in the disturbed environments of permeable sediments and generally outcompete specialists. This reflects their greater metabolic flexibility, particularly their capacity to shift between electron acceptors during oxic-anoxic transitions. Relative habitat specialists have narrower niches, but are highly competitive under more stable conditions. These findings are substantiated through community and metagenomic profiling, biogeochemical measurements, and manipulative experiments. Thus, a long-standing ecological theory explaining differential distribution patterns of macroorganisms appears to extend to microorganisms and we provide a mechanistic rationale for these observations. Though further studies are required to extend these findings beyond permeable sediments, it is probable that metabolic flexibility is a key factor governing distributions of generalist and specialist taxa across ecosystems.

## Materials and Methods

### Sampling of permeable sediments

Permeable sediments were sampled from Middle Park Beach, Port Phillip Bay, Sediments for microbial community profiling were collected over eight different sampling dates over the course of a year (A: 28/10/2016; B: 13/12/2016; C: 19/1/2017; D: 28/3/2017; E: 9/5/2017; F: 30/6/2017; G: 23/8/2017; H: 19/10/2017). Cores of 30 cm were used to collect sediments from the subtidal zone (~1 m deep at low tide) and intertidal zone (~1 m deep at high tide). Cores were kept on ice until delivery to the laboratory and were then immediately sectioned into shallow (0-3 cm), intermediate (14-17 cm), and deep (27-30 cm) samples. All samples were subsequently stored at −20°C until further processing.

### Amplicon sequencing

For amplicon sequencing, total community DNA was extracted from 0.25 g of sediment using the modified Griffith’s protocol^50^ The yield, purity, and integrity of DNA from each extraction was confirmed using a Qubit Fluorometer, Nanodrop 1000 Spectrophotometer, and agarose gel electrophoresis. For each sample, the V4 hypervariable region for 16S rRNA gene was amplified using the universal Earth Microbiome Project primer pairs F515 and R806^51^ and subjected to Illumina paired-end sequencing at the Australian Centre for Ecogenomics, University of Queensland. Paired-end raw reads were demultiplexed and adapter sequences were trimmed, yielding 1,362,535 reads across all samples. Forward and reverse sequences were merged using the q2-vsearch plugin^52^. A quality filtering step was applied using a sliding window of four bases with an average base call accuracy of 99% (Phred score 20). The reads were truncated down to 250 base pairs to remove low quality reads before de-noising using the deblur pipeline^53^ in QIIME 2^54^. Samples with read counts less than 1000 were removed from the further analysis. A total of 42 samples remained after removing six samples. Amplicon sequence variants (ASVs) occurring once were removed from the dataset. A total of 12,566 ASVs remained after removing 270 singletons. For taxonomic assignment, all reference reads that matched the F515/R806 primer pair were extracted from the Genome Taxonomy Database (GTDB)^41^ and used to train a naïve bayes classifier by using the fit-classifier-naive-bayes function with default parameters.

### Biodiversity analysis

All statistical analysis and visualizations were performed with R software version 3.5.0 (April 2018) using the packages phyloseq^55^, vegan^56^, and ggplot2^57^ Prior to statistical analysis, all sequences were rarefied at 5,000 sequences per sample. Alpha diversity was calculated using several metrics, including Shannon index, which measures both species richness and evenness. We tested for significant differences in Shannon index between depth, tidal zone, and date using an ANOVA (one-way analysis of variance) with Tukey’s *post hoc* tests (*p* < 0.05). Beta diversity was calculated using weighted UniFrac distances^58^ of log10-transformed data and visualized using principal coordinate analysis (PCoA). A pairwise analysis of similarities (ANOSIM) was used to test for significant differences in community similarity between depths, tidal zone, and date. First, permutational multivariate analysis of variance (PERMANOVA) was performed using 999 permutations to test for significant differences. Second, a beta dispersion test (PERMDISP) was used to ascertain if observed differences were influenced by dispersion.

### Quantitative PCR

Quantitative PCR (qPCR) was used to absolutely quantify the copy number of the 16S rRNA genes in the samples. Amplifications were performed using a 96-well plate in a pre-heated LightCycler^®^ 480 Instrument II (Roche, Basel, Switzerland). Each well contained a 10 μl reaction mixture comprising 1 μl DNA template, 5 μl PlatinumT SYBRGreen qPCR SuperMix-UDG with ROX, 0.5 μl each of the universal 16S rRNA gene V4 primers F515 and R806 (10 μM)^51^, and 3 μl UltraPure Water (Thermo Fisher Scientific, Waltham, MA, USA). Each amplification was performed in technical triplicate. Cycling conditions were as follows: 3 min denaturation at 94°C followed by 40 cycles of 45 s denaturation at 94°C, 60 s annealing at 50°C, and 90 s extension at 72°C. Copy number was quantified against a pMA vector standard containing a single copy of the *Escherichia coli* 16S rRNA gene. Dilutions ranged from 10^3^ to 10^8^ copies μl^-1^ and the qPCR amplification efficiency ranged from 85-94% (R^2^ > 0.99).

### Chlorophyll *a* measurements

Chlorophyll *a* was extracted using a previously described method^59^ Briefly, 5 mL of 90% acetone (v/v) was added to 5 g of sediments in 50 ml Falcon tubes. Samples were then stored overnight in the dark at 4°C. Subsequently, all samples were subsequently centrifuged at 550 × *g* for 15 minutes and 3 mL of supernatant was transferred into cuvettes. Chlorophyll absorbance was measured spectrophotometrically using a Hitachi U-2800 spectrophotometer (Hitachi High-Technologies Corporation, Tokyo, Japan) at five different wavelengths (630, 647, 664, 665, and 750 nm). Spectra were read before and after acidification with 10 μL of 1 M HCl (v/v). After calculating the difference in absorbance between the first and second measurement, chlorophyll *a* concentration was determined using the equation of Lorenzen^59^.

### Shotgun metagenome sequencing

**Table S1** summarizes details of the metagenomic datasets. For this study, we sequenced eight new metagenomes (subtidal deep A, intertidal deep A, subtidal shallow C, intertidal shallow C, subtidal intermediate C, intertidal shallow C, subtidal deep C, intertidal deep C) and analyzed five previously reported metagenomes (subtidal shallow A, subtidal intermediate A, intertidal shallow A, intertidal intermediate A, flow-through reactor)^38^ DNA was extracted from the 0.3 g of sediment, collected during the October 2016 (A samples) and January 2017 (C samples) field trips, using the MoBio PowerSoil Isolation kit according to manufacturer’s instructions. Metagenomic shotgun libraries were prepared for each sample using the Nextera XT DNA Sample Preparation Kit (Illumina Inc., San Diego, CA, USA) and sequencing was performed on an Illumina NextSeq500 platform with a 2 × 150 bp High Output run. Sequencing yielded 574,093,137 read pairs across the eight metagenomes. To supplement the 16S amplicon sequencing data, community profiles in permeable sediments were independently generated from metagenome reads that mapped to the universal single copy ribosomal marker gene *rplP* using SingleM v.0.12.1 (.https://github.com/wwood/singlem)

### Shotgun metagenome assembly and binning

The BBDuk function of the BBTools v38.51 (https://sourceforge.net/projects/bbmap/) was used to clip contaminating adapters (k-mer size of 23 and hamming distance of 1), filter PhiX sequences (k-mer size of 31 and hamming distance of 1), and trim bases with a Phred score below 20 from the raw metagenomes. 482,529,838 high-quality read pairs with lengths over 50 bp were retained for downstream analysis. Reads were assembled individually and collectively with MEGAHIT v1.2.9^60^ (--k-min 27, --k-max 127, --k-step 10). Bowtie2 v2.3.5^61^ was used to map short reads back to assembled contigs using default parameters to generate coverage profiles. Subsequently, genomic binning was performed using CONCOCT v1.1.0^62^, MaxBin2 v2.2.6^63^, and MetaBAT2 v2.13^64^ and bins from the same assembly were then dereplicated using DAS_Tool v1.1^65^ Spurious contigs with incongruent genomic and taxonomic properties and 16S rRNA genes in the resulting bins were removed using RefineM v0.0.25^66^ Applying a threshold average nucleotide identity of 99%, bins from different assemblies were consolidated to a non-redundant set of metagenome-assembled genomes (MAGs) using dRep v2.3.2^67^ Completeness and contamination of MAGs were assessed using CheckM v1.1.2^68^ In total, 38 high quality (completeness > 90% and contamination < 5%) and 97 medium quality (completeness > 50% and contamination < 10%)^43^ MAGs were recovered and their corresponding taxonomy was assigned by GTDB-TK v1.0.2^41^. Open reading frames (ORFs) in MAGs were predicted using Prodigal v2.6.3 metagenomic setting^69^

### Shotgun metagenome functional analysis

To estimate the metabolic capability of the sediment communities, metagenomes and derived genomes were searched against custom protein databases of representative metabolic marker genes using DIAMOND v.0.9.22 (query cover > 80%)^70^. Searches were carried out using all quality-filtered unassembled reads with lengths over 140 bp. In addition, we searched ORFs from the 135 MAGs retrieved from this study and 12 MAGs that were previously reported^38^. These genes are involved in sulfur cycling (AsrA, FCC, Sqr, DsrA, Sor, SoxB), nitrogen cycling (AmoA, HzsA, NifH, NarG, NapA, NirS, NirK, NrfA, NosZ, NxrA, NorB), iron cycling (Cyc2, OmcB), reductive dehalogenation (RdhA), photosynthesis (PsaA, PsbA, energy-converting microbial rhodopsin), methane cycling (McrA, MmoA, PmoA), hydrogen cycling (large subunit of NiFe-, FeFe-, and Fe-hydrogenases), carbon monoxide oxidation (CoxL), succinate oxidation (SdhA), fumarate reduction (FrdA), and acetogenesis (AcsB)^71–73^ Results were further filtered based on an identity threshold of 50%, except for group 4 NiFe-hydrogenases, FeFe-hydrogenases, CoxL, AmoA, and NxrA (all 60%), PsaA (80%), and PsbA (70%). Subgroup classification of reads was based on the closest match to the sequences in databases. The presence of an additional set of genes involved in oxidative phosphorylation (AtpA), NADH oxidation (NuoF), aerobic respiration (CoxA, CcoN, CyoA, CydA), formate oxidation (FdhA), arsenic cycling (ARO, ArsC), iron cycling (MtrB), selenium cycling (YgfK) in MAGs and contig ORFs was screened by hidden Markov models (HMM)^74^, with search cutoff scores as described previously^75^ Resulting hits were manually inspected to remove false positives. The screening of these genes in uinassembled reads was carried out using DIAMOND blastp algorithm (using binned and contig hits as reference sequences) with a minimum percentage identity of 60% (NuoF), 70% (AtpA, FdhA, ARO), or 50% (all other databases). Read counts to each gene were normalized to reads per kilobase million (RPKM) by dividing the actual read count by the total number of reads (in millions) and then dividing by the gene length (in kilobases; based on average gene length in custom databases and gene length of representative sequence in Swiss-Prot database for HMMs were used). In order to estimate the gene abundance in the microbial community, high-quality unassembled reads were also screened for the 14 universal single copy ribosomal marker genes used in SingleM v.0.12.1 and PhyloSift^76^ by DIAMOND (query cover > 80%, bitscore > 40) and normalized as above. Subsequently, the average gene copy number of a gene in the community can be calculated by dividing the read count for the gene (in RPKM) by the geometric mean of the read count of the 14 universal single copy ribosomal marker genes (in RPKM).

### Phylogenetic analysis

Phylogenetic analysis was used to verify the presence of key metabolic genes in permeable sediment MAGs and determine which lineages were present. Phylogenetic trees were constructed for 18 genes involved in energy conservation: dissimilatory sulfite reductase (DsrA), sulfide-quinone oxidoreductase (Sqr), flavocytochrome *c* sulfide dehydrogenase (FCC), thiohydrolase (SoxB), acetyl-CoA synthase (AcsB), carbon monoxide dehydrogenase (CoxL), group 1 [NiFe]-hydrogenases, group 3 [NiFe]-hydrogenases, two nitrate reductases (NarG, NapA), three nitrite reductases (NirS, NirK, NrfA), nitric oxide reductase (NorB), nitrous oxide reductase (NosZ), decaheme iron reductase (MtrB), reductive dehalogenase (RdhA), fumarate reductase (FrdA), and energy-converting microbial rhodopsins. In all cases, protein sequences retrieved from the MAGs by homology-based searches were aligned against a subset of reference sequences from the custom protein databases using ClustalW^77^ in MEGA7^78^. Evolutionary relationships were visualized by constructing maximum-likelihood phylogenetic trees; specifically, initial trees for the heuristic search were obtained automatically by applying Neighbour-Join and BioNJ algorithms to a matrix of pairwise distances estimated using a JTT model, and then selecting the topology with superior log likelihood value. All residues were used and trees were bootstrapped with 50 replicates.

### Biogeochemical experiments

Slurry experiments were performed to investigate the functional capacity of surface and deep sands. Each slurry comprised a 160 mL serum vial containing 30 g of sieved sand (wet weight) and 70 mL of seawater (filtered on 0.45 μm Whatman membrane filters). The serum vials were sealed with butyl rubber stoppers and Wheaton closed-top seals. Anoxic slurries were used to measure hydrogenogenic fermentation and sulfate reduction in shallow and deep sands collected on November 12, 2018. Briefly, the slurries were purged with high-purity helium and the headspace was amended with 100 ppmv H_2_. Glucose was added to a final concentration of 1 mM for the glucose addition group. All vials were incubated on a shaker (100 rpm) at room temperature. For H_2_ measurements, a 2 mL subsample was collected from headspace every 24 h and analysed by gas chromatography. For sulfide measurements, a total of 8 mL of seawater was extracted from each slurry and filtered for spectrophotometric analysis. Three independent slurries were performed per treatment. Oxic slurries were used to measure aerobic sulfide oxidation in shallow and deep sands collected on December 6, 2018. The serum vials were aerated with lab air and sodium sulfide (Na_2_S.9H_2_O) was added to a final concentration of 500 μM. All vials were incubated on a shaker (100 rpm) at room temperature. A total of 8 mL of seawater was extracted from each slurry and filtered for spectrophotometric analysis. The autoclaved vial was used as the control group to control for the photochemical oxidation of sulfide in aqueous solution. The amount of biogenic sulfide oxidation that occurred between each timepoint was determined by calculating the difference between the treatment and control groups.

### Molecular hydrogen and sulfide measurements

To measure molecular hydrogen (H_2_), 2 mL gas samples extracted during the slurry experiments were injected into a VICI Trace Gas Analyser (TGA) Model 6K (Valco Instruments Co. Inc., USA) fitted with a pulsed discharge helium ionisation detector (PDHID) as previously described^79^. Ultra-pure helium (99.999% pure, AirLiquide) was used as a carrier gas at a pressure of 90 psi. The temperatures of column A (HayeSep DB), column B (Molesieve 5Å), and the detector were 55 °C, 140 °C and 100 °C respectively. The instrument was calibrated using standards of ultra-pure H_2_ (99.999% pure, AirLiquide) in ultra-pure He. Sulfide concentrations were quantified through the methylene blue method with GBC UV-Visible 918 Spectrophotometer at 670 nm as previously described^80^.

### Long-term incubation experiments

A long-term incubation experiment was performed to compare how habitat stability and variability affects the community structure of permeable sediments. Surface (0-3 cm) and deep (20-25 cm) sediments were collected from Middle Park beach on October 9, 2019. They were incubated in slurries comprising a 160 mL serum vial containing 30 g of sieved sand (wet weight) and 70 mL of seawater (filtered on 0.45 μm Whatman membrane filters). The vials were sealed with butyl rubber stoppers and Wheaton closed-top seals. All vials were incubated on a shaker (100 rpm) at room temperature. Three different treatments were applied for both surface and deep. For the light oxic slurries, vials were aerated daily with laboratory air and continuously exposed to 60 μmol photons m^-2^ s^-1^. For the dark anoxic slurries, vials were purged with high-purity nitrogen gas and covered with aluminium foil. For the oxic-anoxic transition slurries, vials were transferred between light oxic to dark anoxic conditions every 24 hours. All incubations were performed in triplicate. DNA was extracted from the original sediments (control group) and each slurry after 14 days of incubation. Community structure was determined by 16S amplicon sequencing as described above.

## Footnotes

### Data availability statement

All amplicon sequencing data, raw metagenomes, and metagenome-assembled genomes will be deposited to the Sequence Read Archive.

## Acknowledgements

This study was supported by an ARC Discovery Project (DP180101762; awarded to P.L.M.C. and C.G), an ARC DECRA Fellowship (DE170100310; salary for C.G.), an ARC Laureate Fellowship (FL150100038; awarded to P.H.), and an NHMRC EL2 Fellowship (APP1178715; salary for C.G.). Y.J.C. was supported by PhD scholarships from Monash University and the Taiwan Ministry of Education. We thank Dustin Marshall and Kim Handley for helpful discussions, Rachael Lappan for providing R scripts, and Thanavit Jirapanjawat, David Brehm, Vera Eate, Sharlynn Koh, Tess Hutchinson, and Zahra Islam for technical and field assistance.

## Author contributions

C.G. and P.L.M.C. conceived, designed, and supervised this study. Y.-J.C. designed experiments, performed all field and laboratory work, and analyzed data. P.M.L., C.G., D.J.W., and P.H. analysed metagenomes. G.S. and S.K.B. provided experimental and analytical support. A.J.K. contributed to conceptualisation, analysis, and interpretation. Y.-J.C. and C.G. wrote the paper with input from all authors.

The authors declare no conflict of interest.

