## Supplementary Information for "Metabolic flexibility allows generalist bacteria to become dominant in a frequently disturbed ecosystem"

### Supplementary material

**Figure S1.** Comparison of Shannon index based on the 16S amplicon sequencing results across **(a)** eight sampling dates and **(b)** two tidal types.

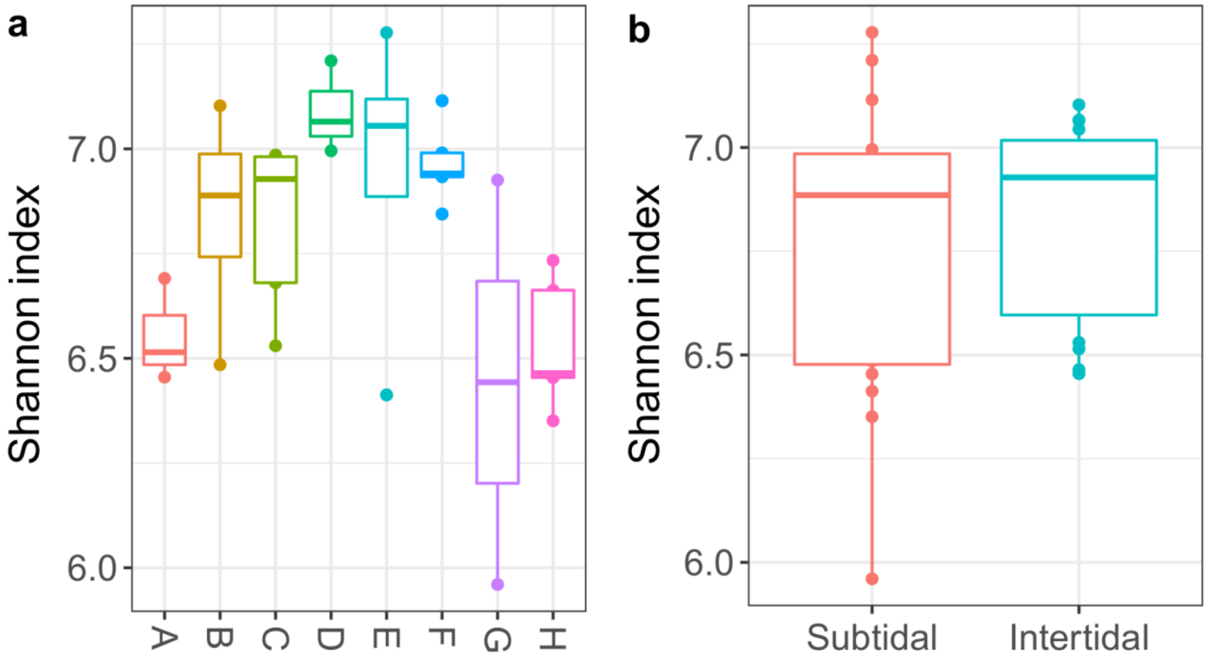

**Figure S2.** Different alpha diversity measurements based on the 16S amplicon sequencing results across **(a)** sediment depth and **(b)** sampling date.

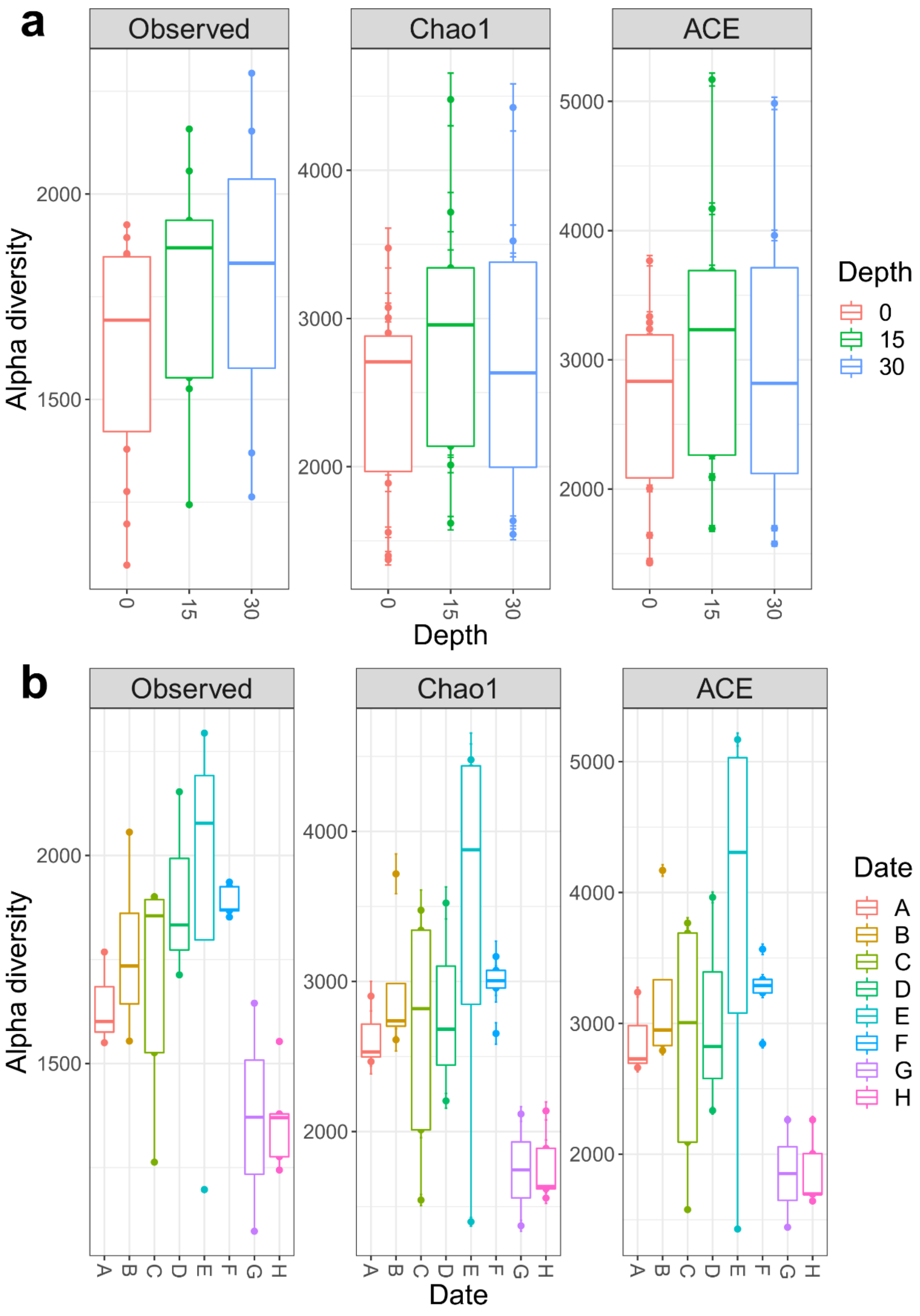

**Figure S3.** Microbial communities of sediments at the level of phylum. Communities were sampled across eight different sampling dates (A: 28/10/2016; B: 13/12/2016; C: 19/1/2017; D: 28/3/2017; E: 9/5/2017; F: 30/6/2017; G: 23/8/2017; H: 19/10/2017), two different tidal zones (I: intertidal; S = subtidal), and three different sediment depths (0 = 0-3 cm, 15 = 14-17 cm, 30 = 27-30 cm).

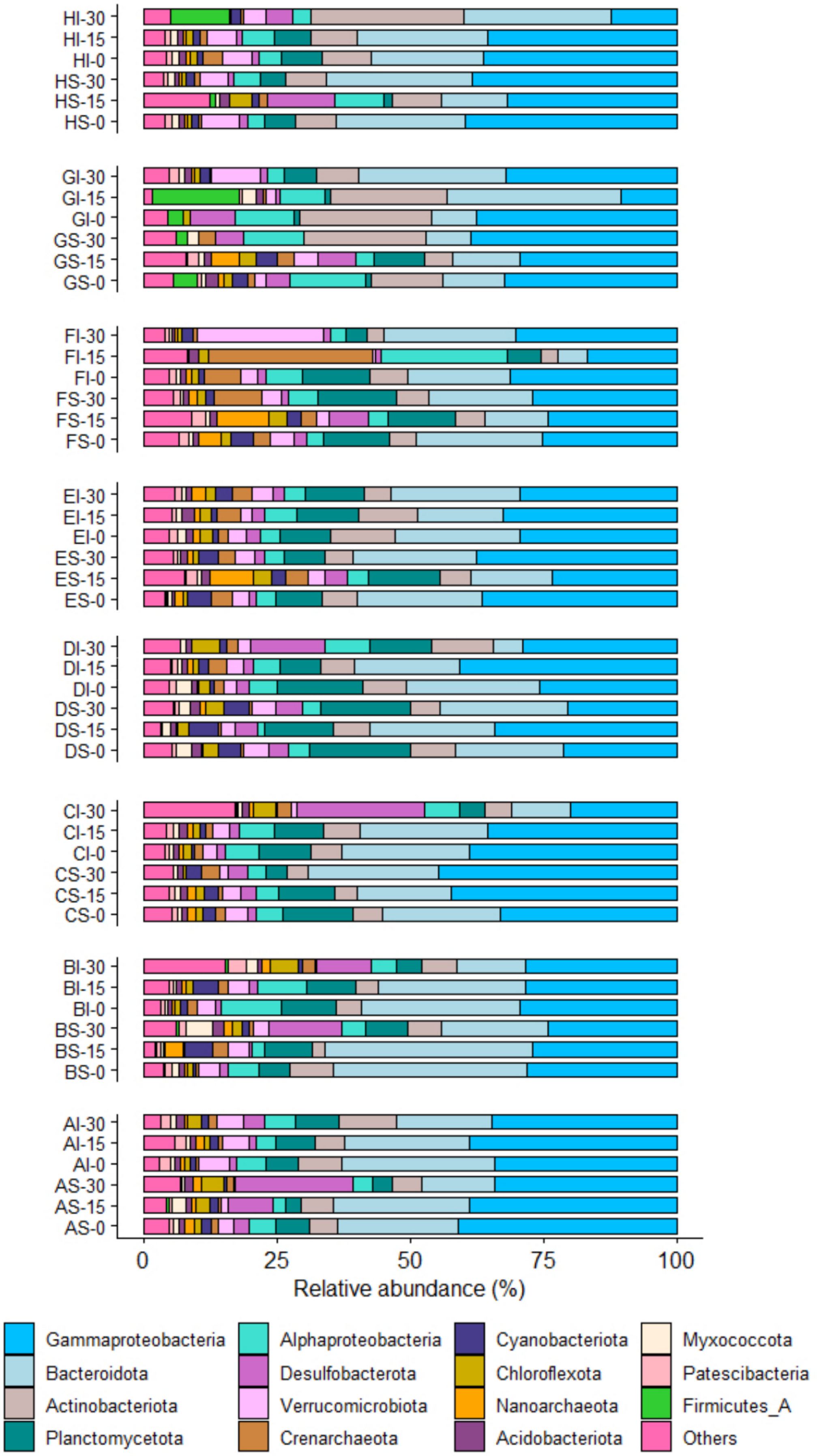

HI-30
HI-15
HI-0
HS-30
HS-15
HS-0
GI-30
GI-15
GI-0
GS-30
GS-15
GS-0
FI-30
FI-15
FI-0
FS-30
FS-15
FS-0
EI-30
EI-15
EI-0
ES-30
ES-15
ES-0
DI-30
DI-15
DI-0
DS-30
DS-15
DS-0
CI-30
CI-15
CI-0
CS-30
CS-15
CS-0
BI-30
BI-15
BI-0
BS-30
BS-15
BS-0
AI-30
AI-15
AI-0
AS-30
AS-15
AS-0
0
25
50
75
100
Relative abundance (%)
Gammaproteobacteria
Bacteroidota
Actinobacteriota
Planctomycetota
Alphaproteobacteria
Desulfobacterota
Verrucomicrobiota
Crenarchaeota
Cyanobacteriota
Chloroflexota
Nanoarchaeota
Acidobacteriota
Myxococcota
Patescibacteria
Firmicutes_A
Others

**Figure S4.** Prevalence (occupancy) and abundance of amplicon sequence variants (ASVs) from the seven most abundant orders within the permeable sediments based on 16S amplicon sequencing.

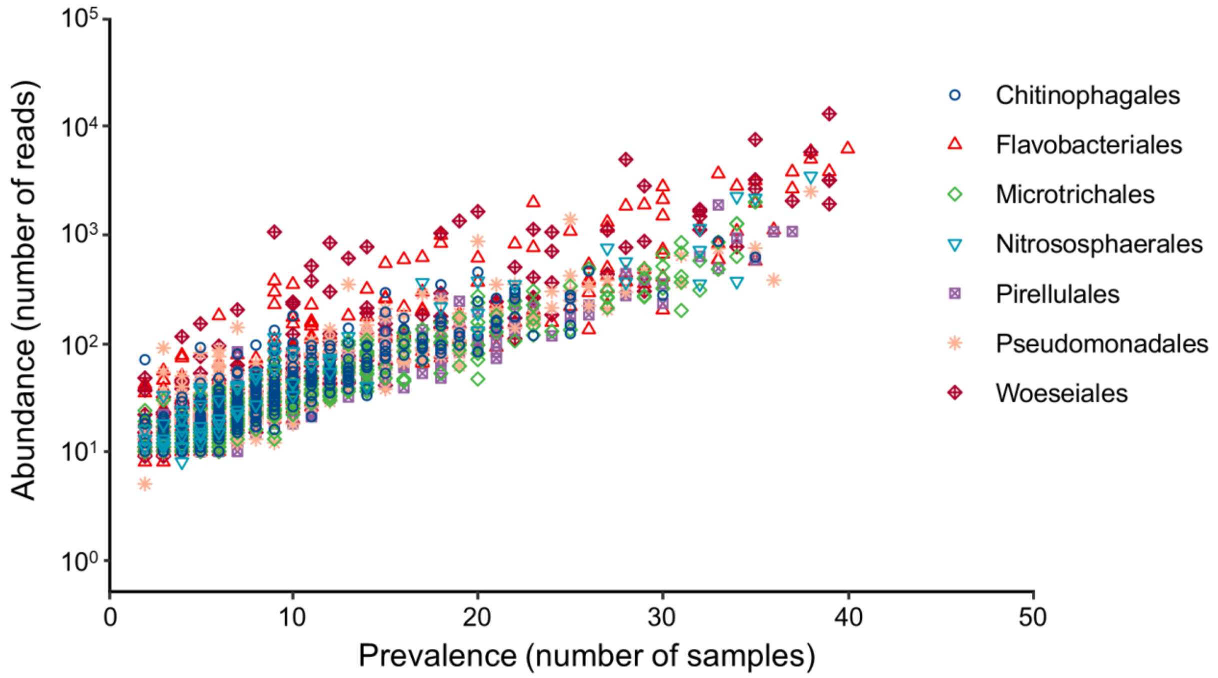

**Figure S5.** Community structure of permeable sediment metagenomes profiled using the single-copy ribosomal marker gene *rplP* in SingleM. The metagenomes are from sands collected at two tidal different zones (subtidal, intertidal), three different depths (shallow: 0-3 cm, intermediate: 14-17 cm, deep: 27-30 cm), and two different dates (A: 28/10/2016; C: 19/1/2017).

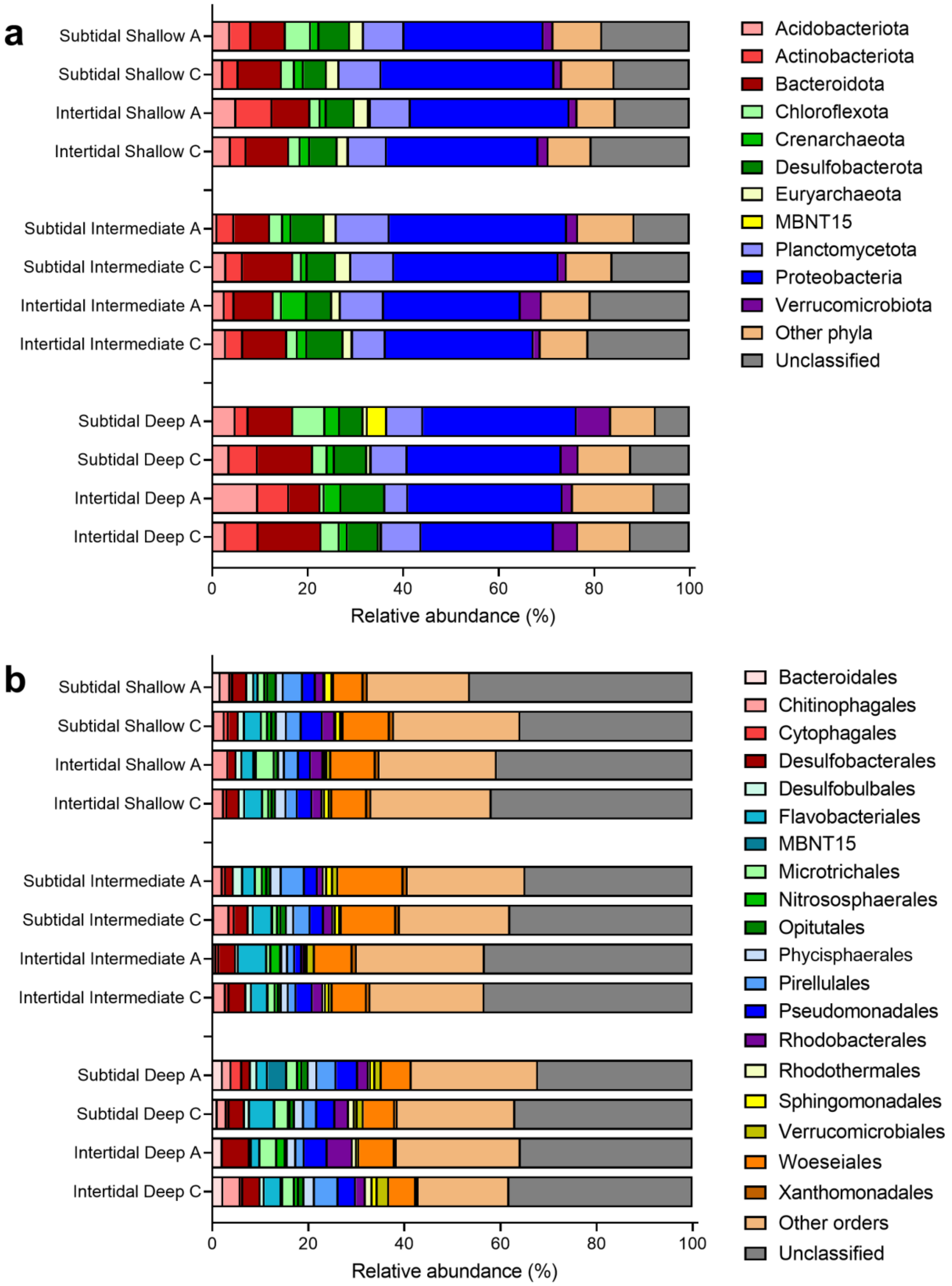

**Figure S6.** Maximum-likelihood tree of amino acid sequences of the sulfide-quinone oxidoreductase (Sqr), a marker for sulfide oxidation. The tree shows sequences from permeable sediment metagenome-assembled genomes (blue) alongside representative reference sequences (black). The subgroup of each reference sequence is denoted. The tree was constructed using the JTT matrix-based model, used all sites, and was bootstrapped with 50 replicates and midpoint-rooted.

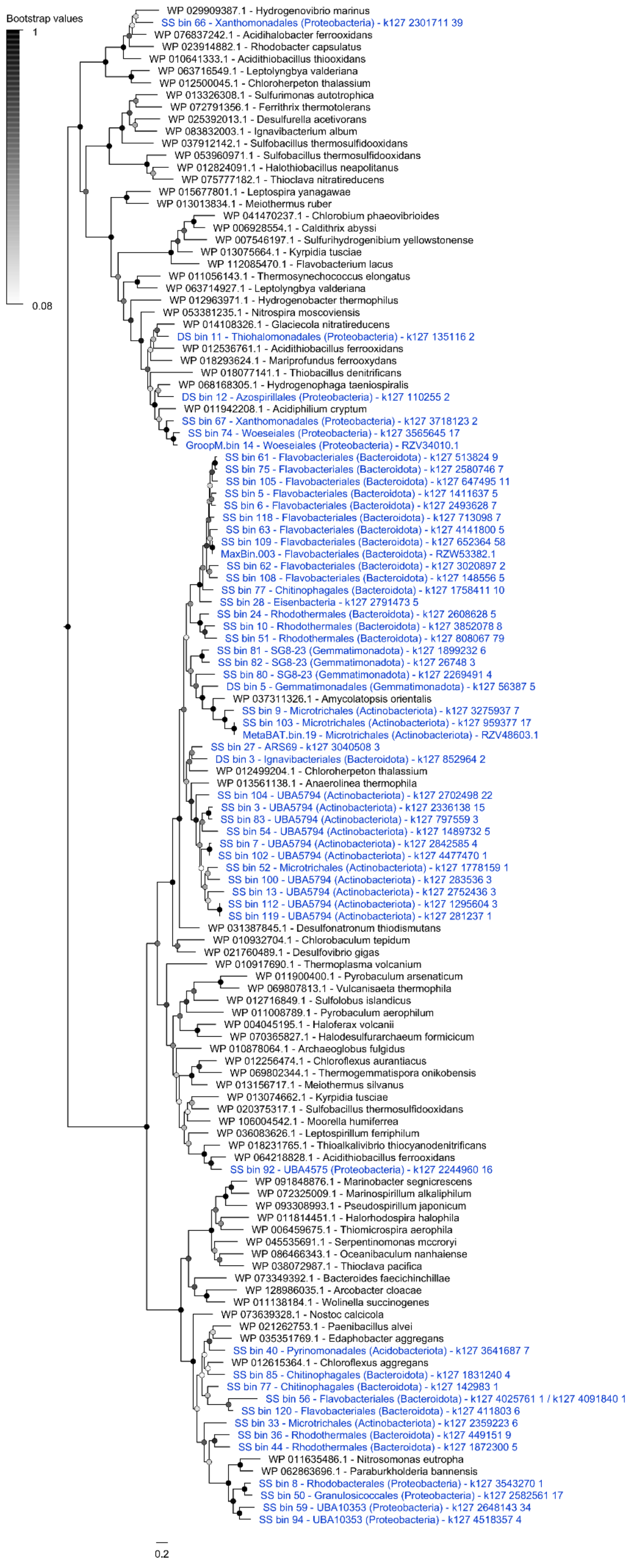

Bootstrap values
1
0.08
WP 029909387.1 - Hydrogenovibrio marinus
SS bin 66 - Xanthomonadales (Proteobacteria) - k127 2301711 39
WP 076837242.1 - Acidihalobacter ferrooxidans
WP 023914882.1 - Rhodobacter capsulatus
WP 010641333.1 - Acidithiobacillus thiooxidans
WP 063716549.1 - Leptolyngbya valderiana
WP 012500045.1 - Chloroherpeton thalassium
WP 013326308.1 - Sulfurimonas autotrophica
WP 072791356.1 - Ferrithrix thermotolerans
WP 025392013.1 - Desulfurella acetivorans
WP 083832003.1 - Ignavibacterium album
WP 037912142.1 - Sulfobacillus thermosulfidooxidans
WP 053960971.1 - Sulfobacillus thermosulfidooxidans
WP 012824091.1 - Halothiobacillus neapolitanus
WP 075777182.1 - Thioclava nitratireducens
WP 015677801.1 - Leptospira yanagawae
WP 013013834.1 - Meiothermus ruber
WP 041470237.1 - Chlorobium phaeovibrioides
WP 006928554.1 - Caldithrix abyssi
WP 007546197.1 - Sulfurihydrogenibium yellowstonense
WP 013075664.1 - Kyrpidia tusciae
WP 112085470.1 - Flavobacterium lacus
WP 011056143.1 - Thermosynechococcus elongatus
WP 063714927.1 - Leptolyngbya valderiana
WP 012963971.1 - Hydrogenobacter thermophilus
WP 053381235.1 - Nitrospira moscoviensis
WP 014108326.1 - Glaciecola nitratireducens
DS bin 11 - Thiohalomonadales (Proteobacteria) - k127 135116 2
WP 012536761.1 - Acidithiobacillus ferrooxidans
WP 018293624.1 - Mariprofundus ferrooxydans
WP 018077141.1 - Thiobacillus denitrificans
WP 068168305.1 - Hydrogenophaga taeniospiralis
DS bin 12 - Azospirillales (Proteobacteria) - k127 110255 2
WP 011942208.1 - Acidiphilium cryptum
SS bin 67 - Xanthomonadales (Proteobacteria) - k127 3718123 2
SS bin 74 - Woeseiales (Proteobacteria) - k127 3565645 17
GroopM.bin 14 - Woeseiales (Proteobacteria) - RZV34010.1
SS bin 61 - Flavobacteriales (Bacteroidota) - k127 513824 9
SS bin 75 - Flavobacteriales (Bacteroidota) - k127 2580746 7
SS bin 105 - Flavobacteriales (Bacteroidota) - k127 647495 11
SS bin 5 - Flavobacteriales (Bacteroidota) - k127 1411637 5
SS bin 6 - Flavobacteriales (Bacteroidota) - k127 2493628 7
SS bin 118 - Flavobacteriales (Bacteroidota) - k127 713098 7
SS bin 63 - Flavobacteriales (Bacteroidota) - k127 4141800 5
SS bin 109 - Flavobacteriales (Bacteroidota) - k127 652364 58
MaxBin.003 - Flavobacteriales (Bacteroidota) - RZW53382.1
SS bin 62 - Flavobacteriales (Bacteroidota) - k127 3020897 2
SS bin 108 - Flavobacteriales (Bacteroidota) - k127 148556 5
SS bin 77 - Chitinophagales (Bacteroidota) - k127 1758411 10
SS bin 28 - Eisenbacteria - k127 2791473 5
SS bin 24 - Rhodothermales (Bacteroidota) - k127 2608628 5
SS bin 10 - Rhodothermales (Bacteroidota) - k127 3852078 8
SS bin 51 - Rhodothermales (Bacteroidota) - k127 808067 79
SS bin 81 - SG8-23 (Gemmatimonadota) - k127 1899232 6
SS bin 82 - SG8-23 (Gemmatimonadota) - k127 26748 3
SS bin 80 - SG8-23 (Gemmatimonadota) - k127 2269491 4
DS bin 5 - Gemmatimonadales (Gemmatimonadota) - k127 56387 5
WP 037311326.1 - Amycolatopsis orientalis
SS bin 9 - Microtrichales (Actinobacteriota) - k127 3275937 7
SS bin 103 - Microtrichales (Actinobacteriota) - k127 959377 17
MetaBAT.bin.19 - Microtrichales (Actinobacteriota) - RZV48603.1
SS bin 27 - ARS69 - k127 3040508 3
DS bin 3 - Ignavibacteriales (Bacteroidota) - k127 852964 2
WP 012499204.1 - Chloroherpeton thalassium
WP 013561138.1 - Anaerolinea thermophila
SS bin 104 - UBA5794 (Actinobacteriota) - k127 2702498 22
SS bin 3 - UBA5794 (Actinobacteriota) - k127 2336138 15
SS bin 83 - UBA5794 (Actinobacteriota) - k127 797559 3
SS bin 54 - UBA5794 (Actinobacteriota) - k127 1489732 5
SS bin 7 - UBA5794 (Actinobacteriota) - k127 2842585 4
SS bin 102 - UBA5794 (Actinobacteriota) - k127 4477470 1
SS bin 52 - Microtrichales (Actinobacteriota) - k127 1778159 1
SS bin 100 - UBA5794 (Actinobacteriota) - k127 283536 3
SS bin 13 - UBA5794 (Actinobacteriota) - k127 2752436 3
SS bin 112 - UBA5794 (Actinobacteriota) - k127 1295604 3
SS bin 119 - UBA5794 (Actinobacteriota) - k127 281237 1
WP 031387845.1 - Desulfonatronum thiodismutans
WP 010932704.1 - Chlorobaculum tepidum
WP 021760489.1 - Desulfovibrio gigas
WP 010917690.1 - Thermoplasma volcanium
WP 011900400.1 - Pyrobaculum arsenaticum
WP 069807813.1 - Vulcanisaeta thermophila
WP 012716849.1 - Sulfolobus islandicus
WP 011008789.1 - Pyrobaculum aerophilum
WP 004045195.1 - Haloferax volcanii
WP 070365827.1 - Halodesulfurarchaeum formicicum
WP 010878064.1 - Archaeoglobus fulgidus
WP 012256474.1 - Chloroflexus aurantiacus
WP 069802344.1 - Thermogemmatispora onikobensis
WP 013156717.1 - Meiothermus silvanus
WP 013074662.1 - Kyrpidia tusciae
WP 020375317.1 - Sulfobacillus thermosulfidooxidans
WP 106004542.1 - Moorella humiferrea
WP 036083626.1 - Leptospirillum ferriphilum
WP 018231765.1 - Thioalkalivibrio thiocyanodenitrificans
WP 064218828.1 - Acidithiobacillus ferrooxidans
SS bin 92 - UBA4575 (Proteobacteria) - k127 2244960 16
WP 091848876.1 - Marinobacter segnicrescens
WP 072325009.1 - Marinospirillum alkaliphilum
WP 093308993.1 - Pseudospirillum japonicum
WP 011814451.1 - Halorhodospira halophila
WP 006459675.1 - Thiomicrospira aerophila
WP 045535691.1 - Serpentinomonas mccroryi
WP 086466343.1 - Oceanibaculum nanhaiense
WP 038072987.1 - Thioclava pacifica
WP 073349392.1 - Bacteroides faecichinchillae
WP 128986035.1 - Arcobacter cloacae
WP 011138184.1 - Wolinella succinogenes
WP 073639328.1 - Nostoc calcicola
WP 021262753.1 - Paenibacillus alvei
WP 035351769.1 - Edaphobacter aggregans
SS bin 40 - Pyrinomonadales (Acidobacteriota) - k127 3641687 7
WP 012615364.1 - Chloroflexus aggregans
SS bin 85 - Chitinophagales (Bacteroidota) - k127 1831240 4
SS bin 77 - Chitinophagales (Bacteroidota) - k127 142983 1
SS bin 56 - Flavobacteriales (Bacteroidota) - k127 4025761 1 / k127 4091840 1
SS bin 120 - Flavobacteriales (Bacteroidota) - k127 411803 6
SS bin 33 - Microtrichales (Actinobacteriota) - k127 2359223 6
SS bin 36 - Rhodothermales (Bacteroidota) - k127 449151 9
SS bin 44 - Rhodothermales (Bacteroidota) - k127 1872300 5
WP 011635486.1 - Nitrosomonas eutropha
WP 062863696.1 - Paraburkholderia bannensis
SS bin 8 - Rhodobacterales (Proteobacteria) - k127 3543270 1
SS bin 50 - Granulosicoccales (Proteobacteria) - k127 2582561 17
SS bin 59 - UBA10353 (Proteobacteria) - k127 2648143 34
SS bin 94 - UBA10353 (Proteobacteria) - k127 4518357 4
0.2

**Figure S7.** Maximum-likelihood tree of amino acid sequences of flavocytochrome *c* sulfide dehydrogenase (FCC), a marker for sulfide oxidation. The tree shows sequences from permeable sediment metagenome-assembled genomes (blue) alongside representative reference sequences (black). The tree was constructed using the JTT matrix-based model, used all sites, and was bootstrapped with 50 replicates and midpoint-rooted.

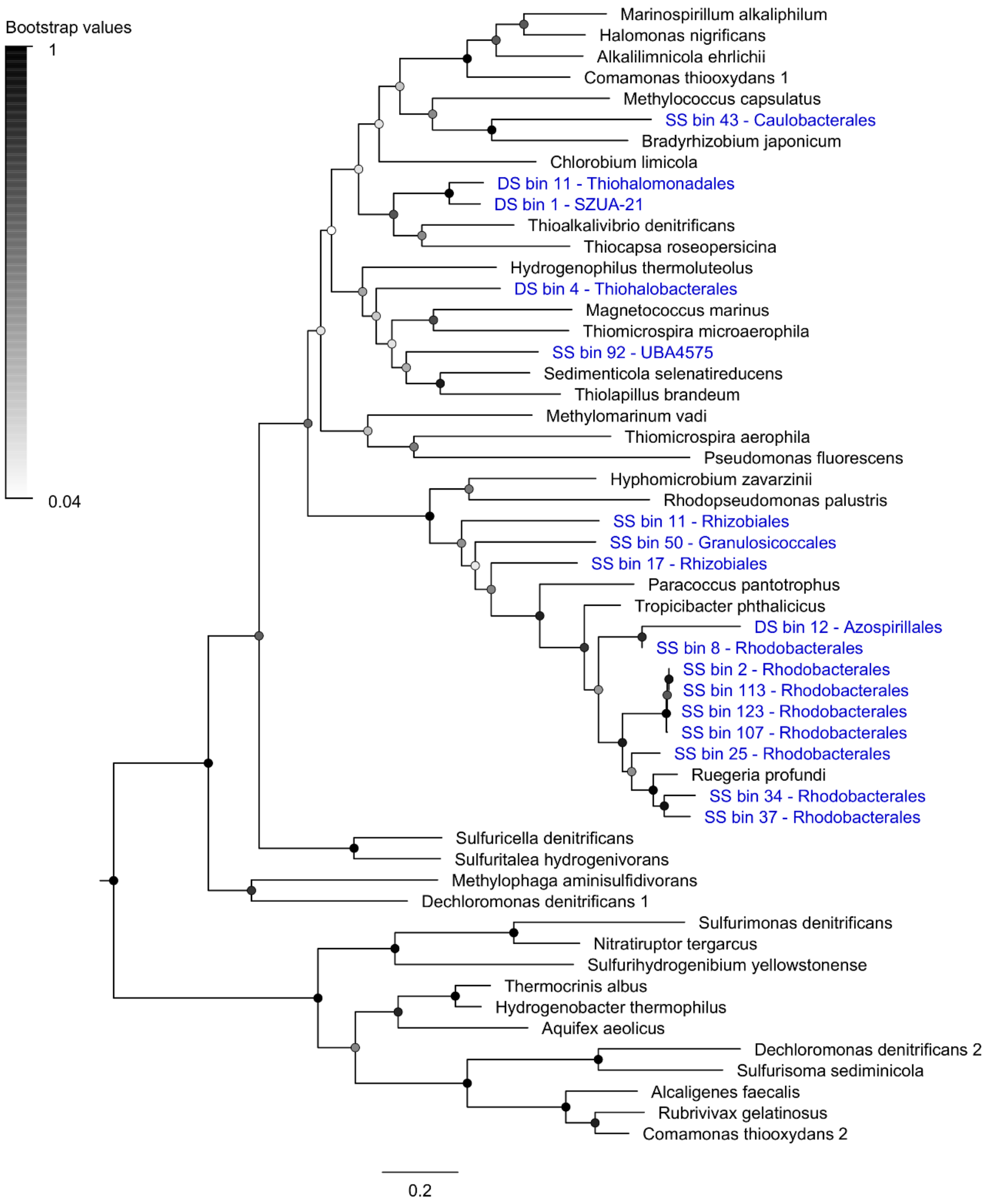

**Figure S8.** Maximum-likelihood tree of amino acid sequences of dissimilatory sulfite reductase A subunit (DsrA). The tree shows sequences from permeable sediment metagenome-assembled genomes (blue) alongside representative reference sequences (black). This enzyme is a marker for dissimilatory sulfite reduction (middle and bottom major clades; Desulfobacterota bins) and sulfide oxidation (top clade, r-DsrA; Proteobacteria bins). The tree was constructed using the JTT matrix-based model, used all sites, and was bootstrapped with 50 replicates and midpoint-rooted.

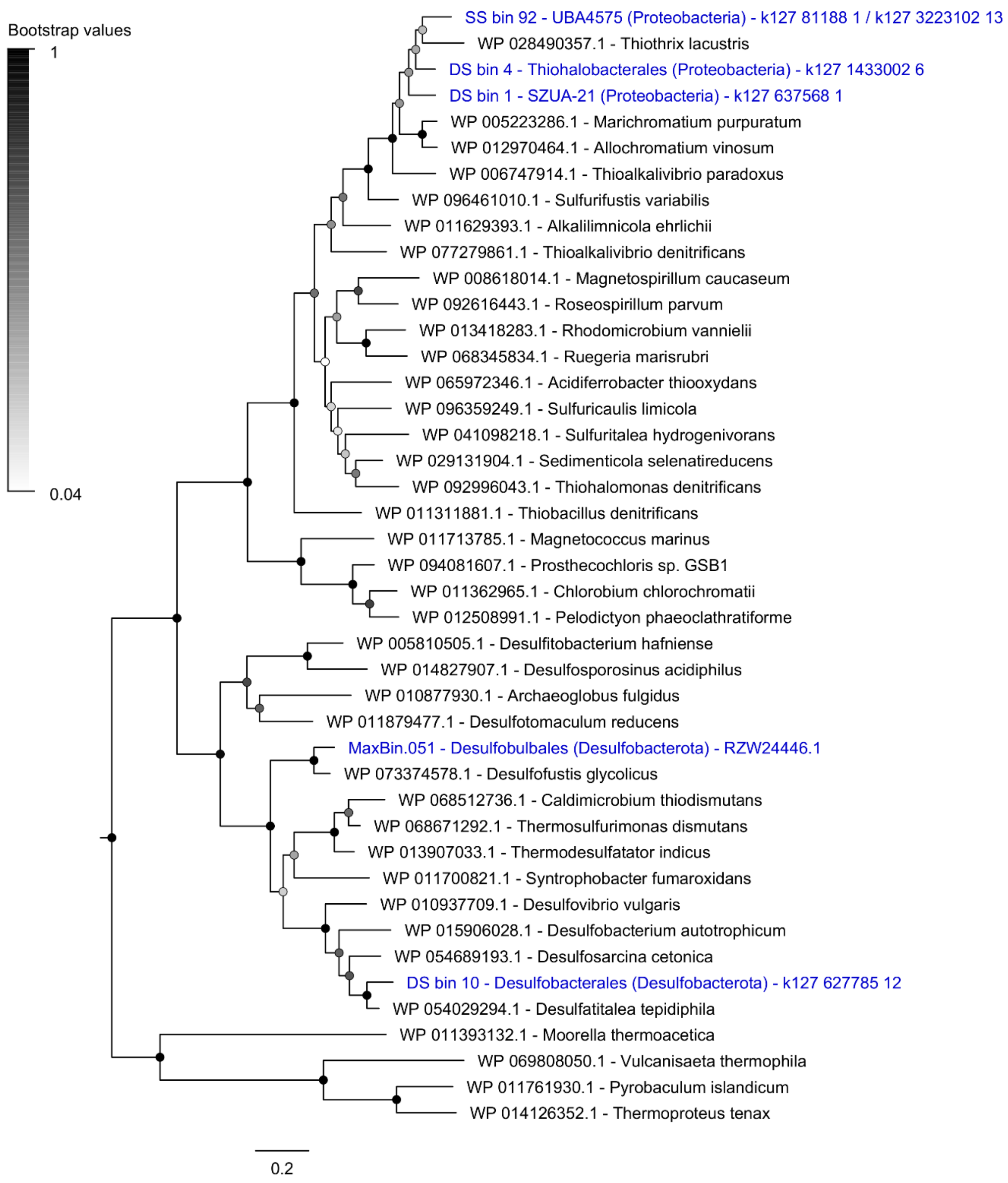

**Figure S9.** Maximum-likelihood tree of amino acid sequences of the thiosulfohydrolase (SoxB), a marker for thiosulfate oxidation. The tree shows sequences from permeable sediment metagenome-assembled genomes (blue) alongside representative reference sequences (black). The subgroup of each reference sequence is denoted. The tree was constructed using the JTT matrix-based model, used all sites, and was bootstrapped with 50 replicates and midpoint-rooted.

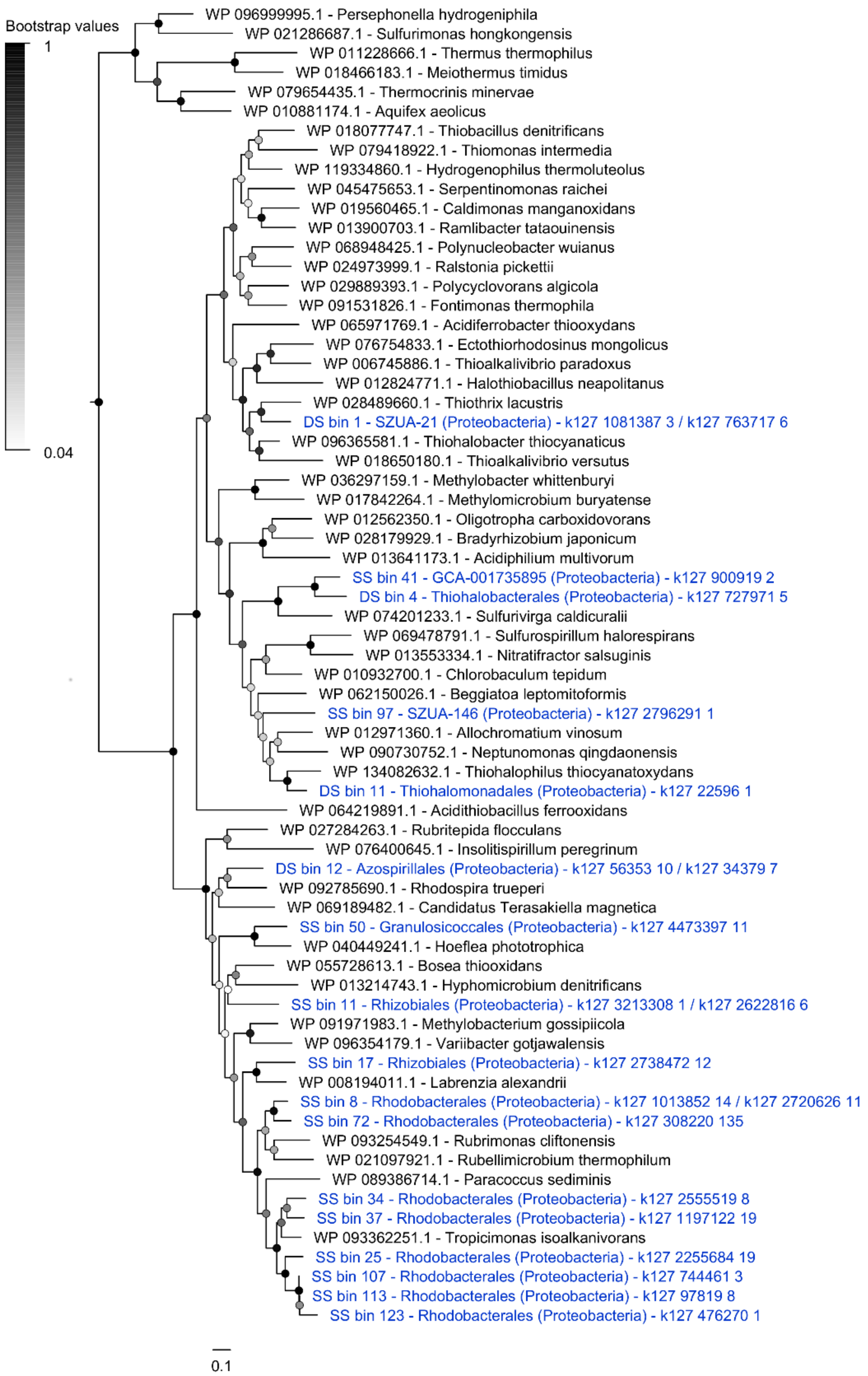

**Figure S10.** Maximum-likelihood tree of amino acid sequences of carbon monoxide dehydrogenase large subunit (CoxL), a marker for aerobic carbon monoxide oxidation. The tree shows sequences from permeable sediment metagenome-assembled genomes (blue) alongside representative reference sequences (black). The tree was constructed using the JTT matrix-based model, used all sites, and was bootstrapped with 50 replicates and midpoint-rooted.

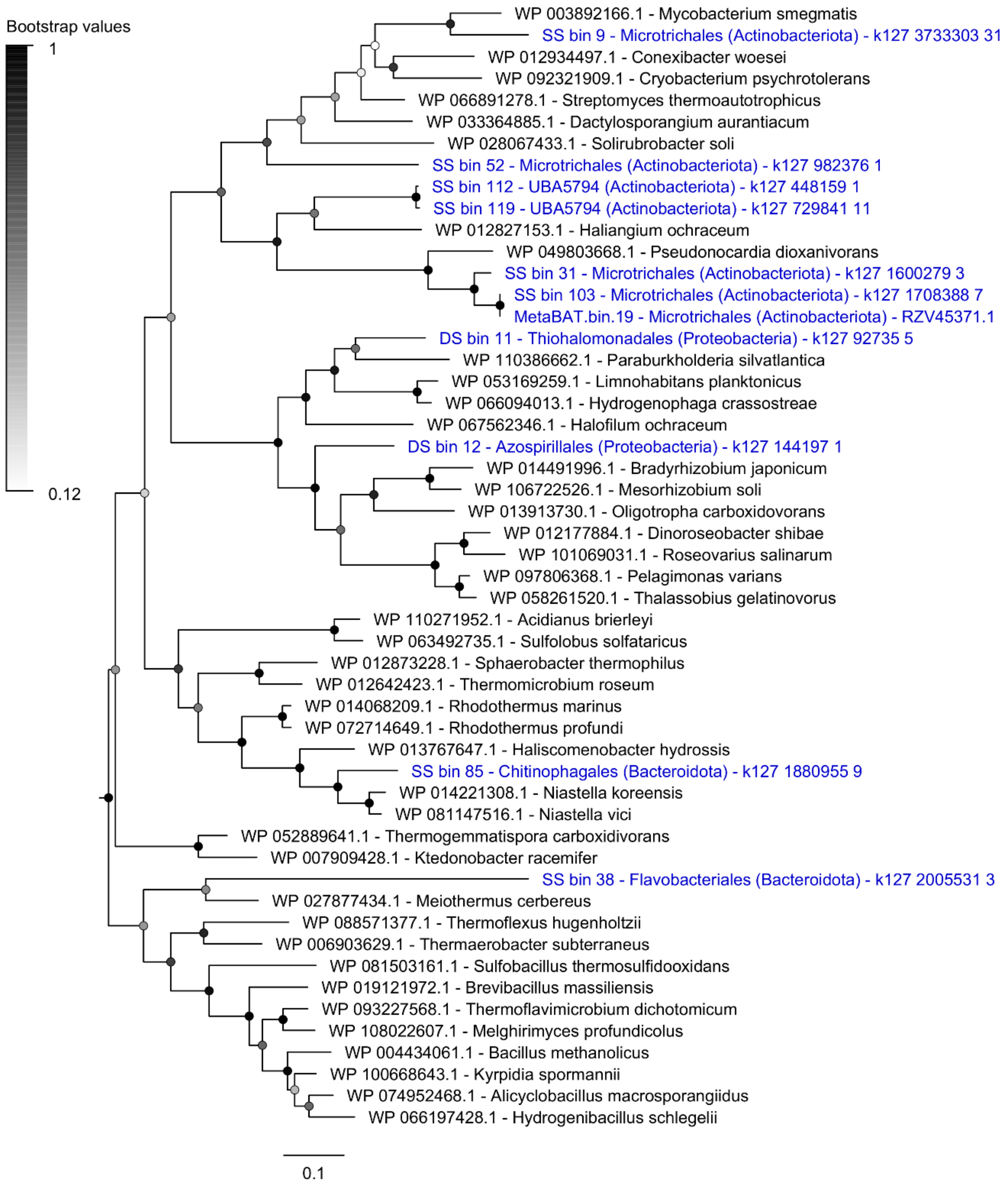

**Figure S11.** Maximum-likelihood tree of amino acid sequences of group 1 [NiFe]-hydrogenase large subunits, a marker for hydrogen oxidation during respiratory processes. The tree shows sequences from permeable sediment metagenome-assembled genomes (blue) alongside representative reference sequences (black). The subgroup of each reference sequence is denoted according to the HydDB classification scheme. The tree was constructed using the JTT matrix-based model, used all sites, and was bootstrapped with 50 replicates and midpoint-rooted.

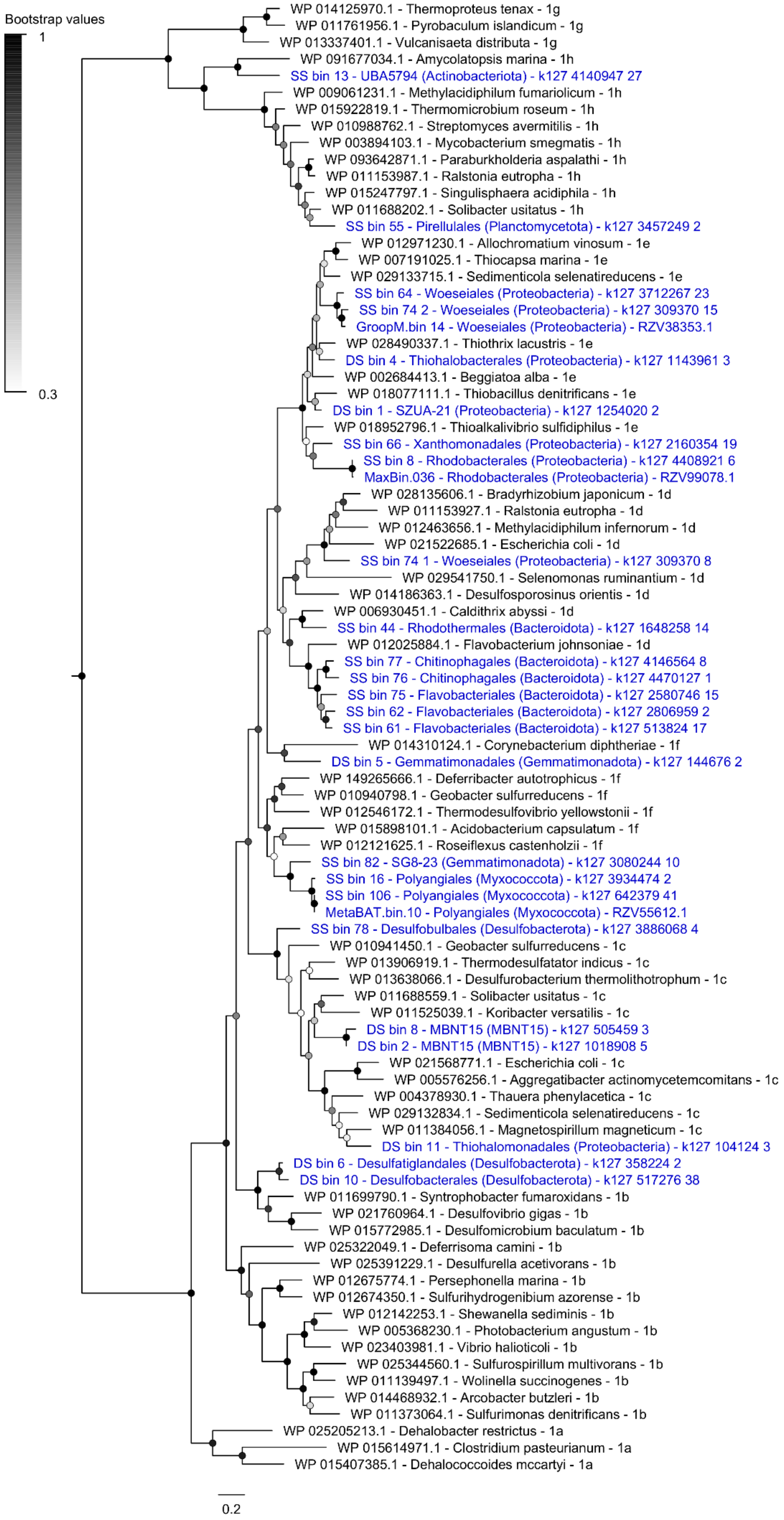

Bootstrap values
1
0.3
WP 014125970.1 - Thermoproteus tenax - 1g
WP 011761956.1 - Pyrobaculum islandicum - 1g
WP 013337401.1 - Vulcanisaeta distributa - 1g
WP 091677034.1 - Amycolatopsis marina - 1h
SS bin 13 - UBA5794 (Actinobacteriota) - k127 4140947 27
WP 009061231.1 - Methylacidiphilum fumariolicum - 1h
WP 015922819.1 - Thermomicrobium roseum - 1h
WP 010988762.1 - Streptomyces avermitilis - 1h
WP 003894103.1 - Mycobacterium smegmatis - 1h
WP 093642871.1 - Paraburkholderia aspalathi - 1h
WP 011153987.1 - Ralstonia eutropha - 1h
WP 015247797.1 - Singulisphaera acidiphila - 1h
WP 011688202.1 - Solibacter usitatus - 1h
SS bin 55 - Pirellulales (Planctomycetota) - k127 3457249 2
WP 012971230.1 - Allochromatium vinosum - 1e
WP 007191025.1 - Thiocapsa marina - 1e
WP 029133715.1 - Sedimenticola selenatireducens - 1e
SS bin 64 - Woeseiales (Proteobacteria) - k127 3712267 23
SS bin 74 2 - Woeseiales (Proteobacteria) - k127 309370 15
GroopM.bin 14 - Woeseiales (Proteobacteria) - RZV38353.1
WP 028490337.1 - Thiothrix lacustris - 1e
DS bin 4 - Thiohalobacterales (Proteobacteria) - k127 1143961 3
WP 002684413.1 - Beggiatoa alba - 1e
WP 018077111.1 - Thiobacillus denitrificans - 1e
DS bin 1 - SZUA-21 (Proteobacteria) - k127 1254020 2
WP 018952796.1 - Thioalkalivibrio sulfidiphilus - 1e
SS bin 66 - Xanthomonadales (Proteobacteria) - k127 2160354 19
SS bin 8 - Rhodobacterales (Proteobacteria) - k127 4408921 6
MaxBin.036 - Rhodobacterales (Proteobacteria) - RZV99078.1
WP 028135606.1 - Bradyrhizobium japonicum - 1d
WP 011153927.1 - Ralstonia eutropha - 1d
WP 012463656.1 - Methylacidiphilum infernorum - 1d
WP 021522685.1 - Escherichia coli - 1d
SS bin 74 1 - Woeseiales (Proteobacteria) - k127 309370 8
WP 029541750.1 - Selenomonas ruminantium - 1d
WP 014186363.1 - Desulfosporosinus orientis - 1d
WP 006930451.1 - Caldithrix abyssi - 1d
SS bin 44 - Rhodothermales (Bacteroidota) - k127 1648258 14
WP 012025884.1 - Flavobacterium johnsoniae - 1d
SS bin 77 - Chitinophagales (Bacteroidota) - k127 4146564 8
SS bin 76 - Chitinophagales (Bacteroidota) - k127 4470127 1
SS bin 75 - Flavobacteriales (Bacteroidota) - k127 2580746 15
SS bin 62 - Flavobacteriales (Bacteroidota) - k127 2806959 2
SS bin 61 - Flavobacteriales (Bacteroidota) - k127 513824 17
WP 014310124.1 - Corynebacterium diphtheriae - 1f
DS bin 5 - Gemmatimonadales (Gemmatimonadota) - k127 144676 2
WP 149265666.1 - Deferribacter autotrophicus - 1f
WP 010940798.1 - Geobacter sulfurreducens - 1f
WP 012546172.1 - Thermodesulfovibrio yellowstonii - 1f
WP 015898101.1 - Acidobacterium capsulatum - 1f
WP 012121625.1 - Roseiflexus castenholzii - 1f
SS bin 82 - SG8-23 (Gemmatimonadota) - k127 3080244 10
SS bin 16 - Polyangiales (Myxococcota) - k127 3934474 2
SS bin 106 - Polyangiales (Myxococcota) - k127 642379 41
MetaBAT.bin.10 - Polyangiales (Myxococcota) - RZV55612.1
SS bin 78 - Desulfobulbales (Desulfobacterota) - k127 3886068 4
WP 010941450.1 - Geobacter sulfurreducens - 1c
WP 013906919.1 - Thermodesulfatator indicus - 1c
WP 013638066.1 - Desulfurobacterium thermolithotrophum - 1c
WP 011688559.1 - Solibacter usitatus - 1c
WP 011525039.1 - Koribacter versatilis - 1c
DS bin 8 - MBNT15 (MBNT15) - k127 505459 3
DS bin 2 - MBNT15 (MBNT15) - k127 1018908 5
WP 021568771.1 - Escherichia coli - 1c
WP 005576256.1 - Aggregatibacter actinomycetemcomitans - 1c
WP 004378930.1 - Thauera phenylacetica - 1c
WP 029132834.1 - Sedimenticola selenatireducens - 1c
WP 011384056.1 - Magnetospirillum magneticum - 1c
DS bin 11 - Thiohalomonadales (Proteobacteria) - k127 104124 3
DS bin 6 - Desulfatiglandales (Desulfobacterota) - k127 358224 2
DS bin 10 - Desulfobacterales (Desulfobacterota) - k127 517276 38
WP 011699790.1 - Syntrophobacter fumaroxidans - 1b
WP 021760964.1 - Desulfovibrio gigas - 1b
WP 015772985.1 - Desulfomicrobium baculatum - 1b
WP 025322049.1 - Deferrisoma camini - 1b
WP 025391229.1 - Desulfurella acetivorans - 1b
WP 012675774.1 - Persephonella marina - 1b
WP 012674350.1 - Sulfurihydrogenibium azorense - 1b
WP 012142253.1 - Shewanella sediminis - 1b
WP 005368230.1 - Photobacterium angustum - 1b
WP 023403981.1 - Vibrio halioticoli - 1b
WP 025344560.1 - Sulfurospirillum multivorans - 1b
WP 011139497.1 - Wolinella succinogenes - 1b
WP 014468932.1 - Arcobacter butzleri - 1b
WP 011373064.1 - Sulfurimonas denitrificans - 1b
WP 025205213.1 - Dehalobacter restrictus - 1a
WP 015614971.1 - Clostridium pasteurianum - 1a
WP 015407385.1 - Dehalococcoides mccartyi - 1a
0.2

**Figure S12.** Maximum-likelihood tree of amino acid sequences of dissimilatory nitrate reductase G subunit (NarG), a marker for denitrification and dissimilatory nitrate reduction to ammonium. The tree shows sequences from permeable sediment metagenome-assembled genomes (blue) alongside representative reference sequences (black). The tree was constructed using the JTT matrix-based model, used all sites, and was bootstrapped with 50 replicates and midpoint-rooted.

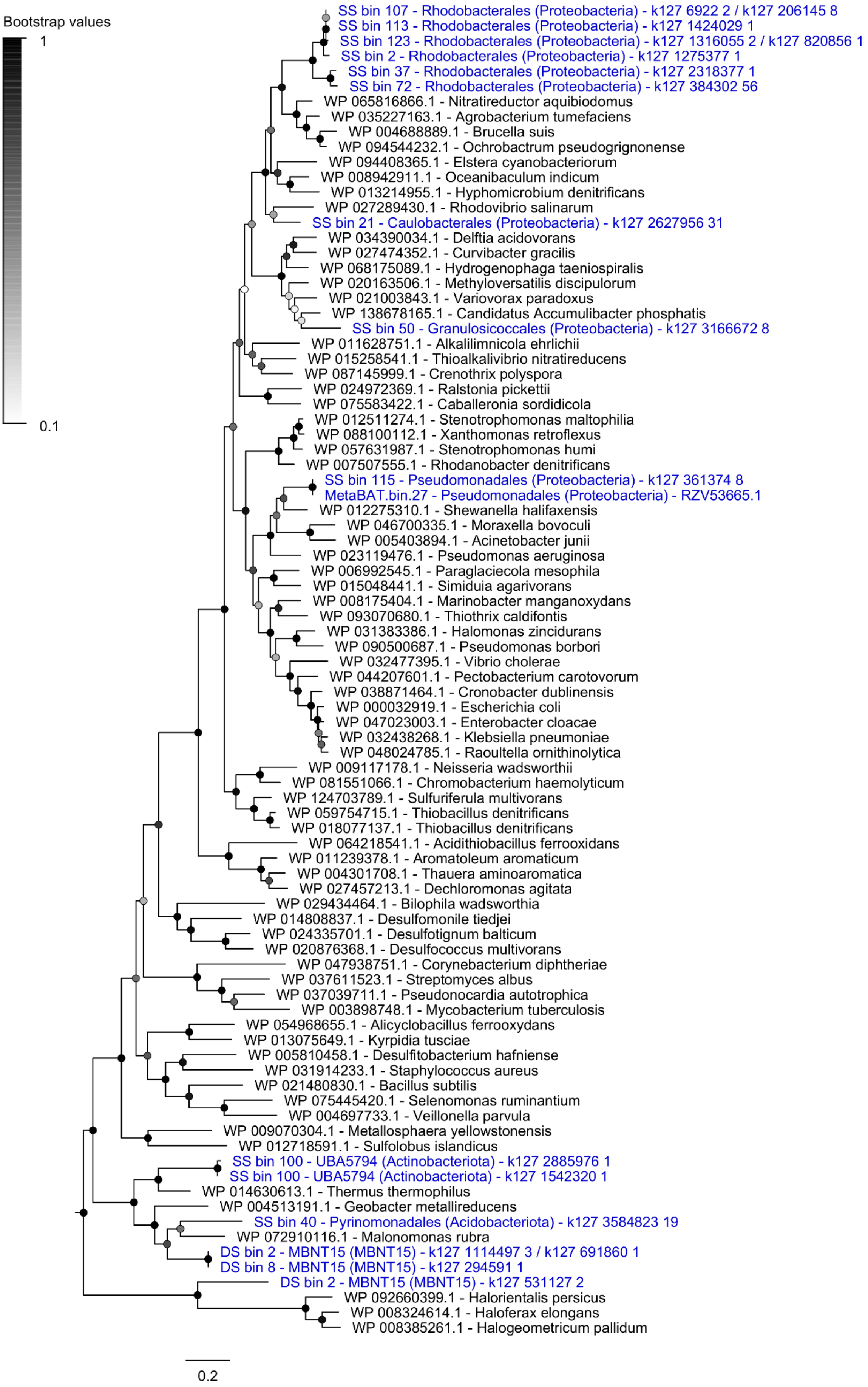
Bootstrap values
1
0.1
SS bin 107 - Rhodobacterales (Proteobacteria) - k127 6922 2 / k127 206145 8
SS bin 113 - Rhodobacterales (Proteobacteria) - k127 1424029 1
SS bin 123 - Rhodobacterales (Proteobacteria) - k127 1316055 2 / k127 820856 1
SS bin 2 - Rhodobacterales (Proteobacteria) - k127 1275377 1
SS bin 37 - Rhodobacterales (Proteobacteria) - k127 2318377 1
SS bin 72 - Rhodobacterales (Proteobacteria) - k127 384302 56
WP 065816866.1 - Nitratireductor aquibiodomus
WP 035227163.1 - Agrobacterium tumefaciens
WP 004688889.1 - Brucella suis
WP 094544232.1 - Ochrobactrum pseudogrignonense
WP 094408365.1 - Elstera cyanobacteriorum
WP 008942911.1 - Oceanibaculum indicum
WP 013214955.1 - Hyphomicrobium denitrificans
WP 027289430.1 - Rhodovibrio salinarum
SS bin 21 - Caulobacterales (Proteobacteria) - k127 2627956 31
WP 034390034.1 - Delftia acidovorans
WP 027474352.1 - Curvibacter gracilis
WP 068175089.1 - Hydrogenophaga taeniospiralis
WP 020163506.1 - Methyloversatilis discipulorum
WP 021003843.1 - Variovorax paradoxus
WP 138678165.1 - Candidatus Accumulibacter phosphatis
SS bin 50 - Granulosicoccales (Proteobacteria) - k127 3166672 8
WP 011628751.1 - Alkalilimnicola ehrlichii
WP 015258541.1 - Thioalkalivibrio nitratireducens
WP 087145999.1 - Crenothrix polyspora
WP 024972369.1 - Ralstonia pickettii
WP 075583422.1 - Caballeronia sordidicola
WP 012511274.1 - Stenotrophomonas maltophilia
WP 088100112.1 - Xanthomonas retroflexus
WP 057631987.1 - Stenotrophomonas humi
WP 007507555.1 - Rhodanobacter denitrificans
SS bin 115 - Pseudomonadales (Proteobacteria) - k127 361374 8
MetaBAT.bin.27 - Pseudomonadales (Proteobacteria) - RZV53665.1
WP 012275310.1 - Shewanella halifaxensis
WP 046700335.1 - Moraxella bovoculi
WP 005403894.1 - Acinetobacter junii
WP 023119476.1 - Pseudomonas aeruginosa
WP 006992545.1 - Paraglaciecola mesophila
WP 015048441.1 - Simiduia agarivorans
WP 008175404.1 - Marinobacter manganoxydans
WP 093070680.1 - Thiothrix caldifontis
WP 031383386.1 - Halomonas zincidurans
WP 090500687.1 - Pseudomonas borbori
WP 032477395.1 - Vibrio cholerae
WP 044207601.1 - Pectobacterium carotovorum
WP 038871464.1 - Cronobacter dublinensis
WP 000032919.1 - Escherichia coli
WP 047023003.1 - Enterobacter cloacae
WP 032438268.1 - Klebsiella pneumoniae
WP 048024785.1 - Raoultella ornithinolytica
WP 009117178.1 - Neisseria wadsworthii
WP 081551066.1 - Chromobacterium haemolyticum
WP 124703789.1 - Sulfuriferula multivorans
WP 059754715.1 - Thiobacillus denitrificans
WP 018077137.1 - Thiobacillus denitrificans
WP 064218541.1 - Acidithiobacillus ferrooxidans
WP 011239378.1 - Aromatoleum aromaticum
WP 004301708.1 - Thauera aminoaromatica
WP 027457213.1 - Dechloromonas agitata
WP 029434464.1 - Bilophila wadsworthia
WP 014808837.1 - Desulfomonile tiedjei
WP 024335701.1 - Desulfotignum balticum
WP 020876368.1 - Desulfococcus multivorans
WP 047938751.1 - Corynebacterium diphtheriae
WP 037611523.1 - Streptomyces albus
WP 037039711.1 - Pseudonocardia autotrophica
WP 003898748.1 - Mycobacterium tuberculosis
WP 054968655.1 - Alicyclobacillus ferrooxydans
WP 013075649.1 - Kyrpidia tusciae
WP 005810458.1 - Desulfitobacterium hafniense
WP 031914233.1 - Staphylococcus aureus
WP 021480830.1 - Bacillus subtilis
WP 075445420.1 - Selenomonas ruminantium
WP 004697733.1 - Veillonella parvula
WP 009070304.1 - Metallosphaera yellowstonensis
WP 012718591.1 - Sulfolobus islandicus
SS bin 100 - UBA5794 (Actinobacteriota) - k127 2885976 1
SS bin 100 - UBA5794 (Actinobacteriota) - k127 1542320 1
WP 014630613.1 - Thermus thermophilus
WP 004513191.1 - Geobacter metallireducens
SS bin 40 - Pyrinomonadales (Acidobacteriota) - k127 3584823 19
WP 072910116.1 - Malonomonas rubra
DS bin 2 - MBNT15 (MBNT15) - k127 1114497 3 / k127 691860 1
DS bin 8 - MBNT15 (MBNT15) - k127 294591 1
DS bin 2 - MBNT15 (MBNT15) - k127 531127 2
WP 092660399.1 - Halorientalis persicus
WP 008324614.1 - Haloferax elongans
WP 008385261.1 - Halogeometricum pallidum
0.2

**Figure S13.** Maximum-likelihood tree of amino acid sequences of periplasmic nitrate reductase A subunit (NapA), a marker for denitrification and dissimilatory nitrate reduction to ammonium. The tree shows sequences from permeable sediment metagenome-assembled genomes (blue) alongside representative reference sequences (black). The tree was constructed using the JTT matrix-based model, used all sites, and was bootstrapped with 50 replicates and midpoint-rooted.

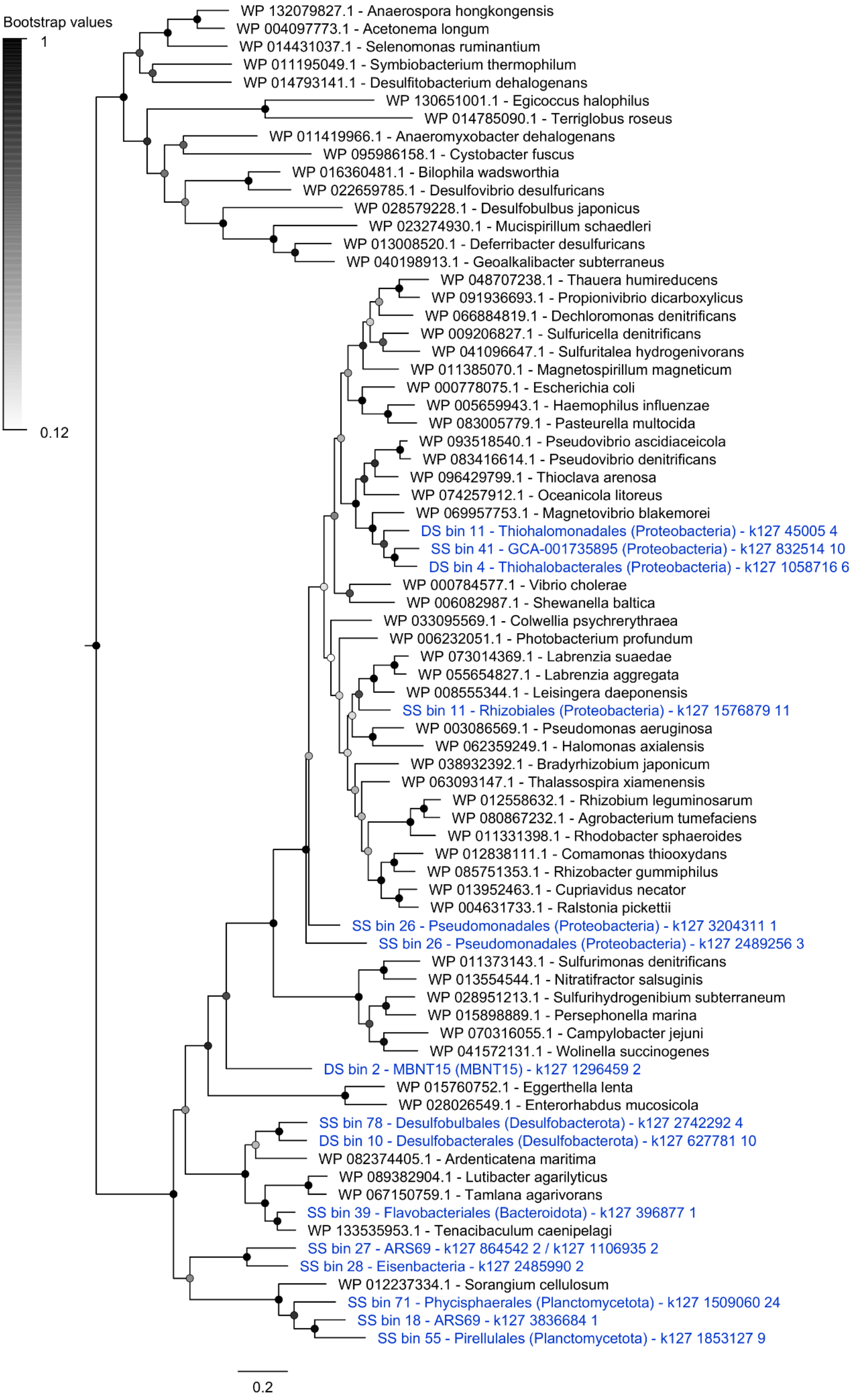
Bootstrap values
1
0.12
WP 132079827.1 - Anaerospora hongkongensis
WP 004097773.1 - Acetonema longum
WP 014431037.1 - Selenomonas ruminantium
WP 011195049.1 - Symbiobacterium thermophilum
WP 014793141.1 - Desulfitobacterium dehalogenans
WP 130651001.1 - Egicoccus halophilus
WP 014785090.1 - Terriglobus roseus
WP 011419966.1 - Anaeromyxobacter dehalogenans
WP 095986158.1 - Cystobacter fuscus
WP 016360481.1 - Bilophila wadsworthia
WP 022659785.1 - Desulfovibrio desulfuricans
WP 028579228.1 - Desulfobulbus japonicus
WP 023274930.1 - Mucispirillum schaedleri
WP 013008520.1 - Deferribacter desulfuricans
WP 040198913.1 - Geoalkalibacter subterraneus
WP 048707238.1 - Thauera humireducens
WP 091936693.1 - Propionivibrio dicarboxylicus
WP 066884819.1 - Dechloromonas denitrificans
WP 009206827.1 - Sulfuricella denitrificans
WP 041096647.1 - Sulfuritalea hydrogenivorans
WP 011385070.1 - Magnetospirillum magneticum
WP 000778075.1 - Escherichia coli
WP 005659943.1 - Haemophilus influenzae
WP 083005779.1 - Pasteurella multocida
WP 093518540.1 - Pseudovibrio ascidiaceicola
WP 083416614.1 - Pseudovibrio denitrificans
WP 096429799.1 - Thioclava arenosa
WP 074257912.1 - Oceanicola litoreus
WP 069957753.1 - Magnetovibrio blakemorei
DS bin 11 - Thiohalomonadales (Proteobacteria) - k127 45005 4
SS bin 41 - GCA-001735895 (Proteobacteria) - k127 832514 10
DS bin 4 - Thiohalobacterales (Proteobacteria) - k127 1058716 6
WP 000784577.1 - Vibrio cholerae
WP 006082987.1 - Shewanella baltica
WP 033095569.1 - Colwellia psychrerythraea
WP 006232051.1 - Photobacterium profundum
WP 073014369.1 - Labrenzia suaedae
WP 055654827.1 - Labrenzia aggregata
WP 008555344.1 - Leisingera daeponensis
SS bin 11 - Rhizobiales (Proteobacteria) - k127 1576879 11
WP 003086569.1 - Pseudomonas aeruginosa
WP 062359249.1 - Halomonas axialensis
WP 038932392.1 - Bradyrhizobium japonicum
WP 063093147.1 - Thalassospira xiamenensis
WP 012558632.1 - Rhizobium leguminosarum
WP 080867232.1 - Agrobacterium tumefaciens
WP 011331398.1 - Rhodobacter sphaeroides
WP 012838111.1 - Comamonas thiooxydans
WP 085751353.1 - Rhizobacter gummiphilus
WP 013952463.1 - Cupriavidus necator
WP 004631733.1 - Ralstonia pickettii
SS bin 26 - Pseudomonadales (Proteobacteria) - k127 3204311 1
SS bin 26 - Pseudomonadales (Proteobacteria) - k127 2489256 3
WP 011373143.1 - Sulfurimonas denitrificans
WP 013554544.1 - Nitratifractor salsuginis
WP 028951213.1 - Sulfurihydrogenibium subterraneum
WP 015898889.1 - Persephonella marina
WP 070316055.1 - Campylobacter jejuni
WP 041572131.1 - Wolinella succinogenes
DS bin 2 - MBNT15 (MBNT15) - k127 1296459 2
WP 015760752.1 - Eggerthella lenta
WP 028026549.1 - Enterorhabdus mucosicola
SS bin 78 - Desulfobulbales (Desulfobacterota) - k127 2742292 4
DS bin 10 - Desulfobacterales (Desulfobacterota) - k127 627781 10
WP 082374405.1 - Ardenticatena maritima
WP 089382904.1 - Lutibacter agarilyticus
WP 067150759.1 - Tamlana agarivorans
SS bin 39 - Flavobacteriales (Bacteroidota) - k127 396877 1
WP 133535953.1 - Tenacibaculum caenipelagi
SS bin 27 - ARS69 - k127 864542 2 / k127 1106935 2
SS bin 28 - Eisenbacteria - k127 2485990 2
WP 012237334.1 - Sorangium cellulosum
SS bin 71 - Phycisphaerales (Planctomycetota) - k127 1509060 24
SS bin 18 - ARS69 - k127 3836684 1
SS bin 55 - Pirellulales (Planctomycetota) - k127 1853127 9
0.2

**Figure S14.** Maximum-likelihood tree of amino acid sequences of copper-containing nitrite reductase K subunit (NirK), a marker for denitrification. The tree shows sequences from permeable sediment metagenome-assembled genomes (blue) alongside representative reference sequences (black). The tree was constructed using the JTT matrix-based model, used all sites, and was bootstrapped with 50 replicates and midpoint-rooted.

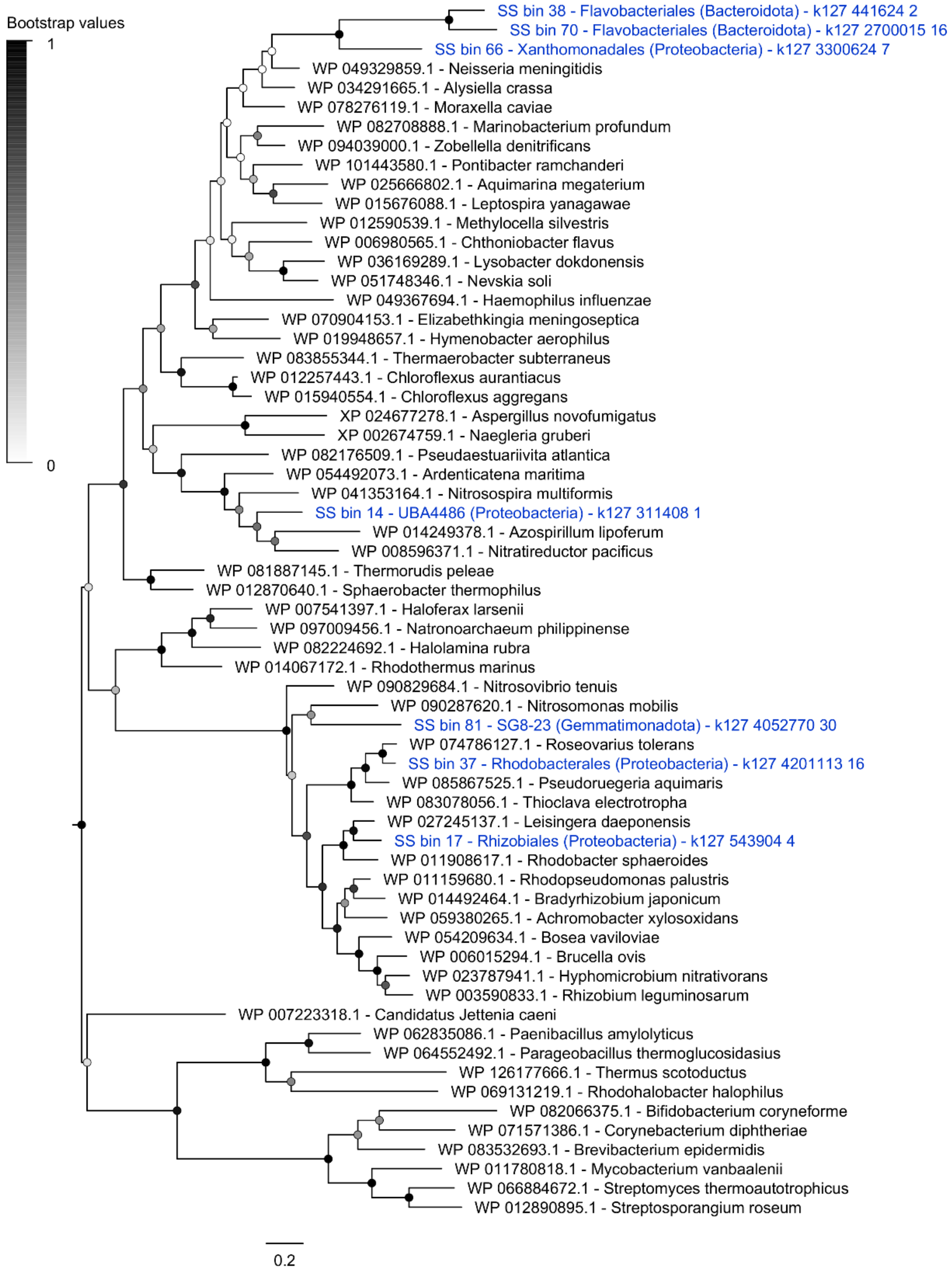

**Figure S15.** Maximum-likelihood tree of amino acid sequences of cytochrome $cd_1$ nitrite reductase S subunit (NirS), a marker for denitrification. The tree shows sequences from permeable sediment metagenome-assembled genomes (blue) alongside representative reference sequences (black). The tree was constructed using the JTT matrix-based model, used all sites, and was bootstrapped with 50 replicates and midpoint-rooted.

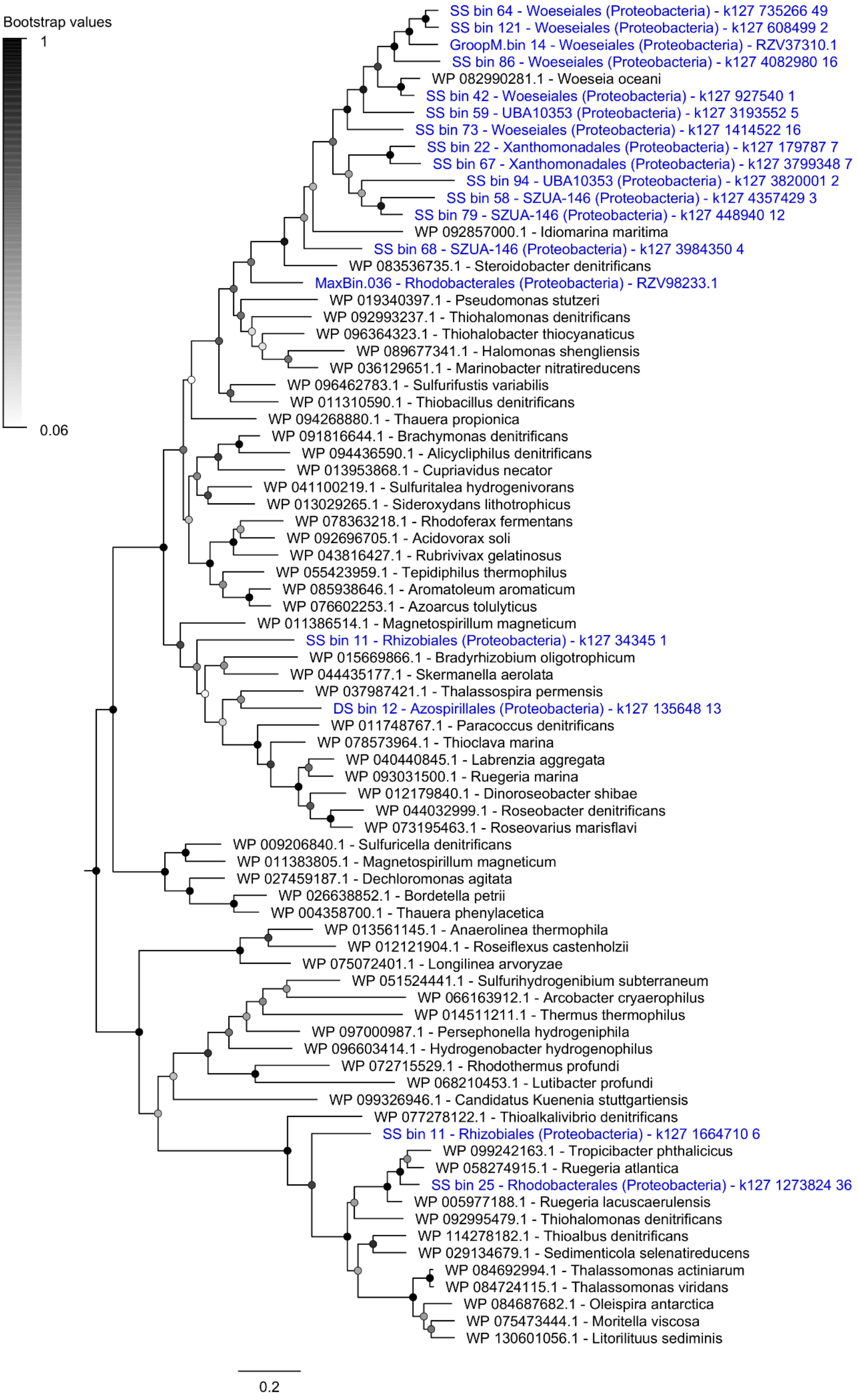
Bootstrap values
1
0.06
SS bin 64 - Woeseiales (Proteobacteria) - k127 735266 49
SS bin 121 - Woeseiales (Proteobacteria) - k127 608499 2
GroopM.bin 14 - Woeseiales (Proteobacteria) - RZV37310.1
SS bin 86 - Woeseiales (Proteobacteria) - k127 4082980 16
WP 082990281.1 - Woeseia oceani
SS bin 42 - Woeseiales (Proteobacteria) - k127 927540 1
SS bin 59 - UBA10353 (Proteobacteria) - k127 3193552 5
SS bin 73 - Woeseiales (Proteobacteria) - k127 1414522 16
SS bin 22 - Xanthomonadales (Proteobacteria) - k127 179787 7
SS bin 67 - Xanthomonadales (Proteobacteria) - k127 3799348 7
SS bin 94 - UBA10353 (Proteobacteria) - k127 3820001 2
SS bin 58 - SZUA-146 (Proteobacteria) - k127 4357429 3
SS bin 79 - SZUA-146 (Proteobacteria) - k127 448940 12
WP 092857000.1 - Idiomarina maritima
SS bin 68 - SZUA-146 (Proteobacteria) - k127 3984350 4
WP 083536735.1 - Steroidobacter denitrificans
MaxBin.036 - Rhodobacterales (Proteobacteria) - RZV98233.1
WP 019340397.1 - Pseudomonas stutzeri
WP 092993237.1 - Thiohalomonas denitrificans
WP 096364323.1 - Thiohalobacter thiocyanaticus
WP 089677341.1 - Halomonas shengliensis
WP 036129651.1 - Marinobacter nitratireducens
WP 096462783.1 - Sulfurifustis variabilis
WP 011310590.1 - Thiobacillus denitrificans
WP 094268880.1 - Thauera propionica
WP 091816644.1 - Brachymonas denitrificans
WP 094436590.1 - Alicycliphilus denitrificans
WP 013953868.1 - Cupriavidus necator
WP 041100219.1 - Sulfuritalea hydrogenivorans
WP 013029265.1 - Sideroxydans lithotrophicus
WP 078363218.1 - Rhodoferax fermentans
WP 092696705.1 - Acidovorax soli
WP 043816427.1 - Rubrivivax gelatinosus
WP 055423959.1 - Tepidiphilus thermophilus
WP 085938646.1 - Aromatoleum aromaticum
WP 076602253.1 - Azoarcus tolulyticus
WP 011386514.1 - Magnetospirillum magneticum
SS bin 11 - Rhizobiales (Proteobacteria) - k127 34345 1
WP 015669866.1 - Bradyrhizobium oligotrophicum
WP 044435177.1 - Skermanella aerolata
WP 037987421.1 - Thalassospira permensis
DS bin 12 - Azospirillales (Proteobacteria) - k127 135648 13
WP 011748767.1 - Paracoccus denitrificans
WP 078573964.1 - Thioclava marina
WP 040440845.1 - Labrenzia aggregata
WP 093031500.1 - Ruegeria marina
WP 012179840.1 - Dinoroseobacter shibae
WP 044032999.1 - Roseobacter denitrificans
WP 073195463.1 - Roseovarius marisflavi
WP 009206840.1 - Sulfuricella denitrificans
WP 011383805.1 - Magnetospirillum magneticum
WP 027459187.1 - Dechloromonas agitata
WP 026638852.1 - Bordetella petrii
WP 004358700.1 - Thauera phenylacetica
WP 013561145.1 - Anaerolinea thermophila
WP 012121904.1 - Roseiflexus castenholzii
WP 075072401.1 - Longilinea arvoryzae
WP 051524441.1 - Sulfurihydrogenibium subterraneum
WP 066163912.1 - Arcobacter cryaerophilus
WP 014511211.1 - Thermus thermophilus
WP 097000987.1 - Persephonella hydrogeniphila
WP 096603414.1 - Hydrogenobacter hydrogenophilus
WP 072715529.1 - Rhodothermus profundi
WP 068210453.1 - Lutibacter profundi
WP 099326946.1 - Candidatus Kuenenia stuttgartiensis
WP 077278122.1 - Thioalkalivibrio denitrificans
SS bin 11 - Rhizobiales (Proteobacteria) - k127 1664710 6
WP 099242163.1 - Tropicibacter phthalicicus
WP 058274915.1 - Ruegeria atlantica
SS bin 25 - Rhodobacterales (Proteobacteria) - k127 1273824 36
WP 005977188.1 - Ruegeria lacuscaerulensis
WP 092995479.1 - Thiohalomonas denitrificans
WP 114278182.1 - Thioalbus denitrificans
WP 029134679.1 - Sedimenticola selenatireducens
WP 084692994.1 - Thalassomonas actiniarum
WP 084724115.1 - Thalassomonas viridans
WP 084687682.1 - Oleispira antarctica
WP 075473444.1 - Moritella viscosa
WP 130601056.1 - Litorilituus sediminis
0.2

**Figure S16.** Maximum-likelihood tree of amino acid sequences of nitric oxide reductase B subunit (NorB), a marker for nitric oxide reduction. The tree shows sequences from permeable sediment metagenome-assembled genomes (blue) alongside representative reference sequences (black). The tree was constructed using the JTT matrix-based model, used all sites, and was bootstrapped with 50 replicates and midpoint-rooted.

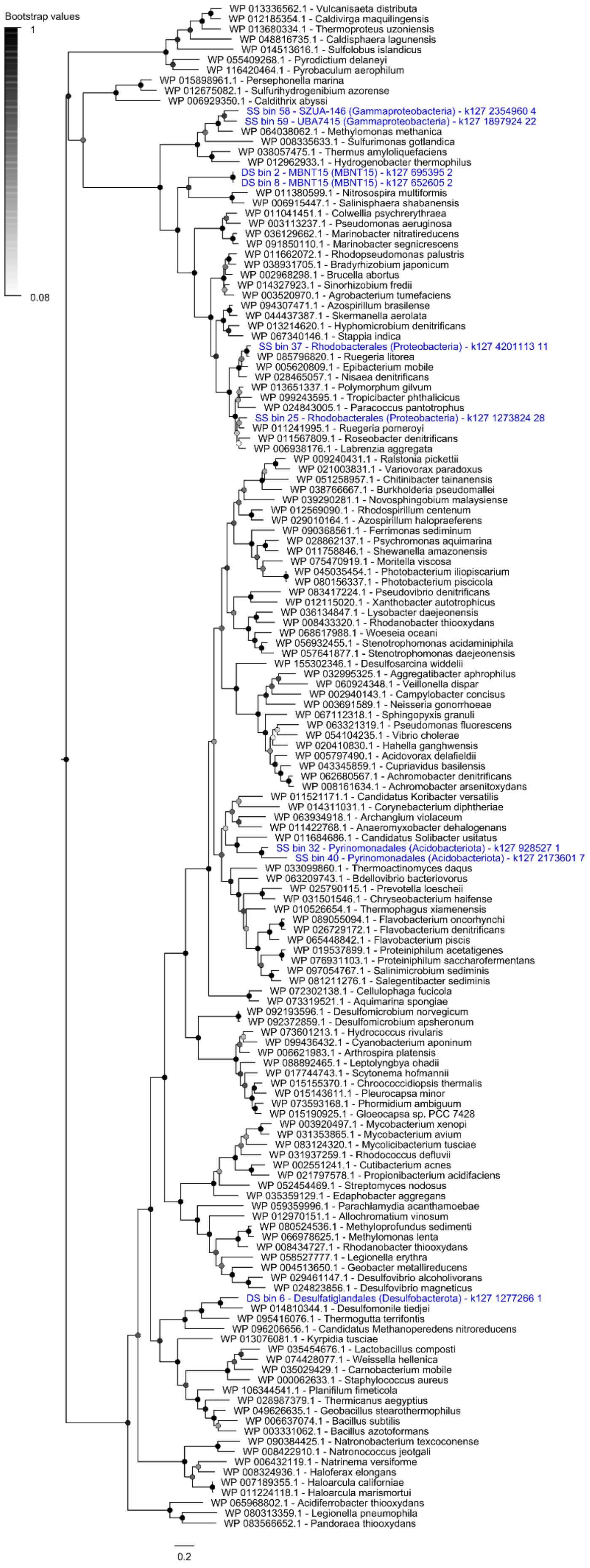
Bootstrap values
1
0.08
WP 013336562.1 - Vulcanisaeta distributa
WP 012185354.1 - Caldivirga maquilingensis
WP 013680334.1 - Thermoproteus uzoniensis
WP 048816735.1 - Caldisphaera lagunensis
WP 014513616.1 - Sulfolobus islandicus
WP 055409268.1 - Pyrodictium delaneyi
WP 116420464.1 - Pyrobaculum aerophilum
WP 015898961.1 - Persephonella marina
WP 012675082.1 - Sulfurihydrogenibium azorense
WP 006929350.1 - Caldithrix abyssi
SS bin 58 - SZUA-146 (Gammaproteobacteria) - k127 2354960 4
SS bin 59 - UBA7415 (Gammaproteobacteria) - k127 1897924 22
WP 064038062.1 - Methylomonas methanica
WP 008335633.1 - Sulfurimonas gotlandica
WP 038057475.1 - Thermus amyloliquefaciens
WP 012962933.1 - Hydrogenobacter thermophilus
DS bin 2 - MBNT15 (MBNT15) - k127 695395 2
DS bin 8 - MBNT15 (MBNT15) - k127 652605 2
WP 011380599.1 - Nitrosospira multiformis
WP 006915447.1 - Salinisphaera shabanensis
WP 011041451.1 - Colwellia psychrerythraea
WP 003113237.1 - Pseudomonas aeruginosa
WP 036129662.1 - Marinobacter nitratireducens
WP 091850110.1 - Marinobacter segnicrescens
WP 011662072.1 - Rhodopseudomonas palustris
WP 038931705.1 - Bradyrhizobium japonicum
WP 002968298.1 - Brucella abortus
WP 014327923.1 - Sinorhizobium fredii
WP 003520970.1 - Agrobacterium tumefaciens
WP 094307471.1 - Azospirillum brasilense
WP 044437387.1 - Skermanella aerolata
WP 013214620.1 - Hyphomicrobium denitrificans
WP 067340146.1 - Stappia indica
SS bin 37 - Rhodobacterales (Proteobacteria) - k127 4201113 11
WP 085796820.1 - Ruegeria litorea
WP 005620809.1 - Epibacterium mobile
WP 028465057.1 - Nisaea denitrificans
WP 013651337.1 - Polymorphum gilvum
WP 099243595.1 - Tropicibacter phthalicicus
WP 024843005.1 - Paracoccus pantotrophus
SS bin 25 - Rhodobacterales (Proteobacteria) - k127 1273824 28
WP 011241995.1 - Ruegeria pomeroyi
WP 011567809.1 - Roseobacter denitrificans
WP 006938176.1 - Labrenzia aggregata
WP 009240431.1 - Ralstonia pickettii
WP 021003831.1 - Variovorax paradoxus
WP 051258957.1 - Chitinibacter tainanensis
WP 038766667.1 - Burkholderia pseudomallei
WP 039290281.1 - Novosphingobium malaysiense
WP 012569090.1 - Rhodospirillum centenum
WP 029010164.1 - Azospirillum halopraeferens
WP 090368561.1 - Ferrimonas sediminum
WP 028862137.1 - Psychromonas aquimarina
WP 011758846.1 - Shewanella amazonensis
WP 075470919.1 - Moritella viscosa
WP 045035454.1 - Photobacterium iliopiscarium
WP 080156337.1 - Photobacterium piscicola
WP 083417224.1 - Pseudovibrio denitrificans
WP 012115020.1 - Xanthobacter autotrophicus
WP 036134847.1 - Lysobacter daejeonensis
WP 008433320.1 - Rhodanobacter thiooxydans
WP 068617988.1 - Woeseia oceani
WP 056932455.1 - Stenotrophomonas acidaminiphila
WP 057641877.1 - Stenotrophomonas daejeonensis
WP 155302346.1 - Desulfosarcina widdelii
WP 032995325.1 - Aggregatibacter aphrophilus
WP 060924348.1 - Veillonella dispar
WP 002940143.1 - Campylobacter concisus
WP 003691589.1 - Neisseria gonorrhoeae
WP 067112318.1 - Sphingopyxis granuli
WP 063321319.1 - Pseudomonas fluorescens
WP 054104235.1 - Vibrio cholerae
WP 020410830.1 - Hahella ganghwensis
WP 005797490.1 - Acidovorax delafieldii
WP 043345859.1 - Cupriavidus basilensis
WP 062680567.1 - Achromobacter denitrificans
WP 008161634.1 - Achromobacter arsenitoxydans
WP 011521171.1 - Candidatus Koribacter versatilis
WP 014311031.1 - Corynebacterium diphtheriae
WP 063934918.1 - Archangium violaceum
WP 011422768.1 - Anaeromyxobacter dehalogenans
WP 011684686.1 - Candidatus Solibacter usitatus
SS bin 32 - Pyrinomonadales (Acidobacteriota) - k127 928527 1
SS bin 40 - Pyrinomonadales (Acidobacteriota) - k127 2173601 7
WP 033099860.1 - Thermoactinomyces daqus
WP 063209743.1 - Bdellovibrio bacteriovorus
WP 025790115.1 - Prevotella loescheii
WP 031501546.1 - Chryseobacterium haifense
WP 010526654.1 - Thermophagus xiamenensis
WP 089055094.1 - Flavobacterium oncorhynchi
WP 026729172.1 - Flavobacterium denitrificans
WP 065448842.1 - Flavobacterium piscis
WP 019537899.1 - Proteiniphilum acetatigenes
WP 076931103.1 - Proteiniphilum saccharofermentans
WP 097054767.1 - Salinimicrobium sediminis
WP 081211276.1 - Salegentibacter sediminis
WP 072302138.1 - Cellulophaga fucicola
WP 073319521.1 - Aquimarina spongiae
WP 092193596.1 - Desulfomicrobium norvegicum
WP 092372859.1 - Desulfomicrobium apsheronum
WP 073601213.1 - Hydrococcus rivularis
WP 099436432.1 - Cyanobacterium aponinum
WP 006621983.1 - Arthrospira platensis
WP 088892465.1 - Leptolyngbya ohadii
WP 017744743.1 - Scytonema hofmannii
WP 015155370.1 - Chroococcidiopsis thermalis
WP 015143611.1 - Pleurocapsa minor
WP 073593168.1 - Phormidium ambiguum
WP 015190925.1 - Gloeocapsa sp. PCC 7428
WP 003920497.1 - Mycobacterium xenopi
WP 031353865.1 - Mycobacterium avium
WP 083124320.1 - Mycolicibacterium tusciae
WP 031937259.1 - Rhodococcus defluvii
WP 002551241.1 - Cutibacterium acnes
WP 021797578.1 - Propionibacterium acidifaciens
WP 052454469.1 - Streptomyces nodosus
WP 035359129.1 - Edaphobacter aggregans
WP 059359996.1 - Parachlamydia acanthamoebae
WP 012970151.1 - Allochromatium vinosum
WP 080524536.1 - Methyloprofundus sedimenti
WP 066978625.1 - Methylomonas lenta
WP 008434727.1 - Rhodanobacter thiooxydans
WP 058527777.1 - Legionella erythra
WP 004513650.1 - Geobacter metallireducens
WP 029461147.1 - Desulfovibrio alcoholivorans
WP 024823856.1 - Desulfovibrio magneticus
DS bin 6 - Desulfatiglandales (Desulfobacterota) - k127 1277266 1
WP 014810344.1 - Desulfomonile tiedjei
WP 095416076.1 - Thermogutta terrifontis
WP 096206656.1 - Candidatus Methanoperedens nitroreducens
WP 013076081.1 - Kyrpidia tusciae
WP 035454676.1 - Lactobacillus composti
WP 074428077.1 - Weissella hellenica
WP 035029429.1 - Carnobacterium mobile
WP 000062633.1 - Staphylococcus aureus
WP 106344541.1 - Planifilum fimeticola
WP 028987379.1 - Thermicanus aegyptius
WP 049626635.1 - Geobacillus stearothermophilus
WP 000637074.1 - Bacillus subtilis
WP 003331062.1 - Bacillus azotoformans
WP 090384425.1 - Natronobacterium texcoconense
WP 008422910.1 - Natronococcus jeotgali
WP 006432119.1 - Natrinema versiforme
WP 008324936.1 - Haloferax elongans
WP 007189355.1 - Haloarcula californiae
WP 011224118.1 - Haloarcula marismortui
WP 065968802.1 - Acidiferrobacter thiooxydans
WP 080313359.1 - Legionella pneumophila
WP 083566652.1 - Pandoraea thiooxydans
0.2

**Figure S17.** Maximum-likelihood tree of amino acid sequences of nitrous oxide reductase Z subunit (NosZ), a marker for nitrous oxide reduction. The tree shows sequences from permeable sediment metagenome-assembled genomes (blue) alongside representative reference sequences (black). The tree was constructed using the JTT matrix-based model, used all sites, and was bootstrapped with 50 replicates and midpoint-rooted.

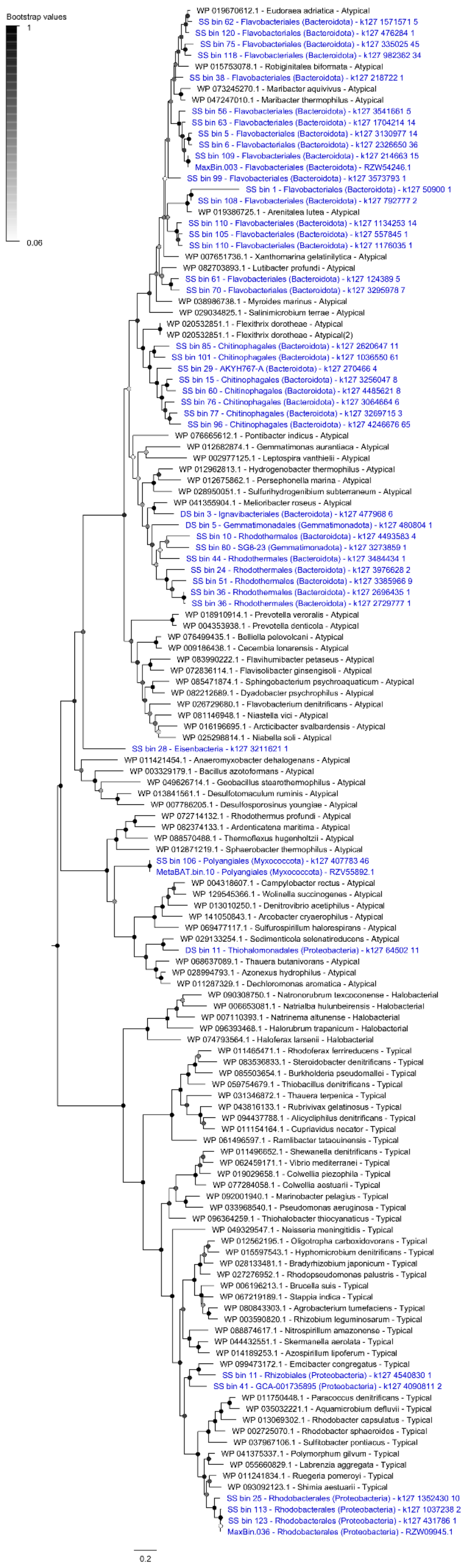
Bootstrap values
1
0.06
0.2
WP 019670612.1 - Eudoraea adriatica - Atypical
SS bin 62 - Flavobacteriales (Bacteroidota) - k127 1571571 5
SS bin 120 - Flavobacteriales (Bacteroidota) - k127 476284 1
SS bin 75 - Flavobacteriales (Bacteroidota) - k127 335025 45
SS bin 118 - Flavobacteriales (Bacteroidota) - k127 982362 34
WP 015753078.1 - Robiginitalea biformata - Atypical
SS bin 38 - Flavobacteriales (Bacteroidota) - k127 218722 1
WP 073245270.1 - Maribacter aquivivus - Atypical
WP 047247010.1 - Maribacter thermophilus - Atypical
SS bin 56 - Flavobacteriales (Bacteroidota) - k127 3541661 5
SS bin 63 - Flavobacteriales (Bacteroidota) - k127 1704214 14
SS bin 5 - Flavobacteriales (Bacteroidota) - k127 3130977 14
SS bin 6 - Flavobacteriales (Bacteroidota) - k127 2326650 36
SS bin 109 - Flavobacteriales (Bacteroidota) - k127 214663 15
MaxBin.003 - Flavobacteriales (Bacteroidota) - RZW54246.1
SS bin 99 - Flavobacteriales (Bacteroidota) - k127 3573793 1
SS bin 1 - Flavobacteriales (Bacteroidota) - k127 50900 1
SS bin 108 - Flavobacteriales (Bacteroidota) - k127 792777 2
WP 019386725.1 - Arenitalea lutea - Atypical
SS bin 110 - Flavobacteriales (Bacteroidota) - k127 1134253 14
SS bin 105 - Flavobacteriales (Bacteroidota) - k127 557845 1
SS bin 110 - Flavobacteriales (Bacteroidota) - k127 1176035 1
WP 007651736.1 - Xanthomarina gelatinilytica - Atypical
WP 082703893.1 - Lutibacter profundi - Atypical
SS bin 61 - Flavobacteriales (Bacteroidota) - k127 124389 5
SS bin 70 - Flavobacteriales (Bacteroidota) - k127 3295978 7
WP 038986738.1 - Myroides marinus - Atypical
WP 029034825.1 - Salinimicrobium terrae - Atypical
WP 020532851.1 - Flexithrix dorotheae - Atypical
WP 020532851.1 - Flexithrix dorotheae - Atypical(2)
SS bin 85 - Chitinophagales (Bacteroidota) - k127 2620647 11
SS bin 101 - Chitinophagales (Bacteroidota) - k127 1036550 61
SS bin 29 - AKYH767-A (Bacteroidota) - k127 270466 4
SS bin 15 - Chitinophagales (Bacteroidota) - k127 3256047 8
SS bin 60 - Chitinophagales (Bacteroidota) - k127 4485621 8
SS bin 76 - Chitinophagales (Bacteroidota) - k127 3064664 6
SS bin 77 - Chitinophagales (Bacteroidota) - k127 3269715 3
SS bin 96 - Chitinophagales (Bacteroidota) - k127 4246676 65
WP 076665612.1 - Pontibacter indicus - Atypical
WP 012682874.1 - Gemmatimonas aurantiaca - Atypical
WP 002977125.1 - Leptospira vanthielii - Atypical
WP 012962813.1 - Hydrogenobacter thermophilus - Atypical
WP 012675862.1 - Persephonella marina - Atypical
WP 028950051.1 - Sulfurihydrogenibium subterraneum - Atypical
WP 041355904.1 - Melioribacter roseus - Atypical
DS bin 3 - Ignavibacteriales (Bacteroidota) - k127 477968 6
DS bin 5 - Gemmatimonadales (Gemmatimonadota) - k127 480804 1
SS bin 10 - Rhodothermales (Bacteroidota) - k127 4493583 4
SS bin 80 - SG8-23 (Gemmatimonadota) - k127 3273859 1
SS bin 44 - Rhodothermales (Bacteroidota) - k127 3484434 1
SS bin 24 - Rhodothermales (Bacteroidota) - k127 3976628 2
SS bin 51 - Rhodothermales (Bacteroidota) - k127 3385966 9
SS bin 36 - Rhodothermales (Bacteroidota) - k127 2696435 1
SS bin 36 - Rhodothermales (Bacteroidota) - k127 2729777 1
WP 018910914.1 - Prevotella veroralis - Atypical
WP 004353938.1 - Prevotella denticola - Atypical
WP 076499435.1 - Belliella pelovolcani - Atypical
WP 009186438.1 - Cecembia lonarensis - Atypical
WP 083990222.1 - Flavihumibacter petaseus - Atypical
WP 072836114.1 - Flavisolibacter ginsengisoli - Atypical
WP 085471874.1 - Sphingobacterium psychroaquaticum - Atypical
WP 082212689.1 - Dyadobacter psychrophilus - Atypical
WP 026729680.1 - Flavobacterium denitrificans - Atypical
WP 081146948.1 - Niastella vici - Atypical
WP 016196695.1 - Arcticibacter svalbardensis - Atypical
WP 025298814.1 - Niabella soli - Atypical
SS bin 28 - Eisenbacteria - k127 3211621 1
WP 011421454.1 - Anaeromyxobacter dehalogenans - Atypical
WP 003329179.1 - Bacillus azotoformans - Atypical
WP 049626714.1 - Geobacillus stearothermophilus - Atypical
WP 013841561.1 - Desulfotomaculum ruminis - Atypical
WP 007786205.1 - Desulfosporosinus youngiae - Atypical
WP 072714132.1 - Rhodothermus profundi - Atypical
WP 082374133.1 - Ardenticatena maritima - Atypical
WP 088570488.1 - Thermoflexus hugenholtzii - Atypical
WP 012871219.1 - Sphaerobacter thermophilus - Atypical
SS bin 106 - Polyangiales (Myxococcota) - k127 407783 46
MetaBAT.bin.10 - Polyangiales (Myxococcota) - RZV55892.1
WP 004318607.1 - Campylobacter rectus - Atypical
WP 129545366.1 - Wolinella succinogenes - Atypical
WP 013010250.1 - Denitrovibrio acetiphilus - Atypical
WP 141050843.1 - Arcobacter cryaerophilus - Atypical
WP 069477117.1 - Sulfurospirillum halorespirans - Atypical
WP 029133254.1 - Sedimenticola selenatireducens - Atypical
DS bin 11 - Thiohalomonadales (Proteobacteria) - k127 64502 11
WP 068637089.1 - Thauera butanivorans - Atypical
WP 028994793.1 - Azonexus hydrophilus - Atypical
WP 011287329.1 - Dechloromonas aromatica - Atypical
WP 090308750.1 - Natronorubrum texcoconense - Halobacterial
WP 006653081.1 - Natrialba hulunbeirensis - Halobacterial
WP 007110393.1 - Natrinema altunense - Halobacterial
WP 096393468.1 - Halorubrum trapanicum - Halobacterial
WP 074793564.1 - Haloferax larsenii - Halobacterial
WP 011465471.1 - Rhodoferax ferrireducens - Typical
WP 083536833.1 - Steroidobacter denitrificans - Typical
WP 085503654.1 - Burkholderia pseudomallei - Typical
WP 059754679.1 - Thiobacillus denitrificans - Typical
WP 031346872.1 - Thauera terpenica - Typical
WP 043816133.1 - Rubrivivax gelatinosus - Typical
WP 094437788.1 - Alicycliphilus denitrificans - Typical
WP 011154164.1 - Cupriavidus necator - Typical
WP 061496597.1 - Ramlibacter tataouinensis - Typical
WP 011496652.1 - Shewanella denitrificans - Typical
WP 062459171.1 - Vibrio mediterranei - Typical
WP 019029658.1 - Colwellia piezophila - Typical
WP 077284058.1 - Colwellia aestuarii - Typical
WP 092001940.1 - Marinobacter pelagius - Typical
WP 033968540.1 - Pseudomonas aeruginosa - Typical
WP 096364259.1 - Thiohalobacter thiocyanaticus - Typical
WP 049329547.1 - Neisseria meningitidis - Typical
WP 012562195.1 - Oligotropha carboxidovorans - Typical
WP 015597543.1 - Hyphomicrobium denitrificans - Typical
WP 028133481.1 - Bradyrhizobium japonicum - Typical
WP 027276952.1 - Rhodopseudomonas palustris - Typical
WP 006196213.1 - Brucella suis - Typical
WP 067219189.1 - Stappia indica - Typical
WP 080843303.1 - Agrobacterium tumefaciens - Typical
WP 003590820.1 - Rhizobium leguminosarum - Typical
WP 088874617.1 - Nitrospirillum amazonense - Typical
WP 044432551.1 - Skermanella aerolata - Typical
WP 014189253.1 - Azospirillum lipoferum - Typical
WP 099473172.1 - Emcibacter congregatus - Typical
SS bin 11 - Rhizobiales (Proteobacteria) - k127 4540830 1
SS bin 41 - GCA-001735895 (Proteobacteria) - k127 4090811 2
WP 011750448.1 - Paracoccus denitrificans - Typical
WP 035032221.1 - Aquamicrobium defluvii - Typical
WP 013069302.1 - Rhodobacter capsulatus - Typical
WP 002725070.1 - Rhodobacter sphaeroides - Typical
WP 037967106.1 - Sulfitobacter pontiacus - Typical
WP 041375337.1 - Polymorphum gilvum - Typical
WP 055660829.1 - Labrenzia aggregata - Typical
WP 011241834.1 - Ruegeria pomeroyi - Typical
WP 093092123.1 - Shimia aestuarii - Typical
SS bin 25 - Rhodobacterales (Proteobacteria) - k127 1352430 10
SS bin 113 - Rhodobacterales (Proteobacteria) - k127 1037238 2
SS bin 123 - Rhodobacterales (Proteobacteria) - k127 431786 1
MaxBin.036 - Rhodobacterales (Proteobacteria) - RZW09945.1

**Figure S18.** Maximum-likelihood tree of amino acid sequences of ammonifying nitrite reductase A subunit (NrfA), a marker for dissimilatory nitrate reduction to ammonium. The tree shows sequences from permeable sediment metagenome-assembled genomes (blue) alongside representative reference sequences (black). The tree was constructed using the JTT matrix-based model, used all sites, and was bootstrapped with 50 replicates and midpoint-rooted.

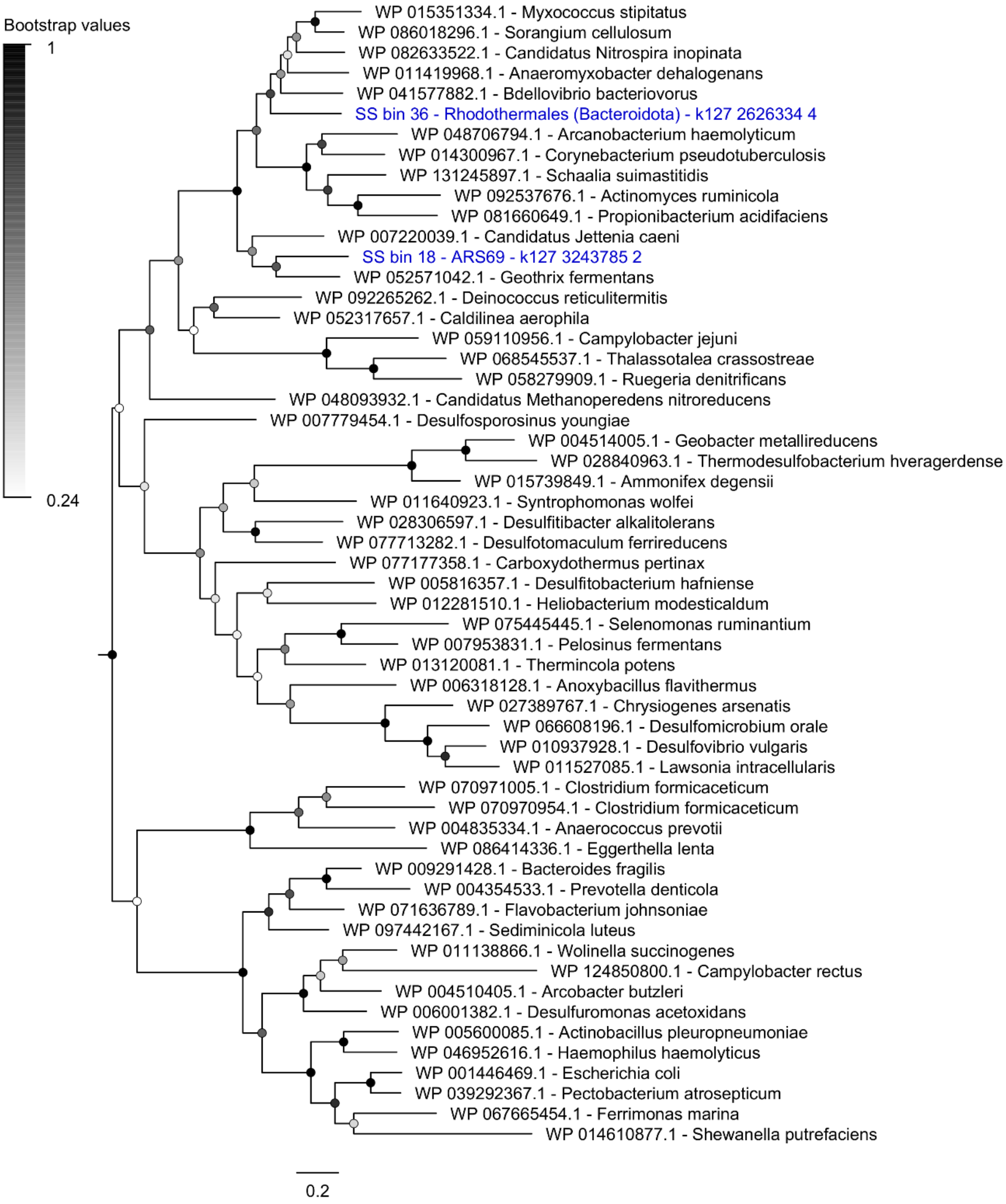

**Figure S19.** Maximum-likelihood tree of amino acid sequences of group 3 [NiFe]-hydrogenase large subunits, a marker for hydrogen production during fermentation processes. The tree shows sequences from permeable sediment metagenome-assembled genomes (blue) alongside representative reference sequences (black). The subgroup of each reference sequence is denoted according to the HydDB classification scheme. The tree was constructed using the JTT matrix-based model, used all sites, and was bootstrapped with 50 replicates and midpoint-rooted.

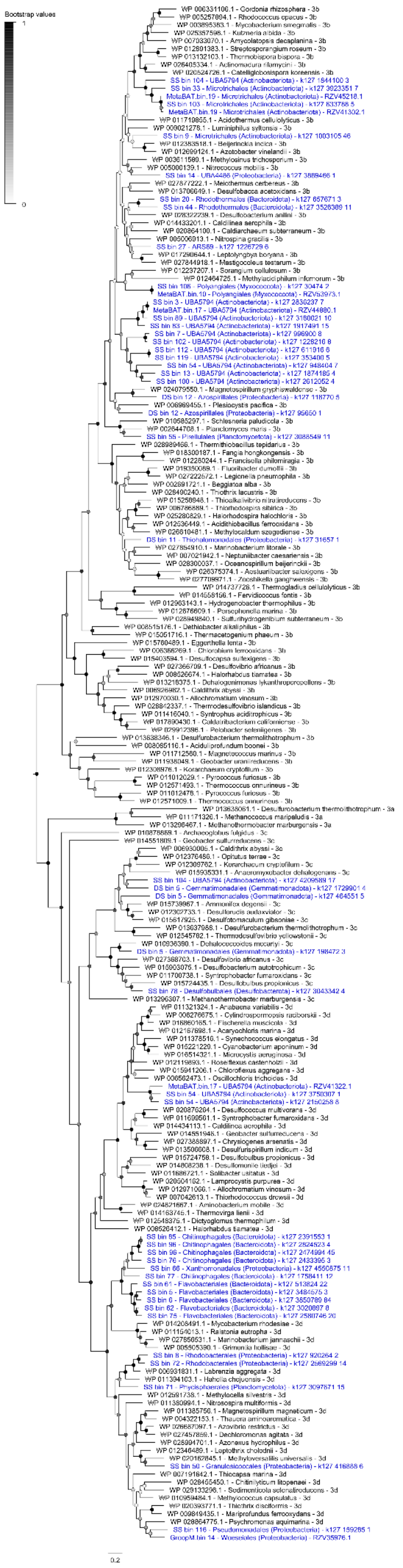
Bootstrap values
1
0
WP 006331106.1 - Gordonia rhizosphera - 3b
WP 005257894.1 - Rhodococcus opacus - 3b
WP 003895383.1 - Mycobacterium smegmatis - 3b
WP 025357598.1 - Kutzneria albida - 3b
WP 007033070.1 - Amycolatopsis decaplanina - 3b
WP 012891383.1 - Streptosporangium roseum - 3b
WP 013132103.1 - Thermobispora bispora - 3b
WP 026405334.1 - Actinomadura rifamycini - 3b
WP 020524726.1 - Catelliglobosispora koreensis - 3b
SS bin 104 - UBA5794 (Actinobacteriota) - k127 1844100 3
SS bin 33 - Microtrichales (Actinobacteriota) - k127 3923351 7
MetaBAT.bin.19 - Microtrichales (Actinobacteriota) - RZV45218.1
SS bin 103 - Microtrichales (Actinobacteriota) - k127 633788 5
MetaBAT.bin.19 - Microtrichales (Actinobacteriota) - RZV41302.1
WP 011719855.1 - Acidothermus cellulolyticus - 3b
WP 009021278.1 - Luminiphilus syltensis - 3b
SS bin 9 - Microtrichales (Actinobacteriota) - k127 1003105 46
WP 012383518.1 - Beijerinckia indica - 3b
WP 012699124.1 - Azotobacter vinelandii - 3b
WP 003611580.1 - Methylosinus trichosporium - 3b
WP 005000139.1 - Nitrococcus mobilis - 3b
SS bin 14 - UBA4486 (Proteobacteria) - k127 3889466 1
WP 027877222.1 - Meiothermus cerbereus - 3b
WP 013706649.1 - Desulfobacca acetoxidans - 3b
SS bin 20 - Rhodothermales (Bacteroidota) - k127 657671 3
SS bin 44 - Rhodothermales (Bacteroidota) - k127 3528309 11
WP 028322239.1 - Desulfobacterium anilini - 3b
WP 014433201.1 - Caldilinea aerophila - 3b
WP 020864100.1 - Caldiarchaeum subterraneum - 3b
WP 005006913.1 - Nitrospina gracilis - 3b
SS bin 27 - ARS69 - k127 1226729 6
WP 017290644.1 - Leptolyngbya boryana - 3b
WP 027844918.1 - Mastigocoleus testarum - 3b
WP 012237207.1 - Sorangium cellulosum - 3b
WP 012464725.1 - Methylacidiphilum infernorum - 3b
SS bin 106 - Polyangiales (Myxococcota) - k127 30474 2
MetaBAT.bin.10 - Polyangiales (Myxococcota) - RZV53973.1
SS bin 3 - UBA5794 (Actinobacteriota) - k127 2830237 7
MetaBAT.bin.17 - UBA5794 (Actinobacteriota) - RZV44880.1
SS bin 89 - UBA5794 (Actinobacteriota) - k127 3180021 10
SS bin 83 - UBA5794 (Actinobacteriota) - k127 1917491 15
SS bin 7 - UBA5794 (Actinobacteriota) - k127 996900 8
SS bin 102 - UBA5794 (Actinobacteriota) - k127 1228216 8
SS bin 112 - UBA5794 (Actinobacteriota) - k127 611916 6
SS bin 119 - UBA5794 (Actinobacteriota) - k127 353400 5
SS bin 54 - UBA5794 (Actinobacteriota) - k127 948404 7
SS bin 13 - UBA5794 (Actinobacteriota) - k127 1874185 4
SS bin 100 - UBA5794 (Actinobacteriota) - k127 2612052 4
WP 024079550.1 - Magnetospirillum gryphiswaldense - 3b
DS bin 12 - Azospirillales (Proteobacteria) - k127 118770 5
WP 006969455.1 - Plesiocystis pacifica - 3b
DS bin 12 - Azospirillales (Proteobacteria) - k127 95650 1
WP 010585297.1 - Schlesneria paludicola - 3b
WP 002644708.1 - Planctomyces maris - 3b
SS bin 55 - Pirellulales (Planctomycetota) - k127 3088549 11
WP 028989466.1 - Thermithiobacillus tepidarius - 3b
WP 018300187.1 - Fangia hongkongensis - 3b
WP 012280244.1 - Francisella philomiragia - 3b
WP 019350089.1 - Fluoribacter dumoffii - 3b
WP 027222572.1 - Legionella pneumophila - 3b
WP 002691721.1 - Beggiatoa alba - 3b
WP 028490240.1 - Thiothrix lacustris - 3b
WP 015258948.1 - Thioalkalivibrio nitratireducens - 3b
WP 006786889.1 - Thiorhodospira sibirica - 3b
WP 025280829.1 - Halorhodospira halochloris - 3b
WP 012536449.1 - Acidithiobacillus ferrooxidans - 3b
WP 026610481.1 - Methylocaldum szegediense - 3b
DS bin 11 - Thiohalomonadales (Proteobacteria) - k127 31657 1
WP 027854910.1 - Marinobacterium litorale - 3b
WP 007021942.1 - Neptuniibacter caesariensis - 3b
WP 028300037.1 - Oceanospirillum beijerinckii - 3b
WP 026375374.1 - Aestuariibacter salexigens - 3b
WP 027709971.1 - Zooshikella ganghwensis - 3b
WP 014737726.1 - Thermogladius cellulolyticus - 3b
WP 014558156.1 - Fervidicoccus fontis - 3b
WP 012963143.1 - Hydrogenobacter thermophilus - 3b
WP 012676609.1 - Persephonella marina - 3b
WP 028949840.1 - Sulfurihydrogenibium subterraneum - 3b
WP 008515176.1 - Dethiobacter alkaliphilus - 3b
WP 015051716.1 - Thermacetogenium phaeum - 3b
WP 015760489.1 - Eggerthella lenta - 3b
WP 006366269.1 - Chlorobium ferrooxidans - 3b
WP 015403594.1 - Desulfocapsa sulfexigens - 3b
WP 027366709.1 - Desulfovibrio africanus - 3b
WP 008526674.1 - Halorhabdus tiamatea - 3b
WP 013218375.1 - Dehalogenimonas lykanthroporepellens - 3b
WP 006926982.1 - Caldithrix abyssi - 3b
WP 012970030.1 - Allochromatium vinosum - 3b
WP 028842337.1 - Thermodesulfovibrio islandicus - 3b
WP 011416040.1 - Syntrophus aciditrophicus - 3b
WP 017890430.1 - Caldatribacterium californiense - 3b
WP 029912396.1 - Pelobacter seleniigenes - 3b
WP 013638346.1 - Desulfurobacterium thermolithotrophum - 3b
WP 008085116.1 - Aciduliprofundum boonei - 3b
WP 011712560.1 - Magnetococcus marinus - 3b
WP 011938049.1 - Geobacter uraniireducens - 3b
WP 012308976.1 - Korarchaeum cryptofilum - 3b
WP 011012029.1 - Pyrococcus furiosus - 3b
WP 012571493.1 - Thermococcus onnurineus - 3b
WP 011012476.1 - Pyrococcus furiosus - 3b
WP 012571009.1 - Thermococcus onnurineus - 3b
WP 013638061.1 - Desulfurobacterium thermolithotrophum - 3a
WP 011171326.1 - Methanococcus maripaludis - 3a
WP 013296467.1 - Methanothermobacter marburgensis - 3a
WP 010878869.1 - Archaeoglobus fulgidus - 3c
WP 014551809.1 - Geobacter sulfurreducens - 3c
WP 006930006.1 - Caldithrix abyssi - 3c
WP 012376486.1 - Opitutus terrae - 3c
WP 012309762.1 - Korarchaeum cryptofilum - 3c
WP 015935331.1 - Anaeromyxobacter dehalogenans - 3c
SS bin 104 - UBA5794 (Actinobacteriota) - k127 4209589 17
DS bin 5 - Gemmatimonadales (Gemmatimonadota) - k127 1729901 4
DS bin 5 - Gemmatimonadales (Gemmatimonadota) - k127 464551 5
WP 015739967.1 - Ammonifex degensii - 3c
WP 012302733.1 - Desulforudis audaxviator - 3c
WP 015617925.1 - Desulfotomaculum gibsoniae - 3c
WP 013637988.1 - Desulfurobacterium thermolithotrophum - 3c
WP 012545782.1 - Thermodesulfovibrio yellowstonii - 3c
WP 010936390.1 - Dehalococcoides mccartyi - 3c
DS bin 5 - Gemmatimonadales (Gemmatimonadota) - k127 198472 3
WP 027368703.1 - Desulfovibrio africanus - 3c
WP 015903075.1 - Desulfobacterium autotrophicum - 3c
WP 011700738.1 - Syntrophobacter fumaroxidans - 3c
WP 015724435.1 - Desulfobulbus propionicus - 3c
SS bin 78 - Desulfobulbales (Desulfobacterota) - k127 3043342 4
WP 013296307.1 - Methanothermobacter marburgensis - 3c
WP 011321324.1 - Anabaena variabilis - 3d
WP 006276675.1 - Cylindrospermopsis raciborskii - 3d
WP 016860165.1 - Fischerella muscicola - 3d
WP 012167698.1 - Acaryochloris marina - 3d
WP 011378516.1 - Synechococcus elongatus - 3d
WP 015221229.1 - Cyanobacterium aponinum - 3d
WP 016514321.1 - Microcystis aeruginosa - 3d
WP 012119693.1 - Roseiflexus castenholzii - 3d
WP 015941206.1 - Chloroflexus aggregans - 3d
WP 006562473.1 - Oscillochloris trichoides - 3d
MetaBAT.bin.17 - UBA5794 (Actinobacteriota) - RZV41322.1
SS bin 54 - UBA5794 (Actinobacteriota) - k127 3750307 1
SS bin 54 - UBA5794 (Actinobacteriota) - k127 2150258 8
WP 020876264.1 - Desulfococcus multivorans - 3d
WP 011699561.1 - Syntrophobacter fumaroxidans - 3d
WP 014434113.1 - Caldilinea aerophila - 3d
WP 014551946.1 - Geobacter sulfurreducens - 3d
WP 027388897.1 - Chrysiogenes arsenatis - 3d
WP 013506608.1 - Desulfurispirillum indicum - 3d
WP 015724758.1 - Desulfobulbus propionicus - 3d
WP 014808238.1 - Desulfomonile tiedjei - 3d
WP 011686721.1 - Solibacter usitatus - 3d
WP 020504162.1 - Lamprocystis purpurea - 3d
WP 012971066.1 - Allochromatium vinosum - 3d
WP 007042613.1 - Thiorhodococcus drewsii - 3d
WP 024821667.1 - Aminobacterium mobile - 3d
WP 014163745.1 - Thermovirga lienii - 3d
WP 012548375.1 - Dictyoglomus thermophilum - 3d
WP 008526412.1 - Halorhabdus tiamatea - 3d
SS bin 65 - Chitinophagales (Bacteroidota) - k127 2391553 1
SS bin 96 - Chitinophagales (Bacteroidota) - k127 2824523 4
SS bin 96 - Chitinophagales (Bacteroidota) - k127 2474994 45
SS bin 76 - Chitinophagales (Bacteroidota) - k127 2433395 3
SS bin 66 - Xanthomonadales (Proteobacteria) - k127 4550875 11
SS bin 77 - Chitinophagales (Bacteroidota) - k127 1758411 12
SS bin 61 - Flavobacteriales (Bacteroidota) - k127 513824 22
SS bin 5 - Flavobacteriales (Bacteroidota) - k127 3484575 3
SS bin 6 - Flavobacteriales (Bacteroidota) - k127 3850789 84
SS bin 62 - Flavobacteriales (Bacteroidota) - k127 3020897 8
SS bin 75 - Flavobacteriales (Bacteroidota) - k127 2580746 20
WP 014208491.1 - Mycobacterium rhodesiae - 3d
WP 011154013.1 - Ralstonia eutropha - 3d
WP 027856531.1 - Marinobacterium jannaschii - 3d
WP 005505390.1 - Grimontia hollisae - 3d
SS bin 8 - Rhodobacterales (Proteobacteria) - k127 920264 2
SS bin 72 - Rhodobacterales (Proteobacteria) - k127 2569299 14
WP 006931831.1 - Labrenzia aggregata - 3d
WP 011394103.1 - Hahella chejuensis - 3d
SS bin 71 - Phycisphaerales (Planctomycetota) - k127 3097671 15
WP 012591738.1 - Methylocella silvestris - 3d
WP 011380994.1 - Nitrosospira multiformis - 3d
WP 011385756.1 - Magnetospirillum magneticum - 3d
WP 004322153.1 - Thauera aminoaromatica - 3d
WP 026687097.1 - Azovibrio restrictus - 3d
WP 027457859.1 - Dechloromonas agitata - 3d
WP 028994701.1 - Azonexus hydrophilus - 3d
WP 012346489.1 - Leptothrix cholodnii - 3d
WP 020162845.1 - Methyloversatilis universalis - 3d
SS bin 50 - Granulosicoccales (Proteobacteria) - k127 416888 6
WP 007191642.1 - Thiocapsa marina - 3d
WP 028455450.1 - Chitinilyticum litopenaei - 3d
WP 029133296.1 - Sedimenticola selenatireducens - 3d
WP 010959484.1 - Methylococcus capsulatus - 3d
WP 020393771.1 - Thiothrix disciformis - 3d
WP 009849435.1 - Mariprofundus ferrooxydans - 3d
WP 028864775.1 - Psychromonas aquimarina - 3d
SS bin 116 - Pseudomonadales (Proteobacteria) - k127 159265 1
GroopM.bin 14 - Woeseiales (Proteobacteria) - RZV35976.1
0.2

**Figure S20.** Maximum-likelihood tree of amino acid sequences of metal-reducing decaheme-associated outer membrane protein (MtrB), a marker for iron(III) reduction. The tree shows sequences from permeable sediment metagenome-assembled genomes (blue) alongside representative reference sequences (black). The tree was constructed using the JTT matrix-based model, used all sites, and was bootstrapped with 50 replicates and midpoint-rooted. Note no binned reads for OmcB, an analogous outer membrane protein in *Geobacter* and *Desulfuromonas* species, were detected.

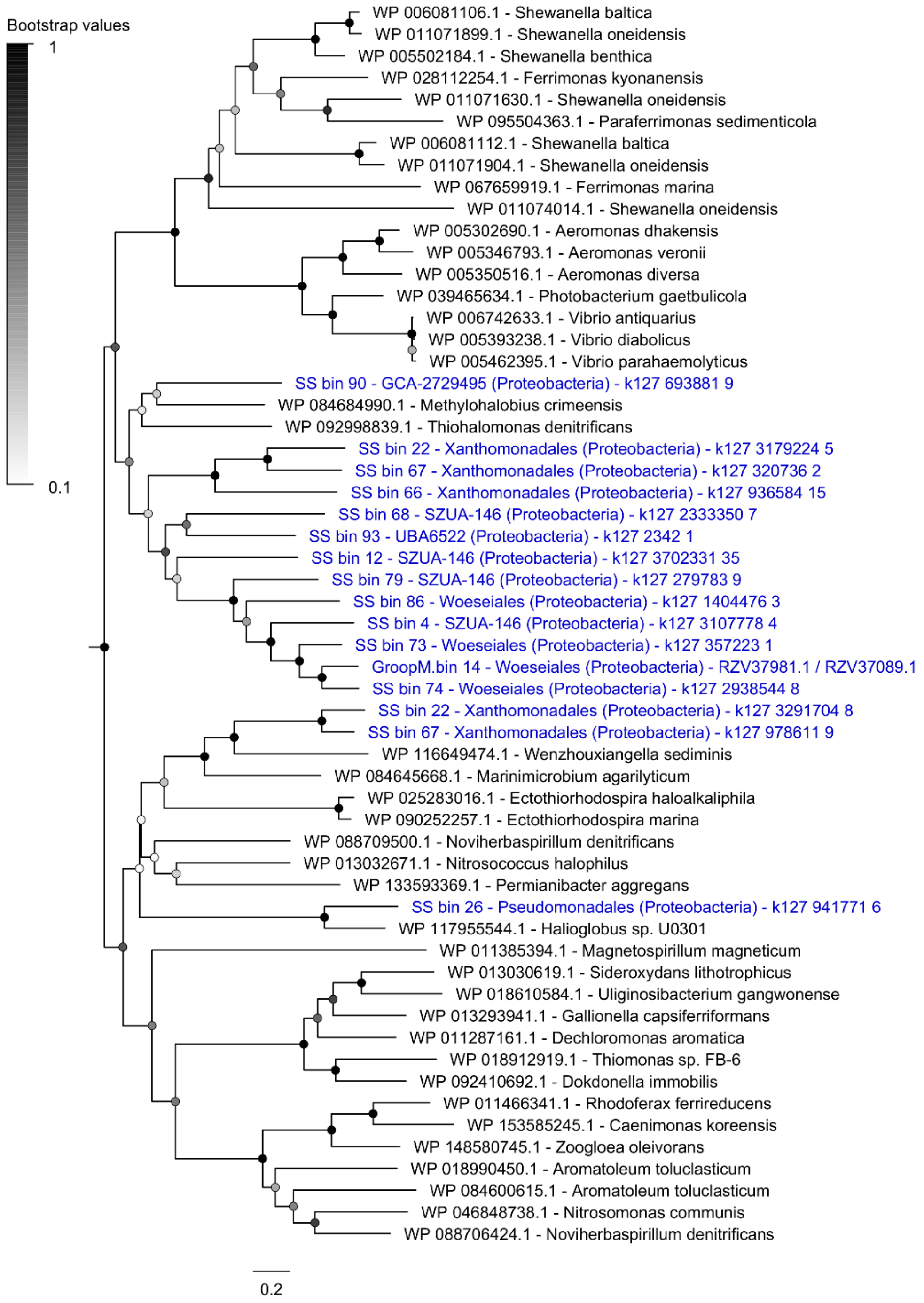

**Figure S21.** Maximum-likelihood tree of amino acid sequences of reductive dehalogenase A subunit (RdhA), a marker for the dehalogenation of diverse organohalide compounds. The tree shows sequences from permeable sediment metagenome-assembled genomes (blue) alongside representative reference sequences (black). The tree was constructed using the JTT matrix-based model, used all sites, and was bootstrapped with 50 replicates and midpoint-rooted.

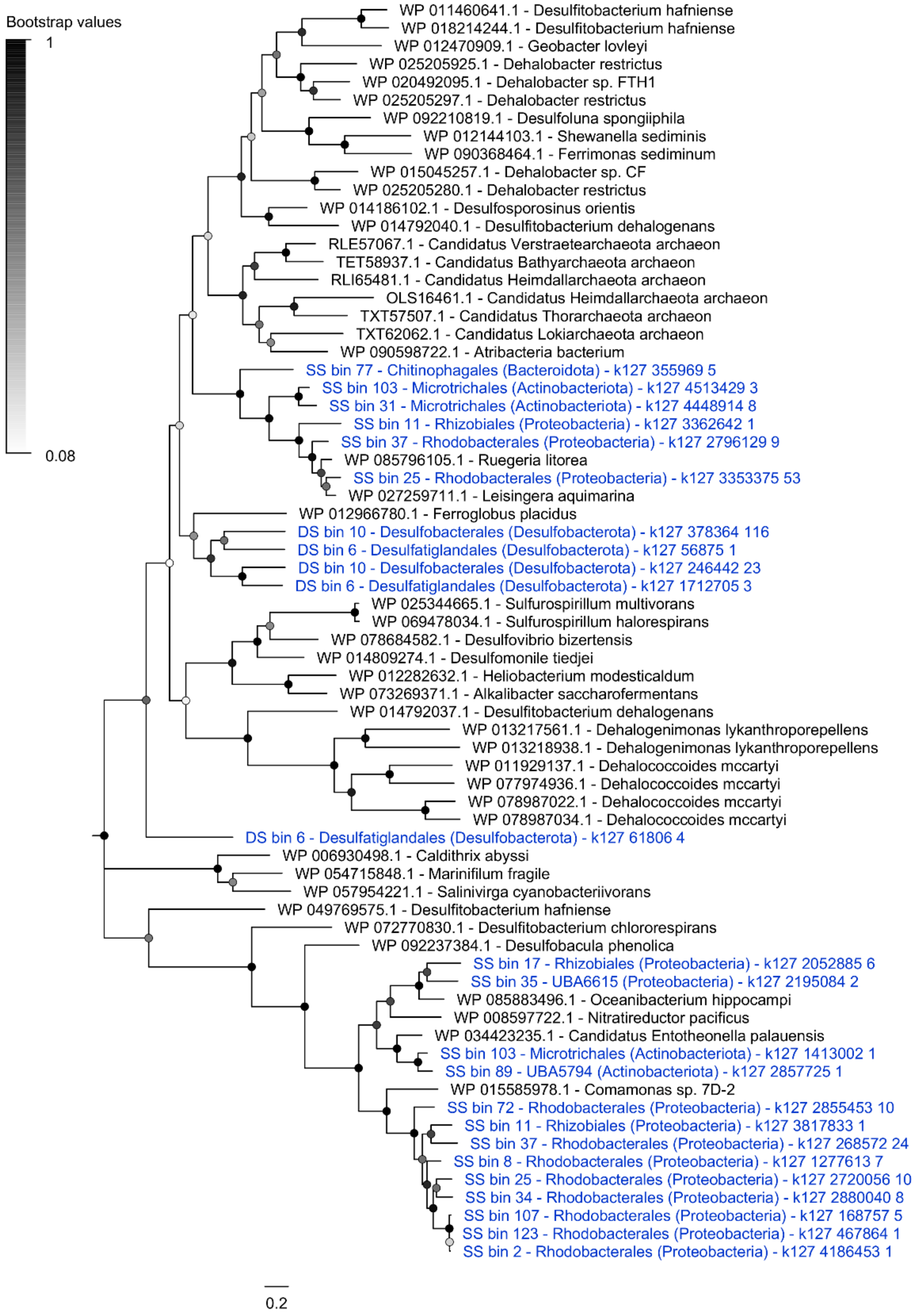

**Figure S22.** Maximum-likelihood tree of amino acid sequences of fumarate reductase A subunit (FrdA), a marker for fumarate reduction. The tree shows sequences from permeable sediment metagenome-assembled genomes (blue) alongside representative reference sequences (black). The tree was constructed using the JTT matrix-based model, used all sites, and was bootstrapped with 50 replicates and midpoint-rooted. Note only canonical fumarate reductases are shown and it is possible that homologous enzymes within the complex II superfamily also mediate fumarate reduction either reversibly or unidirectionally.

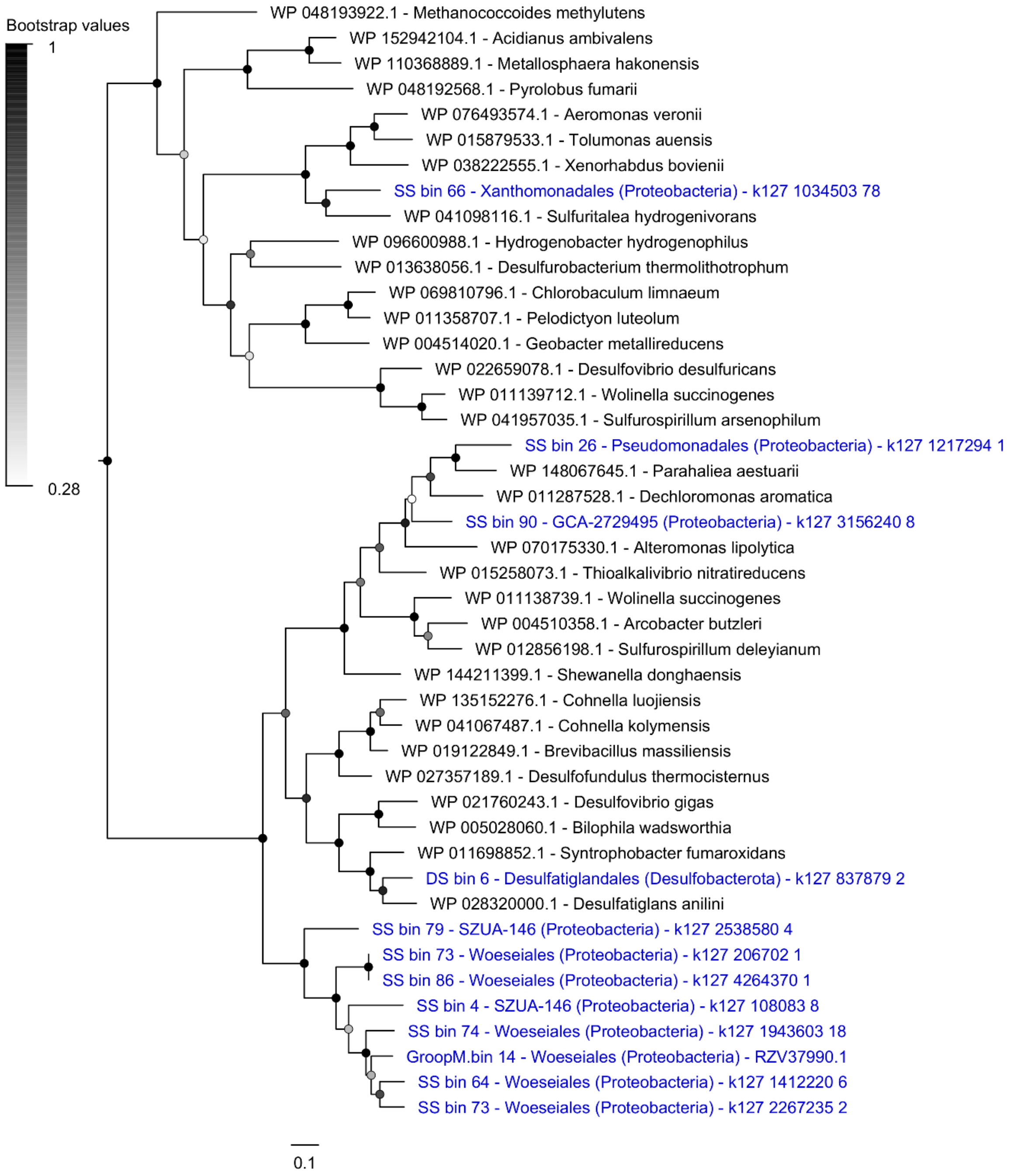

**Figure S23.** Maximum-likelihood tree of amino acid sequences of microbial rhodopsins. Only clades known to support energy transduction through coupling photon capture to proton or sodium translocation are shown. The tree shows sequences from permeable sediment metagenome-assembled genomes (blue) alongside representative reference sequences (black). The tree was constructed using the JTT matrix-based model, used all sites, and was bootstrapped with 50 replicates and midpoint-rooted.

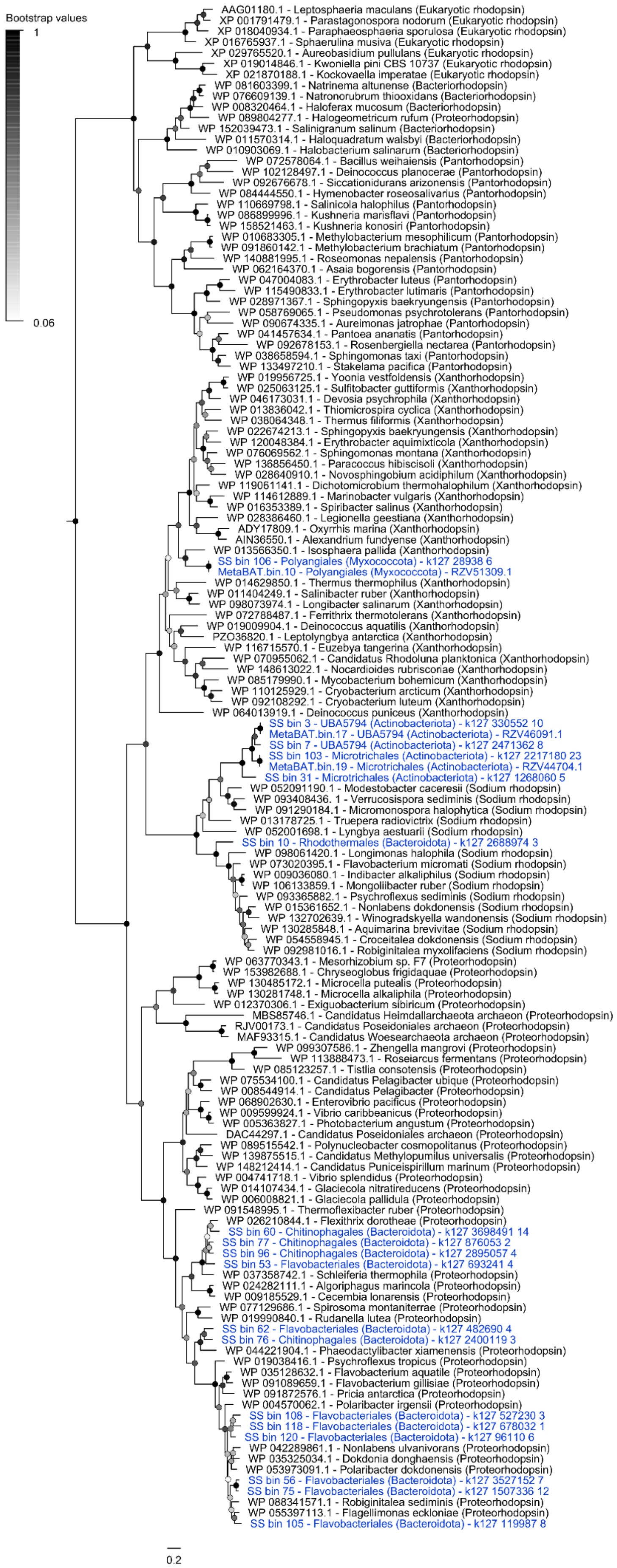

Bootstrap values
1
0.06
AAG01180.1 - Leptosphaeria maculans (Eukaryotic rhodopsin)
XP 001791479.1 - Parastagonospora nodorum (Eukaryotic rhodopsin)
XP 018040934.1 - Paraphaeosphaeria sporulosa (Eukaryotic rhodopsin)
XP 016765937.1 - Sphaerulina musiva (Eukaryotic rhodopsin)
XP 029765520.1 - Aureobasidium pullulans (Eukaryotic rhodopsin)
XP 019014846.1 - Kwoniella pini CBS 10737 (Eukaryotic rhodopsin)
XP 021870188.1 - Kockovaella imperatae (Eukaryotic rhodopsin)
WP 081603399.1 - Natrinema altunense (Bacteriorhodopsin)
WP 076609139.1 - Natronorubrum thiooxidans (Bacteriorhodopsin)
WP 008320464.1 - Haloferax mucosum (Bacteriorhodopsin)
WP 089804277.1 - Halogeometricum rufum (Proteorhodopsin)
WP 152039473.1 - Salinigranum salinum (Bacteriorhodopsin)
WP 011570314.1 - Haloquadratum walsbyi (Bacteriorhodopsin)
WP 010903069.1 - Halobacterium salinarum (Bacteriorhodopsin)
WP 072578064.1 - Bacillus weihaiensis (Pantorhodopsin)
WP 102128497.1 - Deinococcus planocerae (Pantorhodopsin)
WP 092676678.1 - Siccationidurans arizonensis (Pantorhodopsin)
WP 084444550.1 - Hymenobacter roseosalivarius (Pantorhodopsin)
WP 110669798.1 - Salinicola halophilus (Pantorhodopsin)
WP 086899996.1 - Kushneria marisflavi (Pantorhodopsin)
WP 158521463.1 - Kushneria konosiri (Pantorhodopsin)
WP 010683305.1 - Methylobacterium mesophilicum (Pantorhodopsin)
WP 091860142.1 - Methylobacterium brachiatum (Pantorhodopsin)
WP 140881995.1 - Roseomonas nepalensis (Pantorhodopsin)
WP 062164370.1 - Asaia bogorensis (Pantorhodopsin)
WP 047004083.1 - Erythrobacter luteus (Pantorhodopsin)
WP 115490833.1 - Erythrobacter lutimaris (Pantorhodopsin)
WP 028971367.1 - Sphingopyxis baekryungensis (Pantorhodopsin)
WP 058769065.1 - Pseudomonas psychrotolerans (Pantorhodopsin)
WP 090674335.1 - Aureimonas jatrophae (Pantorhodopsin)
WP 041457634.1 - Pantoea ananatis (Pantorhodopsin)
WP 092678153.1 - Rosenbergiella nectarea (Pantorhodopsin)
WP 038658594.1 - Sphingomonas taxi (Pantorhodopsin)
WP 133497210.1 - Stakelama pacifica (Pantorhodopsin)
WP 019956725.1 - Yoonia vestfoldensis (Xanthorhodopsin)
WP 025063125.1 - Sulfitobacter guttiformis (Xanthorhodopsin)
WP 046173031.1 - Devosia psychrophila (Xanthorhodopsin)
WP 013836042.1 - Thiomicrospira cyclica (Xanthorhodopsin)
WP 038064348.1 - Thermus filiformis (Xanthorhodopsin)
WP 022674213.1 - Sphingopyxis baekryungensis (Xanthorhodopsin)
WP 120048384.1 - Erythrobacter aquimixticola (Xanthorhodopsin)
WP 076069562.1 - Sphingomonas montana (Xanthorhodopsin)
WP 136856450.1 - Paracoccus hibiscisoli (Xanthorhodopsin)
WP 028640910.1 - Novosphingobium acidiphilum (Xanthorhodopsin)
WP 119061141.1 - Dichotomicrobium thermohalophilum (Xanthorhodopsin)
WP 114612889.1 - Marinobacter vulgaris (Xanthorhodopsin)
WP 016353389.1 - Spiribacter salinus (Xanthorhodopsin)
WP 028386460.1 - Legionella geestiana (Xanthorhodopsin)
ADY17809.1 - Oxyrrhis marina (Xanthorhodopsin)
AIN36550.1 - Alexandrium fundyense (Xanthorhodopsin)
WP 013566350.1 - Isosphaera pallida (Xanthorhodopsin)
SS bin 106 - Polyangiales (Myxococcota) - k127 28938 6
MetaBAT.bin.10 - Polyangiales (Myxococcota) - RZV51309.1
WP 014629850.1 - Thermus thermophilus (Xanthorhodopsin)
WP 011404249.1 - Salinibacter ruber (Xanthorhodopsin)
WP 098073974.1 - Longibacter salinarum (Xanthorhodopsin)
WP 072788487.1 - Ferrithrix thermotolerans (Xanthorhodopsin)
WP 019009904.1 - Deinococcus aquatilis (Xanthorhodopsin)
PZO36820.1 - Leptolyngbya antarctica (Xanthorhodopsin)
WP 116715570.1 - Euzebya tangerina (Xanthorhodopsin)
WP 070955062.1 - Candidatus Rhodoluna planktonica (Xanthorhodopsin)
WP 148613022.1 - Nocardioides rubriscoriae (Xanthorhodopsin)
WP 085179990.1 - Mycobacterium bohemicum (Xanthorhodopsin)
WP 110125929.1 - Cryobacterium arcticum (Xanthorhodopsin)
WP 092108292.1 - Cryobacterium luteum (Xanthorhodopsin)
WP 064013919.1 - Deinococcus puniceus (Xanthorhodopsin)
SS bin 3 - UBA5794 (Actinobacteriota) - k127 330552 10
MetaBAT.bin.17 - UBA5794 (Actinobacteriota) - RZV46091.1
SS bin 7 - UBA5794 (Actinobacteriota) - k127 2471362 8
SS bin 103 - Microtrichales (Actinobacteriota) - k127 2217180 23
MetaBAT.bin.19 - Microtrichales (Actinobacteriota) - RZV44704.1
SS bin 31 - Microtrichales (Actinobacteriota) - k127 1268060 5
WP 052091190.1 - Modestobacter caceresii (Sodium rhodopsin)
WP 093408436.1 - Verrucosispora sediminis (Sodium rhodopsin)
WP 091290184.1 - Micromonospora halophytica (Sodium rhodopsin)
WP 013178725.1 - Truepera radiovictrix (Sodium rhodopsin)
WP 052001698.1 - Lyngbya aestuarii (Sodium rhodopsin)
SS bin 10 - Rhodothermales (Bacteroidota) - k127 2688974 3
WP 098061420.1 - Longimonas halophila (Sodium rhodopsin)
WP 073020395.1 - Flavobacterium micromati (Sodium rhodopsin)
WP 009036080.1 - Indibacter alkaliphilus (Sodium rhodopsin)
WP 106133859.1 - Mongoliibacter ruber (Sodium rhodopsin)
WP 093365882.1 - Psychroflexus sediminis (Sodium rhodopsin)
WP 015361652.1 - Nonlabens dokdonensis (Sodium rhodopsin)
WP 132702639.1 - Winogradskyella wandonensis (Sodium rhodopsin)
WP 130285848.1 - Aquimarina brevivitae (Sodium rhodopsin)
WP 054558945.1 - Croceitalea dokdonensis (Sodium rhodopsin)
WP 092981016.1 - Robiginitalea myxolifaciens (Sodium rhodopsin)
WP 063770343.1 - Mesorhizobium sp. F7 (Proteorhodopsin)
WP 153982688.1 - Chryseoglobus frigidaquae (Proteorhodopsin)
WP 130485172.1 - Microcella putealis (Proteorhodopsin)
WP 130281748.1 - Microcella alkaliphila (Proteorhodopsin)
WP 012370306.1 - Exiguobacterium sibiricum (Proteorhodopsin)
MBS85746.1 - Candidatus Heimdallarchaeota archaeon (Proteorhodopsin)
RJV00173.1 - Candidatus Poseidoniales archaeon (Proteorhodopsin)
MAF93315.1 - Candidatus Woesearchaeota archaeon (Proteorhodopsin)
WP 099307586.1 - Zhengella mangrovi (Proteorhodopsin)
WP 113888473.1 - Roseiarcus fermentans (Proteorhodopsin)
WP 085123257.1 - Tistlia consotensis (Proteorhodopsin)
WP 075534100.1 - Candidatus Pelagibacter ubique (Proteorhodopsin)
WP 008544914.1 - Candidatus Pelagibacter (Proteorhodopsin)
WP 068902630.1 - Enterovibrio pacificus (Proteorhodopsin)
WP 009599924.1 - Vibrio caribbeanicus (Proteorhodopsin)
WP 005363827.1 - Photobacterium angustum (Proteorhodopsin)
DAC44297.1 - Candidatus Poseidoniales archaeon (Proteorhodopsin)
WP 089515542.1 - Polynucleobacter cosmopolitanus (Proteorhodopsin)
WP 139875515.1 - Candidatus Methylopumilus universalis (Proteorhodopsin)
WP 148212414.1 - Candidatus Puniceispirillum marinum (Proteorhodopsin)
WP 004741718.1 - Vibrio splendidus (Proteorhodopsin)
WP 014107434.1 - Glaciecola nitratireducens (Proteorhodopsin)
WP 006008821.1 - Glaciecola pallidula (Proteorhodopsin)
WP 091548995.1 - Thermoflexibacter ruber (Proteorhodopsin)
WP 026210844.1 - Flexithrix dorotheae (Proteorhodopsin)
SS bin 60 - Chitinophagales (Bacteroidota) - k127 3698491 14
SS bin 77 - Chitinophagales (Bacteroidota) - k127 876053 2
SS bin 96 - Chitinophagales (Bacteroidota) - k127 2895057 4
SS bin 53 - Flavobacteriales (Bacteroidota) - k127 693241 4
WP 037358742.1 - Schleiferia thermophila (Proteorhodopsin)
WP 024282111.1 - Algoriphagus marincola (Proteorhodopsin)
WP 009185529.1 - Cecembia lonarensis (Proteorhodopsin)
WP 077129686.1 - Spirosoma montaniterrae (Proteorhodopsin)
WP 019990840.1 - Rudanella lutea (Proteorhodopsin)
SS bin 62 - Flavobacteriales (Bacteroidota) - k127 482690 4
SS bin 76 - Chitinophagales (Bacteroidota) - k127 2400119 3
WP 044221904.1 - Phaeodactylibacter xiamenensis (Proteorhodopsin)
WP 019038416.1 - Psychroflexus tropicus (Proteorhodopsin)
WP 035128632.1 - Flavobacterium aquatile (Proteorhodopsin)
WP 091089659.1 - Flavobacterium gillisiae (Proteorhodopsin)
WP 091872576.1 - Pricia antarctica (Proteorhodopsin)
WP 004570062.1 - Polaribacter irgensii (Proteorhodopsin)
SS bin 108 - Flavobacteriales (Bacteroidota) - k127 527230 3
SS bin 118 - Flavobacteriales (Bacteroidota) - k127 678032 1
SS bin 120 - Flavobacteriales (Bacteroidota) - k127 96110 6
WP 042289861.1 - Nonlabens ulvanivorans (Proteorhodopsin)
WP 035325034.1 - Dokdonia donghaensis (Proteorhodopsin)
WP 053973091.1 - Polaribacter dokdonensis (Proteorhodopsin)
SS bin 56 - Flavobacteriales (Bacteroidota) - k127 3527152 7
SS bin 75 - Flavobacteriales (Bacteroidota) - k127 1507336 12
WP 088341571.1 - Robiginitalea sediminis (Proteorhodopsin)
WP 055397113.1 - Flagellimonas eckloniae (Proteorhodopsin)
SS bin 105 - Flavobacteriales (Bacteroidota) - k127 119987 8
0.2

**Figure S24.** Maximum-likelihood tree of amino acid sequences of acetyl-CoA synthase B subunit (AcsB), a marker gene for acetate oxidation. The tree shows sequences from permeable sediment metagenome-assembled genomes (blue) alongside representative reference sequences (black). The tree was constructed using the JTT matrix-based model, used all sites, and was bootstrapped with 50 replicates and midpoint-rooted. Note that this enzyme is a marker for both homoacetogenesis (reductive Wood-Ljungdahl pathway) and acetate oxidation (Wood-Ljungdahl pathway), but generally acts in the oxidative direction in sulfate-reducing bacteria.

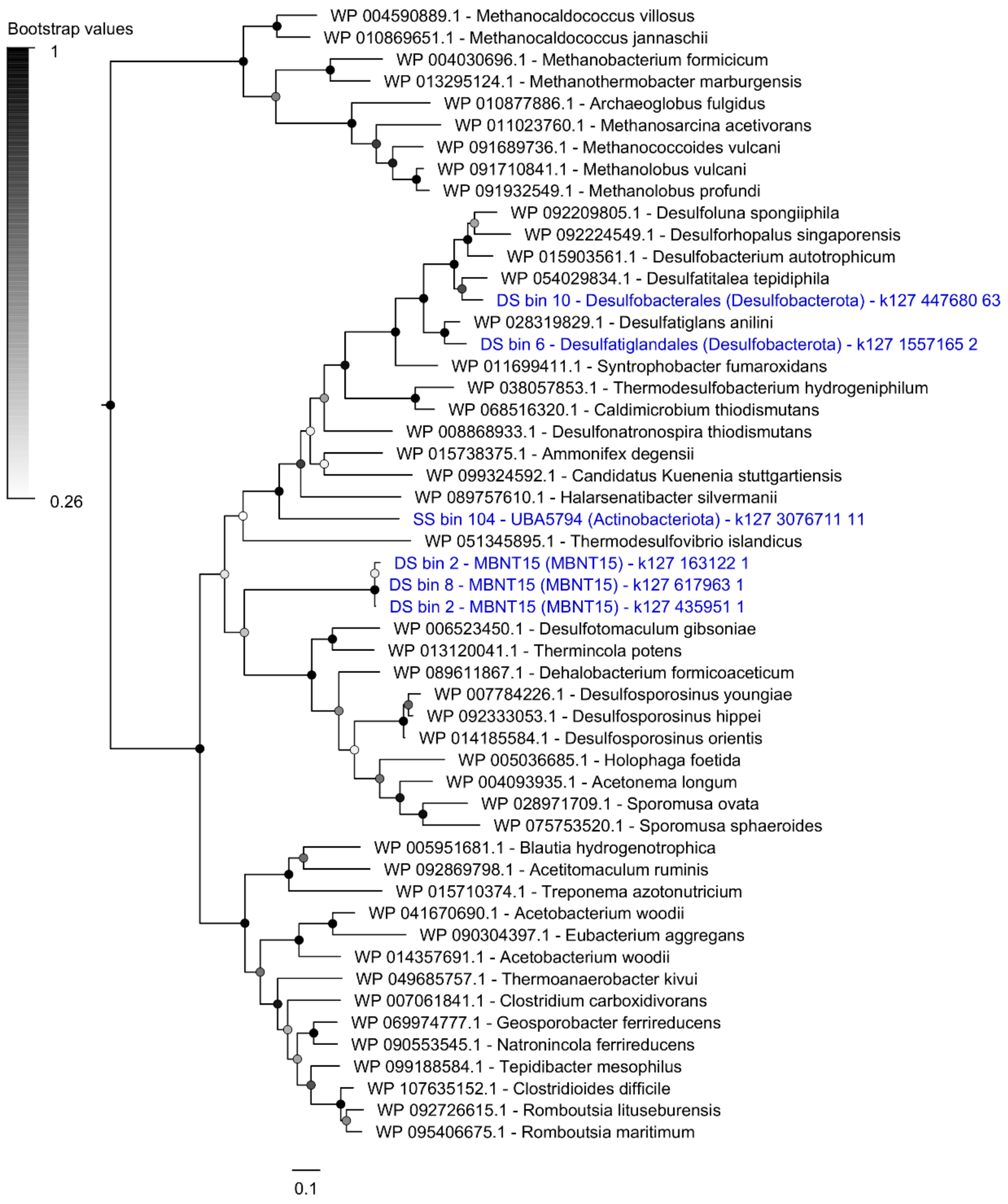

**Table S1 (xlsx).** Counts of the amplicon sequence variants detected in the 48 samples by 16S rRNA gene amplicon sequencing.

**Table S2.** Statistical testing of differences in community composition of the samples. PERMANOVA and PERMDISP were used to test significant differences of beta diversity in date and depth.

| Factor | Analysis | F. Model | $R^2$ | Pr(>F) | pvalBon | pvalFDR |
|---|---|---|---|---|---|---|
| Date | PERMANOVA | 1.2782 | 0.084 | 0.024 | | |
| | PERMDISP | 1.7319 | | 0.195 | | |
| Depth | PERMANOVA | 1.3669 | 0.294 | 0.001 | | |
| | PERMDISP | 0.7586 | | 0.627 | | |
| | Pairwise PERMANOVA | | | | | |
| | 0-15 | | 0.037 | 0.961 | 2.883 | 0.961 |
| | 0-30 | | 0.076 | 0.008 | 0.024 | 0.024 |
| | 15-30 | | 0.086 | 0.020 | 0.060 | 0.030 |

Number of permutations: 999

**Table S3 (xlsx).** Information on metagenomes analyzed for this study.

**Table S4 (xlsx).** Community composition of the samples analysed by shotgun metagenome sequencing based on the single-copy ribosomal marker gene *rplP*.

**Table S5 (xlsx).** Distribution of metabolic genes in metagenome short reads based on homology-based searches.

**Table S6 (xlsx).** Taxonomic information and metabolic capabilities of the 147 metagenome-assembled genomes analyzed.

**Table S7 (xlsx).** Counts of the amplicon sequence variants detected during the manipulative experiment by 16S rRNA gene amplicon sequencing.
